# Weight Perturbation Learning Performs Similarly or Better than Node Perturbation on Broad Classes of Temporally Extended Tasks

**DOI:** 10.1101/2021.10.04.463055

**Authors:** Paul Züge, Christian Klos, Raoul-Martin Memmesheimer

## Abstract

Biological constraints often impose restrictions for plausible plasticity rules such as locality and reward-based rather than supervised learning. Two learning rules that comply with these restrictions are weight (WP) and node (NP) perturbation. NP is often used in learning studies, in particular as a benchmark; it is considered to be superior to WP and more likely neurobiologically realized, as the number of weights and therefore their perturbation dimension typically massively exceeds the number of nodes. Here we show that this conclusion no longer holds when we take two biologically relevant properties into account: First, tasks extend in time. This increases the perturbation dimension of NP but not WP. Second, tasks are low dimensional, with many weight configurations providing solutions. We analytically delineate regimes where these properties let WP perform as well as or better than NP. Furthermore we find that the changes in weight space directions that are irrelevant for the task differ qualitatively between WP and NP and that only in WP gathering batches of subtasks in a trial decreases the number of trials required. This may allow to experimentally distinguish which of the two rules underlies a learning process. Our insights suggest new learning rules, which combine for specific task types the advantages of WP and NP. If the inputs are similarly correlated, temporally correlated perturbations improve NP. Using numerical simulations, we generalize the results to networks with various architectures solving biologically relevant and standard network learning tasks. Our findings, together with WP’s practicability suggest WP as a useful benchmark and plausible model for learning in the brain.

---

Different, usually combined strategies underlie the learning of tasks in humans and other animals [1, 2]. Supervised learning allows large, rapid improvements. It is based on observing in which way an action was erroneous and on the ability of the nervous system to use this information for the improvement of neuronal dynamics in a directed manner. This may be implemented by translating an error vector into a vector of suitable synaptic weight updates [3]. Fast learning could be achieved by directly adapting the dynamics [4]. Reward-based learning (reinforcement learning), in contrast, uses only a scalar feedback signal. It is thus also applicable if errors are known with little specificity, for example because there is sparse, delayed feedback about the cumulative effect of actions, which might only tell whether an action was erroneous but not how the generating neural activity can be improved.

A variety of models for reward-based learning have been developed in the context of theoretical neuroscience and machine learning [5, 6]. Two conceptually straight-forward implementations of such learning in neural networks are weight perturbation (WP) [7, 8] and node perturbation (NP) [9, 10]. Their underlying idea is to add perturbations to the weights or to the summed weighted inputs and to correlate them to the change of task performance. If the reward increases due to an attempted perturbation, the weights or the node dynamics are changed in its direction. If the reward decreases, the changes are chosen oppositely. NP and WP are used to model reward-based learning in biological neural networks, due to four properties [4, 7, 9–14: (i) They are (with minor modifications) biologically plausible. (ii) They are applicable to a broad variety of networks and tasks. (iii) They are accessible to analytical exploration. (iv) They are optimal in the sense that the average of the generated weight change taken over all noise realizations is along the reward gradient. The schemes’ names were originally coined for approaches that directly estimate the individual components of the gradient using single perturbations to each weight or node [15].

WP explores with its random perturbations a space with dimensionality equal to the number of weights. For trials without temporal extent, NP only needs to explore a space with dimensionality equal to the number of nodes. The chain rule, amounting to a simple multiplication with the unweighted input strength, then allows to translate a desirable change in the summed weighted inputs into a change in a particular weight strength. NP thus uses additional information on the structure of the network (namely the linearity of input summation) to reduce the required exploration.

In linear approximation the optimal direction of weight changes aligns with the direction of the gradient of the reward. WP and NP seemingly attempt to find this direction by trying out random perturbations. Since the dimension of the space of possible perturbation directions is large, the probability of finding the gradient is small and a lot of exploration is necessary. This impedes WP and NP. The number of weights and thus the dimensionality of the perturbation space searched by WP is much larger than the number of nodes. NP is thus considered more suitable for reward-based neural network learning [3, 9, 10, 16] and its implementation in biological neural networks [17]. This is supported by quantitative analysis: ref. [9] considered *M* linear perceptrons with *N* random inputs, using a student-teacher task. They found that for WP the optimal convergence rate of the student to the teacher weight matrix is by a factor *NM* worse than for exact gradient descent (GD). This is consistent with the argument that WP needs to search the *NM*-dimensional weight space to find the gradient, which is directly computed by GD. Accordingly, NP is worse than gradient descent by the dimensionality *M* of the node perturbation space.

The prerequisites of the arguments sketched above, however, do not hold in many biological situations. First, tasks in biology often extend in time and have a reward feedback that is temporally distant from the action [1, 5, 10, 14, 18]. Second, the effective dimension of neural trajectories and of learning tasks is often comparably low [19–21]. Our article analytically and numerically explores the perturbation-based learning of tasks with these features.

The article is structured as follows. First, we introduce the employed WP and NP learning models. We then derive analytic expressions for the evolution of expected error (negative reward) in linear networks solving temporally extended, low-dimensional reward-based learning tasks. This allows to identify conditions for which WP outperforms NP as well as the underlying reasons. Furthermore we delineate distinguishing qualitative characteristics of the weight and error dynamics. Finally we numerically show that WP is comparably good or outperforms NP in different biologically relevant and standard network learning applications.

## RESULTS

### Learning models and task principles

Our study models the learning of tasks that are temporally extended. Time is split into discrete steps, indexed by *t* = 1,…, *T*, where *T* is the duration of a trial. During this period, a neural network receives external input and generates output. At the end of a trial, it receives a scalar error feedback *E* about its performance [8, 10, 14, 17, 22]. To quantitatively introduce the learning rules, we consider a neuron *i*, which may be part of a larger network. It generates in the *t*-th time bin an output firing rate *z_it_*, in response to the firing rates *r_jt_* of its *N* presynaptic neurons,

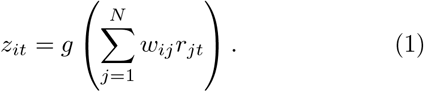

Here *w_ij_* is the weight of the synapse from neuron *j* to neuron *i*. The generally nonlinear activation function *g* implements the relation between the total input current and the output firing rate of the neuron [5, 23]. We note that the individual synaptic input currents *w_ij_r_jt_* in the model sum up linearly. This is a standard assumption, and it is a requirement for the NP scheme [9, 10, 15, 24]. In presence of nonlinear dendritic compartments [25–27], each of these could be an independently perturbed node.

We model WP learning by adding in the beginning of a trial a temporally static weight change 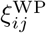 to each of the weights *w_ij_* [8, 22]. The output of the neuron then reads

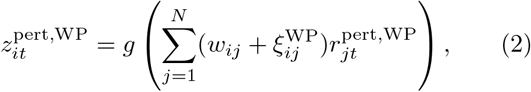

where 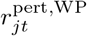 are the input rates, which may have a perturbation due to upstream perturbed weights. 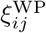 are independent and identically distributed (iid) zero-mean Gaussian white noise perturbations with standard deviation 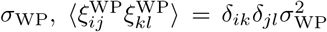, where the angular brackets denote the average over perturbation realizations and *δ* is the Kronecker delta. The perturbations 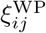 change the output, which in turn influences the reward received at the end of the trial, Fig. 1a) left hand side. We usually assume that the difference in reward between the perturbed and an unperturbed trial with 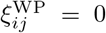 for all *i, j* is used to estimate the gradient: When the reward increases, for small perturbations this means that the tried perturbation has a positive component along the reward gradient. Consequently the update is chosen in the direction of that perturbation. When the reward decreases, the update is chosen in the opposite direction. We use the update rule

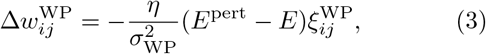

where *η* is the learning rate, *E*^pert^ is the error of the perturbed trial and *E* is the error of the unperturbed trial. For the delayed non-match-to-sample (DNMS) task, the error of the unperturbed trial is replaced by an average over the previous errors for biological plausibility. The proportionality of update size and obtained reward implies that when averaging over the distribution of the perturbations, the weight change

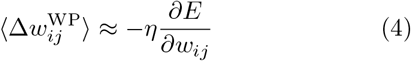

is parallel to the reward gradient (Supplementary Material (SM) 1, Eq. (S2) [28]). Since 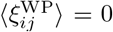, this holds for any baseline in Eq. (3). The employed choice of baseline *E* guarantees that for small perturbations the weight change has a positive component in the direction of the reward gradient and thus always reduces the error for sufficiently small learning rate *η* [8]. In fact it minimizes the update noise, i.e. the fluctuations of updates around the gradient (SM 1, Eq. (S5)).

**Figure 1.**
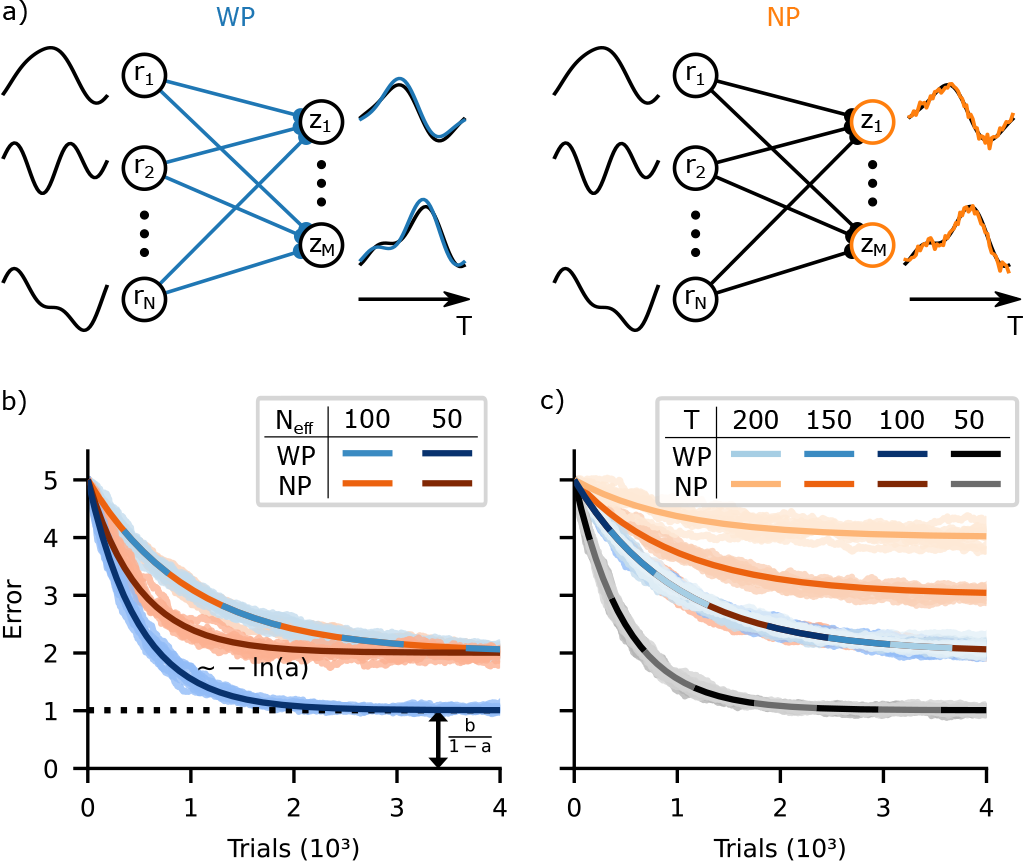
Learning of temporally extended tasks in linear networks. a) Schematic setup of WP and NP. The *M* outputs *z_i_* are weighted sums of the *N* inputs *r_j_*. Left: WP perturbs the weights at the beginning of a trial; the resulting perturbations of the weighted sums of the inputs and thus the outputs reflect the dimensionality and smoothness of the inputs (blue). Right: NP perturbs the weighted sums of the inputs with dynamical noise (orange). b) WP (blue) works just as well as or better than NP (orange) when learning a single temporally extended input-output mapping. The error decay time decreases for WP and NP likewise with decreasing effective input dimension *N*_eff_ (light vs. dark curves). In contrast, the residual error only decreases for WP. c) Increased trial duration *T* does not change the progress of WP learning (blue curves lie on top of each other). In contrast, increasing *T* hinders NP learning by increasing the residual error (compare the increasingly lighter orange curves for larger *T*). If *T* decreases *N*_eff_ (gray curves), convergence is faster and to a lower residual error in both WP (because of the decrease in *N*_eff_) and NP (because of the decrease in *N*_eff_ and *T*). Panel (b) shows error curves from simulations (10 runs, shaded) together with analytical curves for the decay of the expected error (solid), for fixed *T, N* = 100, *M* = 10 and *N*_eff_ ∈ {100, 50}; *σ*_eff_ = 0.04. Theoretical curves and simulations agree well. For WP and *N*_eff_ = 50 the decay rate (−ln(*a*)) and the residual error (dashed line) are highlighted. Panel (c) shows error curves from simulations and theory similar to (b) for fixed *N* = 100 and *T* ∈ {200, 150, 100, 50}. *N*_eff_ is set to 100 but cannot be greater than *T*, such that *T* = 50 forces *N*_eff_ = 50.

WP treats the system as a black box, mapping parameters *w* onto a scalar error function *E*. In other words, it uses the information that the weights are fixed parameters during a trial, but does not take advantage of specifics of the network structure. This is in contrast to NP, which takes into account some minimal structural knowledge, namely the linear summation of input currents, but not the constancy of the weights.

Instead of perturbing the weights directly, NP adds noise to the sum of the inputs,

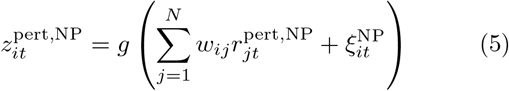

[9, 10], see Fig. 1a) right hand side. 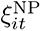 are iid zero-mean Gaussian white noise perturbations with standard deviation 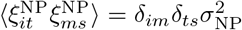. We note that for temporally extended tasks, in contrast to WP the noise must be time dependent to explore the space of time-dependent sums of inputs [10]. For temporally constant noise, only the temporal mean of the total input would be varied and thus improved.

The NP update rule can be defined as

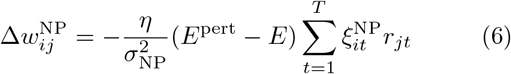

[10], with the eligibility trace 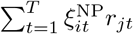. As for WP, this yields an average weight update parallel to the reward gradient, which again holds for any baseline of the weight update. The choice of baseline *E* again minimizes the update noise (SM 1, Eq. (S6)).

The NP update rule effectively incorporates an error backpropagation step, which reduces to a simple multiplication with *r_jt_* due to the linearity of the spatial synaptic input summation. This allows to perturb only summed inputs instead of individual weights and may be expected to increase the performance of NP compared to WP [3, 10, 14, 17].

### Theoretical analysis

We analytically compare WP and NP for temporally extended tasks by training a set of *M* linear perceptrons with *N* inputs. The task is to learn the mapping of a single fixed input sequence of duration *T* to a target output sequence in reward-based manner. The task choice is motivated firstly by biological motor tasks that require such a mapping, like the learning of their single song in certain songbirds (Discussion). Secondly, it yields novel insights as it is the opposite extreme case to having no time dimension and different, random inputs in each trial; this case was treated analytically by ref. [9] (Introduction). Thirdly, our findings yield an understanding of the learning performance for more general temporally extended tasks and networks studied later in this article. The analysis shows how learning depends on task dimensions and the structure of the input. Furthermore it reveals specific disadvantages of WP and NP. Importantly, our theoretical considerations cover very general sequences. In particular, they hold for sequences with and without correlations between subsequent inputs. Furthermore, the sequences can be arbitrarily reordered, also differently in each trial. They may therefore also be interpreted as sets or batches of inputs. In Sec. “Input and perturbation correlations” we consider temporally correlated sequences, for which such an interpretation is not useful anymore. In Secs. “Multiple subtasks” and “Input noise” we relax the assumption of exactly repeated input sets.

The perceptrons generate as outputs the product of their *M* × *N* weight matrix *w* with the inputs,

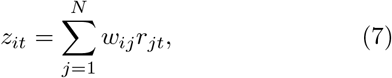

where *i* = 1,…, *M*, Eq. (1) and Fig. 1a). For now we assume that the target output can be produced with target weights *w*\*, that is 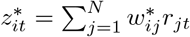. This condition will be alleviated in Sec. “Unrealizable targets”. The learning goal is to reduce the quadratic deviation of each output from its target, which can be expressed through the weight mismatch *W* = *w* − *w*\* and the input correlation matrix 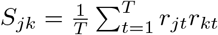 [5],

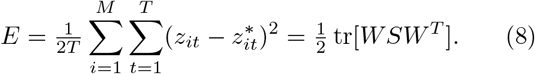

We note that with this quadratic error function the average weight update (cf. Eq. (4)) follows the gradient exactly, both for WP and NP (SM 1, Eqs. (S16,S17)).

We assume that the inputs are composed of *N*_eff_ orthogonal latent inputs; all other input components are zero (this is relaxed in Sec. “Input noise”). Since there are at most *T* linearly independent vectors of length *T*, the effective input dimension *N*_eff_ is bounded by *N*_eff_ ≤ *T*. *T* > 1 thus renders our learning problem non-trivial, by allowing for inputs that are higher dimensional when considering the input-output relation of a single sequence. In biological systems, inputs are low dimensional; *N*_eff_ is often of the order of 10 (Discussion), in particular *N*_eff_ ≪ *N*. As long as inputs are summed linearly, for clarity we will then hypothetically “rotate” the inputs such that only the first *N*_eff_ inputs are nonzero and equal to the latent ones (Fig. 2a). This allows us to speak about relevant and irrelevant inputs instead of relevant (non-zero) and irrelevant (zero) input space directions. It does not affect the WP or NP learning process, because all perturbations are isotropic and the error function is rotationally invariant (SM 3, Eq. (S71)). In other words, all results, in particular the dynamics of the error decay, hold identically for the original networks with non-rotated inputs where all actual inputs may be nonzero. For simplicity in our mathematical analysis we assume that all latent inputs have the same strength *α*^2^, i.e. 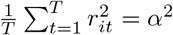 for the nonzero inputs *i* = 1, …, *N*_eff_. A partial treatment of networks with inhomogeneous latent input strength is given in SM 6.

**Figure 2.**
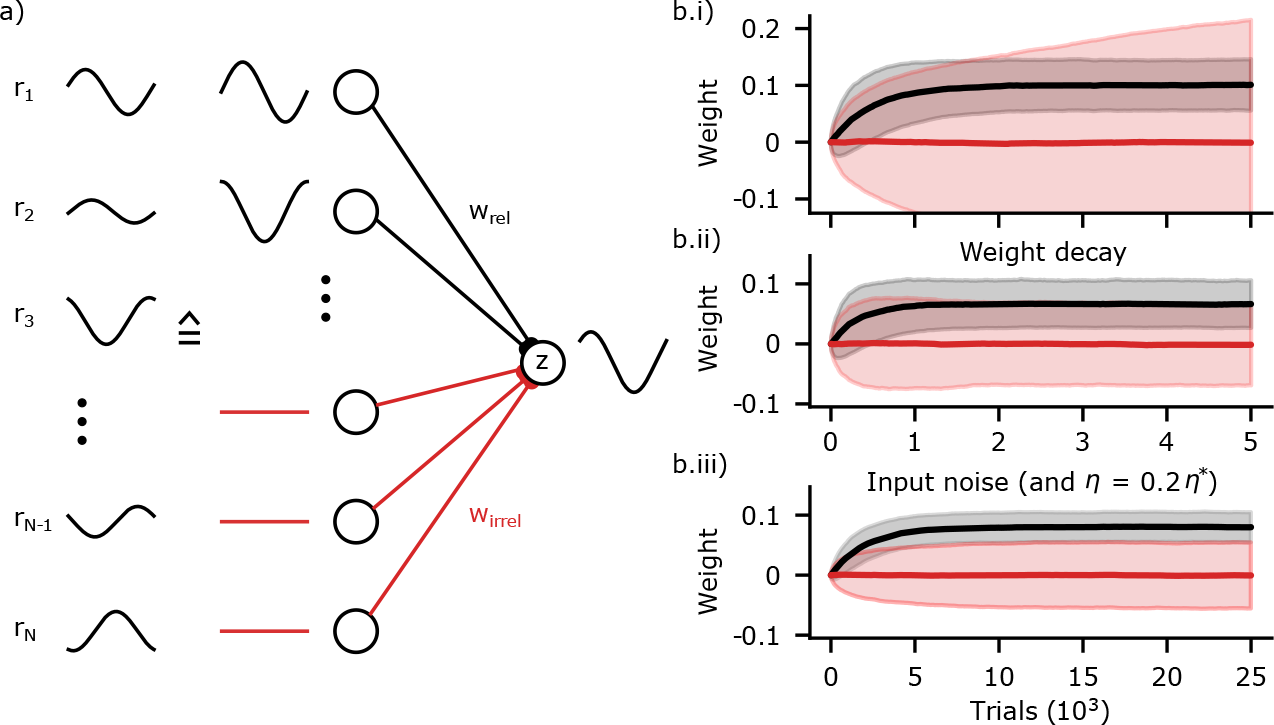
Hypothetical rotation of inputs and weight diffusion. a) Because the inputs (left, black) are summed linearly, they can be “rotated” so that for our tasks the first *N*_eff_ inputs are nonzero and agree with the latent inputs (middle, black). The remaining inputs are then zero (middle, red) and their weights are irrelevant for the output (right, red). b.i) In WP with finite perturbation size *σ*_WP_, the irrelevant weights diffuse without bounds (red), while the relevant weights converge and fluctuate (black) around the teacher weights. Displayed are the mean (solid) and standard deviation (shaded area) of the weight ensembles. b.ii) Weight decay or b.iii) input noise confines the diffusion. In (a), *N*_eff_ = 2, the latent inputs are a sine and a cosine. Parameters in (b.i,ii): *M* = 10, *N, T* = 100, *N*_eff_ = 50, *σ*_eff_ = 0.04, teacher weights *w*_rel,*i*_ = 0.1, weight decay *γ*_WD_ = 0.999; results are averaged over 10 runs. b.iii) Same parameters, except *η* = 0.2*η*\* and added iid white input noise with strength 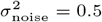 (SNR=2).

### Error dynamics

To elucidate the learning process and its dependence on the network and task parameters, we analytically derive the evolution of the expected error. This requires the computation of the error signal *E*^pert^ − *E* and weight update after a given perturbation to determine the new error. Subsequent averaging over all perturbations yields the expected error at trial *n*, 〈*E*(*n*)〉, as a function of 〈*E*(*n* − 1)〉, specifically a linear recurrence relation

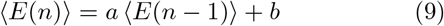

(see SM 2 [28] for the detailed derivation). The speed of learning is determined by the convergence factor *a*, while the per-update error increase *b* limits the final performance. Learning will stop at a finite error when an equilibrium between gradient-related improvement and reward noise-induced deterioration is reached. The recurrence relation is solved by

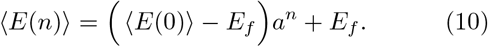

For *a* < 1, the average error 〈*E*(*n*)〉 converges exponentially at a rate −ln(*a*) towards a finite final (residual) error of 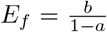, as shown in Fig. 1b). Usually in our settings *a* is sufficiently close to 1 to well approximate the convergence rate by −ln(*a*) ≈ 1 − *a*.

To understand how learning depends on the task parameters, we first consider the speed of learning. The determining convergence factor,

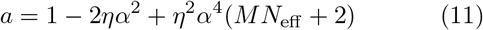

(SM 2, Eq. (S62)), is affected by two opposing effects: On average, updates follow the gradient, thus reducing the error. This is reflected by a reduction of *a* by −2*ηα*^2^ (+*η*^2^*α*^4^), responsible for convergence. However, updates fluctuate, adding a diffusive part to the weight evolution which slows convergence down. Although these fluctuations, having zero mean, do not influence the expected error to linear order, they do so quadratically. Thus their contribution to *a*, *η*^2^*α*^4^(*MN*_eff_ + 1), is quadratic in the learning rate *η*. It is approximately proportional to *MN*_eff_, the number of relevant weights that read out from nonzero inputs: fluctuations in each of these weights yield the same contribution - the exception being the twice as strong fluctuations along the single gradient-parallel direction, which together with the quadratic effect of the mean update cause the +2 in Eq. (11).

The fluctuations originate from a credit assignment problem: Only the perturbation parallel to the error gradient can be credited for causing the linear part of the error signal *E*^pert^ − *E*. WP has no way of directly solving the credit assignment problem of identifying this direction. Thus the perturbations of all *MN* weights are equally amplified in the constructions of their updates, Eq. (3), such that all weights fluctuate. This entails fluctuations in the *MN*_eff_ relevant weights, which influence output and error. NP can at least partially solve the credit assignment problem by using eligibility traces, which are zero for weights that read out from zero inputs. By projecting each of its (*M*) *T*-dimensional output perturbations onto the effectively *N*_eff_-dimensional inputs, NP restricts its updates to the *MN*_eff_-dimensional subspace of relevant weights. The convergence speed thus becomes independent of *T* as for WP. Interestingly, WP and NP therefore converge at the same speed despite their different numbers of fluctuating weights. The reason is that the fluctuations of the relevant weights are the same for both algorithms.

The balance between the improvement resulting from following the gradient (~*η*) and the deterioration due to the fluctuations of relevant weights (~*η*^2^) in Eq. (11) is controlled by the learning rate: small learning rates imply averaging out fluctuations over many updates and therefore dominance of gradient following, leading to convergence. For the remainder of the analysis of this setting, both algorithms will be compared at their optimal learning rate *η*\*, which is defined to yield fastest convergence, in other words: to minimize *a*. This definition is chosen because it is conceptually straightforward and Eq. (11) directly leads to the simple expressions

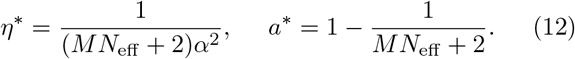

Here, the factor 1/*α*^2^ in *η*\* cancels the scaling of the gradient with the input strength and equals the optimal learning rate for GD (Eq. (S28)). In order to allow for averaging out the update fluctuations, WP and NP learning additionally have to slow down by a factor of approximately *MN*_eff_. Learning diverges for *η* → 2*η*\* where *a* → 1. Eq. (12) shows that WP’s convergence rate is worse than GD’s by a factor generally smaller than the number of weights. Further, NP’s convergence rate is worse by a factor generally larger than the number of nodes. Thus, the number of weights or nodes is insufficient to predict the performance of WP or NP, respectively.

The per-update error increase and the final error, *b* and *E_f_*, result from finite perturbation sizes. Finite perturbation sizes lead, due to the curved, quadratic error function, to an estimate that is at least slightly incompatible with the linear approximation assumed by the update rules (cf. Eq. (4)). This is particularly apparent when the output error and thus the gradient is (practically) zero: Any finite weight or node perturbation then leads to an increase of the error and thus to an opposing weight update instead of no weight modification. This prevents the weights from reaching optimal values and results in a finite final error *E_f_*. The described difference between perturbation-based error estimate and linear approximation is a form of “reward noise”. It is nonzero only for finite perturbation size, as reflected by the dependence of *b* and *E_f_* on *σ* (which is quadratic due to the quadratic error nonlinearity).

For a fair comparison of WP and NP we choose *σ*_WP_ and *σ*_NP_ such that they lead to the same effective perturbation strength 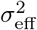, as measured by the total induced output variance. This leads to 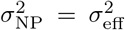 and 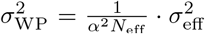 (SM 1, Eq. (S22)). Evaluated at the optimal learning rate *η*\*, the leading order term of the final error is

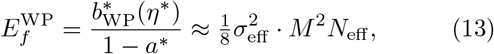

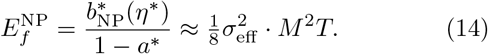

Importantly, the final error of WP is here generally smaller, by a factor *N*_eff_ /*T* ≤ 1. To understand this, we focus for both WP and NP on the output perturbations that they generate. By perturbing the weights, WP induces output perturbations that are linear combinations of the inputs. These are confined to the effectively *MN*_eff_-dimensional subspace in which also the (realizable part of the) output error gradient (*z* − *z*\*)/*T* lies. NP, on the other hand, creates an entirely random *MT*-dimensional perturbation vector (Fig. 1a). Only the projection of this vector onto the output gradient is useful for learning. This projection is smaller for NP’s random vector, since the vector has effectively a larger dimensionality than the output perturbation vector of WP, at the same length. NP compensates this deficit by amplifying the smaller gradient projection more strongly. It thus achieves the same mean update and convergence speed as WP. However, it also more strongly amplifies the reward noise that comes with larger perturbation sizes, which results in a larger final error. The scaling of *E_f_* with *M* ^2^*N*_eff_ or *M* ^2^*T* reflects the effective output perturbation dimensions, *MN*_eff_ or *MT* of WP or NP, and additionally the general scaling of errors with *M* (Eq. (8), SM 3, Eqs. (S126,S127)).

Taken together, we observe that here WP learning works just as well as or better than NP. Both algorithms have the same speed of convergence, but the final error *E_f_* for WP is smaller or equal compared to NP. The rate of convergence decreases with increasing *M* and *N*_eff_. Longer trial durations *T* harm NP by linearly increasing *E_f_*. Larger effective input dimensionality *N*_eff_ similarly harms WP. This result differs from the observation in ref. [9] that WP converges much (*N*-times) slower than NP. The reason is that our networks learn a single temporally extended input-output relation, while those in [9] learn the weights of a teacher network, by trials with random input of duration *T* = 1. We will explore the relation between the results in detail in Sec. “Comparison with ref. [9]”.

### Weight diffusion

When the input has less than maximal dimensionality, *N*_eff_ < *N*, only certain combinations of weights read out nonzero components of the input. This becomes particularly clear for the considered rotated inputs: If WP adds a perturbation to a weight mediating zero input, to an irrelevant weight, the output and the error remain unchanged. This missing feedback leads to an unbounded diffusion-like random walk of irrelevant weights. For unrotated inputs, the weight strength diffuses in irrelevant weight space directions. We will see below (Sec. “Multiple subtasks”) that the weight diffusion harms performance when learning multiple input-output patterns.

We find that for WP in the limit of infinitesimally small perturbations *σ*_WP_ → 0, all weights initially change and then converge (SM, Fig. S1 [28]). This is because the learning-induced drift of relevant and the diffusion of irrelevant weights both stop when the error converges to zero: the error *E* is quadratic, such that for infinitesimally small perturbations *E*^pert^ = *E* at its minimum. In contrast, for finite perturbations a residual error remains and weights continue to change. In particular, irrelevant weights continue to diffuse, Fig. 2b.i. The quantitative details of the weight diffusion process can be analytically understood (SM 4, Eqs. (S131,S134)). Standard mechanisms such as an exponential weight decay [29, 30] confine their growth. Simultaneously they bias the relevant weights towards zero and therewith increase the residual error, Fig. 2b.ii. Also input noise confines the weight spread, by adding error feedback to irrelevant weights. Simultaneously the noise increases the final task performance error and enforces a lower learning rate, see Fig. 2b.iii, Sec. “Input noise” and Fig. 5a,b.

NP does not generate weight diffusion: the rotated inputs make it obvious that in NP the eligibility trace (Eq. (6)) selects only the weights from relevant inputs to be updated, since for irrelevant inputs we have *r_jt_* = 0 for all *t* such that 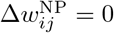. Input noise renders irrelevant inputs and their weight updates nonzero, such that irrelevant weights spread also for NP.

Differences in weight spread and updates between WP and NP suggest experimental measurements to distinguish which one of them underlies learning of a certain task: As long as the noise is weak compared to the signal such that the task can be satisfied with high precision, the spread is with NP much smaller than with WP, Fig. 5b. Further, a large variance in the weight updates that is independent of presynaptic activity together with a resulting weight spread that is largest for weight directions that read out weak latent inputs point to WP (Fig. 5b and SM 6, Eq. (S196)). Consistent with this, prominent random walk-like weight changes that are unrelated to neuronal activity and task learning are common in biological neural networks [31]. Weight updates whose variance scales with input strength but whose final spread is independent of it (Fig. 5b and SM 6, Eq. (S198)) point to NP.

### Unrealizable targets

In the previous sections we assumed that the target outputs 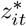 could be exactly realized by setting the perceptron weights *w* equal to some target weights *w*\*, 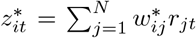. In general, however, the target outputs may contain components *d* that cannot be generated by the network, which is limited to producing linear combinations of the inputs. Unrealizable components are orthogonal to all inputs when interpreted as *T*-dimensional vectors, 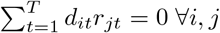. The target may be written as a sum of realizable and orthogonal unrealizable parts, 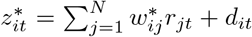. An illustration of such a target is given in Fig. 3a. In practice, unrealizable targets occur for example in machine learning classification tasks, see Sec. “MNIST”.

**Figure 3.**
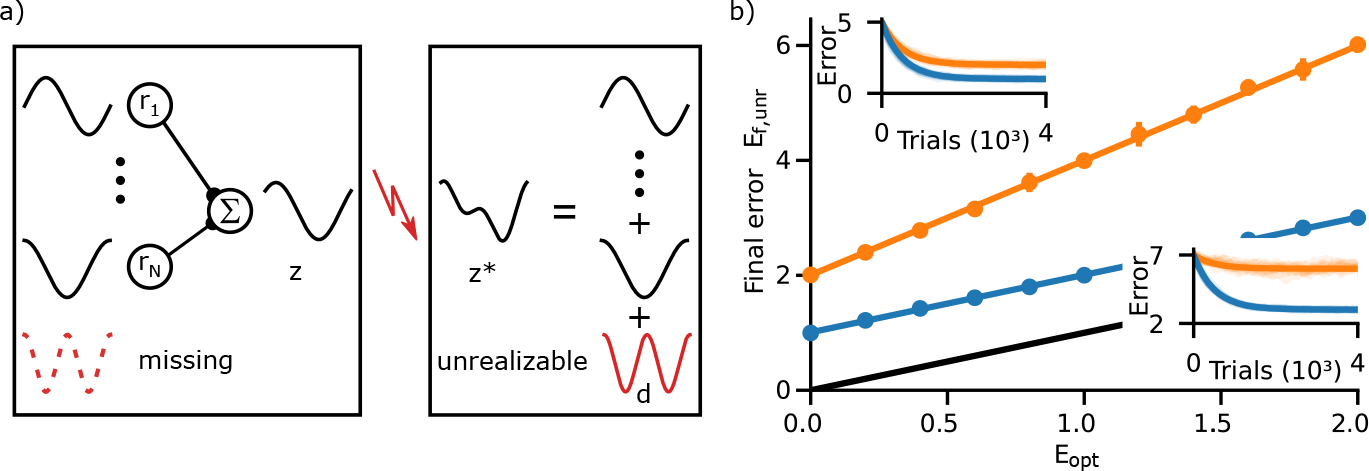
Unrealizable target components harm NP learning. a) General targets may contain a component that is perpendicular to any input and thus unrealizable (red). b) Final error after convergence as a function of the error *E*_opt_ that necessarily remains, since the target is unrealizable. The final error of WP (blue) is only shifted by *E*_opt_, that of NP (orange) increases twice as fast, by approximately 2*E*_opt_. Data points: mean and standard deviation (averaged over 10 simulated runs) of the final error. Curves: theoretical predictions. Black: *E*_opt_. Insets: error dynamics for *E*_opt_ = 0 (left) and *E*_opt_ = 2 (right). Parameters: *M* = 10, *N, T* = 100, *N*_eff_ = 50, *σ*_eff_ = 0.04.

WP induces output perturbations 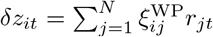, which are linear combinations of the inputs. The components of *z*^pert^ − *z*\* that are orthogonal to all inputs, *d*, thus always remain unchanged, irrespective of the current student weights and applied perturbations. This leads to the same constant additive contribution 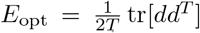 to the perturbed and unperturbed errors *E*^pert^ and *E* (Eq. (8)). It cancels in the weight update rule (Eq. (3)) such that WP learning is unchanged and Eq. (10) still holds when shifting its final error to 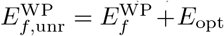 (Eq. (S119)). *E*_opt_ marks the minimum error that necessarily remains even with *w* = *w*\*, due to the unrealizable components.

In contrast, NP perturbs the outputs with white noise. This noise generally has a nonzero component along *d*, which affects *E*_pert_. Since such a component cannot be realized through an update of the weights, the resulting change of the error is non-instructive and represents reward noise that adds noise to the updates. Consequently, while the convergence factor *a* remains unchanged, the final error of NP increases more strongly than for WP, Fig. 3b. At the optimal learning rate the increase in final error due to unrealizable target components is twice that of WP, 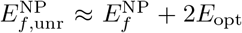 (SM 3, Eq. (S128)). For 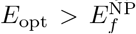 the coupling of node perturbations to unrealizable target components becomes NP’s main contribution to the part of the final error that exceeds the unavoidable *E*_opt_.

### Input and perturbation correlations

Our results hold for very general sequences. In particular correlations in the input may be present or absent without affecting learning. Further, the *T* input-output relations can be temporally permuted. These invariances follow straightforwardly from the weight update equations (SM 7).

WP generates output perturbations that are automatically adapted to the inputs (Sec. “Unrealizable targets”). To adapt NP to tasks with temporal input correlations, we modify it to time-correlated NP, NPc. We generate the correlated perturbations by temporal lowpass filtering of white noise, with filtering time constant 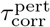. Other possibilities are to compose them from low frequency Fourier modes or to use perturbations that are piecewise constant. As the correlation of the perturbation decays during 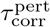, for a reasonable representation of the perturbation we need to specify it at 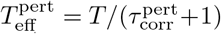 timepoints with temporal distance 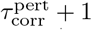. For vanishingly short correlations, 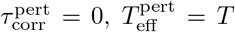 and NPc equals NP; a large filtering time constant generates perturbations that vary slowly and have small effective temporal dimension 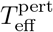.

If the inputs have similar temporal correlations, the filtering of the perturbation concentrates its perturbative power on the realizable output subspace, since this is spanned by the inputs. This reduces the update noise due to the quadratic reward noise from finite perturbation sizes (cf. Eq. (S112)), because perturbations that are more aligned with the output gradient require less amplification to yield a sizable expected update along the weight gradient. Further, it reduces the linear reward noise resulting from coupling to unrealizable target components (cf. Eqs. (S108, S80)). Both noise reductions lower the final error. We find that NPc robustly improves upon NP over a range of filtering time constants similar to that of the inputs, Fig. 4. If the perturbations become too smooth, however, they also loose power in the realizable output subspace, which slows the learning of realizable target components with higher frequencies down. This is because the suppressed modes with their comparably small amplitude contribute little to the projection of the perturbations onto the *T*-dimensional output gradients (despite being prominent in the latter), which results in a small contribution to the error signal (Eq. (S3)). This contribution in turn determines the magnitude of the mean weight update used to match an output mode to the target. Consequently the update part used to match the high frequency modes is small and the matching takes a long time. With optimal perturbation correlation time 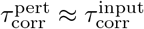 (“Materials and Methods”), NPc approaches the performance (as measured by final error and convergence speed) of NP on a substitute task with 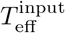 bins, see Fig. 4a,b. This indicates that our analytical considerations for NP transfer to those of NPc with optimally correlated noise if we take into account that the correlations effectively reduce the trial duration to 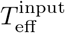. In particular, in Fig. 4 optimal NPc performs worse than WP due to the low dimensionality and the remaining temporal extension of the task as well as the unrealizable target components.

**Figure 4.**
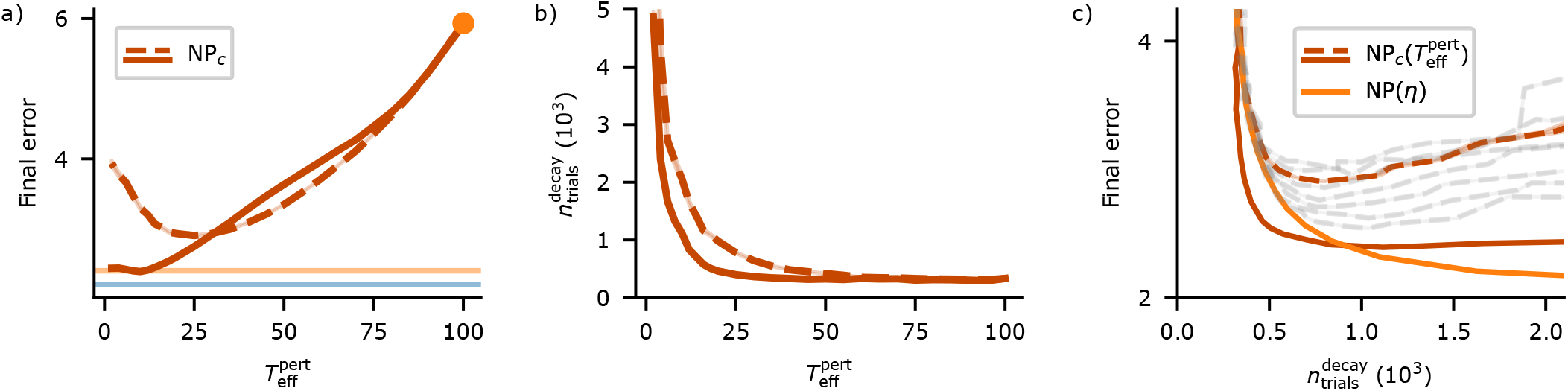
Temporally correlated perturbations improve NP if the input has similar correlations. a) Final error of NPc versus the effective time dimension 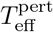 of its perturbations; smaller 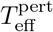 means smoother perturbations. The inputs are constructed with 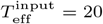 (red continuous) or 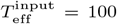 (uncorrelated, red dashed). The final error of NPc decreases compared to that of NP (orange dot) for 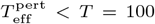. For correlated inputs and 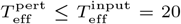, NPc’s final error reaches that of NP in a reduced task with *T* = 20 (orange line, WP: blue line); for uncorrelated inputs and small 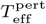 it increases again. This increase is not caused by lack of convergence. b) Convergence time, measured as number of trials until 95% of the final error correction is reached, versus 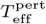. Learning gets considerably slower if the correlation times of the perturbation are longer than those of the input, i.e. 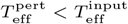. Same curve markings as in (a). c) Simultaneous plot of error and convergence time of NPc (red: data as in (a,b) for correlated (solid) or uncorrelated input (dashed), thus curves are obtained by varying 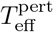; faint gray: curves for uncorrelated input with different learning rates *η*) and NP (orange, curves obtained by varying the learning rate *η*). For correlated inputs and similarly correlated perturbations, NPc yields a true improvement over NP: it has simultaneously smaller error and smaller convergence time. For considerably longer correlation times, the final error saturates, but the convergence time increases (region with 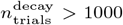). NP with reduced learning rate *η* there yields smaller final error at the same convergence time. For uncorrelated input, NPc does not yield a true improvement. In particular, the smaller final error (a) can be achieved at smaller convergence times by NP with reduced *η*, even if NPc is allowed to additionally adapt its learning rate (gray curves). Parameters: *N*_eff_ = *M* = 10, *N* = *T* = 100, *σ*_eff_ = 0.04, *E*_opt_ = 2, 20 000 trials. (a,b) Mean and standard error of the mean (SEM, partly occluded). (c) Mean values and SEM of final error (red and orange). In gray curves, *η* is modified relative to the red curve by a factor ranging from 0.5 (lowest curve) to 1.2 (highest curve) in steps of 0.1.

### Multiple subtasks

In general learning tasks, inputs and targets may vary from trial to trial. To obtain an intuition for how this affects the speed of WP and NP learning, we here consider a simplified case: The goal is to solve a task with an overall effective input dimension of 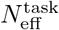. The task has the same properties as the tasks before when each trial was identical. In particular, it has 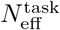 orthogonal latent inputs of strength *α*^2^ and the inputs are rotated such that only the first 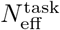 inputs are nonzero. The task is, however, not presented as a whole, but in pieces: in each trial a random subset of 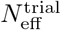 out of the first 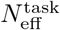 inputs are active to train the network. The error in an individual trial then only depends on its 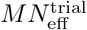 trial-relevant weights, while the performance on the full task depends on the 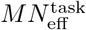 task-relevant weights. The ratio 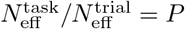 marks the number of trials needed to gather information on all task-relevant weights.

NP only updates the weights relevant in a trial (Sec. “Weight diffusion”). Also for tasks consisting of multiple subtasks it can thus operate at the learning rate that is optimal for a trial, 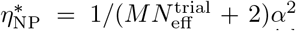 (cf. Eq. (12)). Because an update only improves 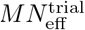 of the 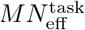 task-relevant weights, the convergence rate −ln *a* ≈ 1 − *a* of the expected error, averaged over the input distribution, is smaller by a factor of 1/*P* than for a single input pattern (SM 5, Eq. (S149)),

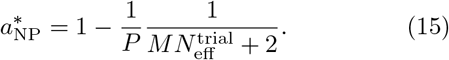

WP, on the other hand, updates all weights such that the weights that are irrelevant for the trial are changed randomly (Sec. “Weight diffusion”). This worsens the performance for the inputs of other trials. Because there are now 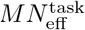 task-relevant weights whose fluctuations hinder learning, WP has an optimal learning rate of only 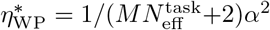. As for NP, each trial’s progress is only on 1/*P* of the task-relevant weights, such that the optimal convergence factor for WP on the full task is (SM 5, Eq. (S148))

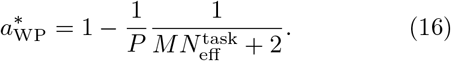

The convergence of WP is thus slower than that of NP by roughly 1/*P*, the ratio of 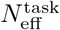 and 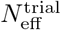, Fig. 6c.

Our results have concrete implications for learning of multiple actions such as sequences of movements [32]. They can be learned by splitting them into subsets, which are called (mini-)batches in machine learning. If we assume for simplicity that individual data points are pairwise orthogonal and have no time dimension, each batch corresponds in our terminology to a subtask, the number of batches to *P*, the dimensionality of the input data to 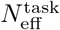 and the batch size *N*_batch_ to 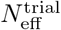. For 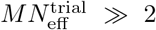, Eqs. (15,16) thus imply that the convergence rate of NP is independent of the batch size while that of WP is proportional to the batch size and reaches NP’s convergence rate for full batch learning (SM, Fig. S5). The same holds for the optimal learning rates as *α*^2^ scales inversely with the batch size (SM 5, Eq. (S160,S162,S163)).

### Comparison with ref. [9]

Ref. [9] has investigated how a student network learns the responses of a teacher network to arbitrary input with GD, NP and WP, using patterns without temporal extent. In contrast to our tasks with typically *N*_eff_ < *N* or 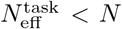, successful learning in ref. [9] requires to match all weights of the teacher network. In other words, the student network is trained at its capacity limit, where (only) one weight configuration fulfills the task. It learns from random input patterns and the teacher’s responses to them. This is a special case of the setup introduced in Sec. “Multiple subtasks”, where (i) the task dimension equals the input dimension, 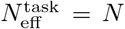 (since the employed random input patterns will lie in arbitrary directions of input space) and (ii) there is no temporal extent of the tasks, *T* = 1. The latter implies that a single input pattern has effective dimension 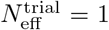: all *N* inputs in a single pattern are linearly dependent since they are scalar, temporal vectors of length 1 (we could rotate the input pattern such that it has only one nonzero entry).

A comparison of our results for the convergence speed in the described special case with those of ref. [9] reveals that they agree approximately when straightforwardly setting 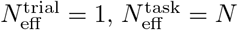 and 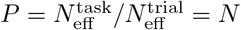 in Eqs. (15,16):

| Alg. | Our results | Results from ref. [9] |
| --- | --- | --- |
| WP | $a_{\text{WP}}^* = 1 - \frac{1}{N} \frac{1}{MN+2}$ | $a_{\text{WP}}^* = 1 - \frac{1}{N+2} \frac{1}{MN+2}$ |
| NP | $a_{\text{NP}}^* = 1 - \frac{1}{N} \frac{1}{M+2}$ | $a_{\text{NP}}^* = 1 - \frac{1}{N+2} \frac{1}{M+2}$ |

The reason for the remaining difference is that the individual inputs in ref. [9] are drawn from a Gaussian distribution, without subsequent normalization to the same strength like in our scheme. We obtain *P* → *N* + 2 and thus perfect agreement if we adapt our setting such that the summed input strength, 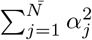, fluctuates as it does in ref. [9].

### Input noise

Inputs in biological neural networks are noisy. To investigate the impact of input noise on WP and NP, we add white noise to all input neurons. Noise in the relevant inputs will cause a finite error that remains even for optimal weights. Moreover the irrelevant weights are not completely irrelevant anymore: they mediate noise (instead of zero) input, have an optimal value of zero and lead to significant output error if they become too large. Since the input noise is different in the perturbed and unperturbed trials, it becomes a source of additional reward noise. We thus expect that larger input noise requires stronger perturbations to ensure that the beneficial, error gradient-related part of the reward signal is not dominated by reward noise (cf. also [33]). Furthermore, an increase in overall noise and additional weights that become more and more important should require the integration of more trials to extract gradient information. We therefore expect a reduction of the optimal learning rate with increasing input noise strength. Our numerical simulations confirm these points, see Fig. 5a (increase in task error and optimal error) and SM, Fig. S6 (estimation of optimal learning rates and perturbation sizes).

**Figure 5.**
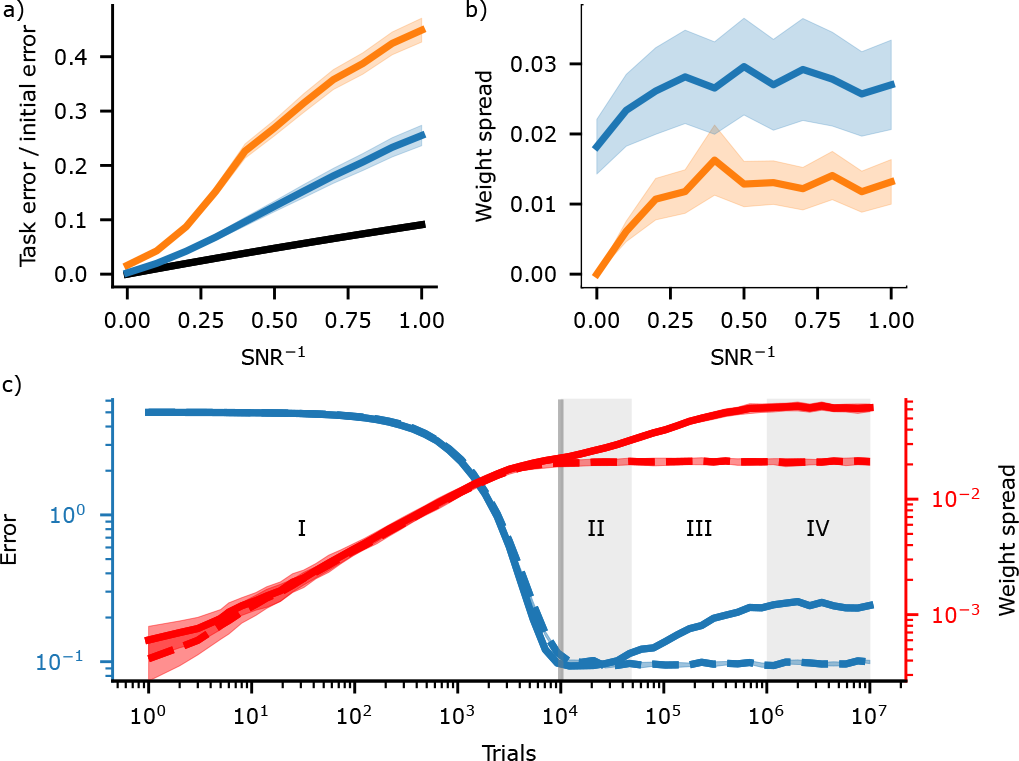
Influence of input noise on the task error and on the spread of irrelevant weights. a) Task error (fraction of the initial error) of WP (blue) and NP (orange) at 10 000 trials (curves: means, shading: standard deviations) as a function of inverse signal-to-noise-ratio (SNR^−^1). The noise in the relevant inputs renders the optimal error (black) nonzero. The plot covers SNRs ranging from infinity down to 1. b) Spread (standard deviation) of irrelevant weights (mean and standard deviation) at 10 000 trials. c) Evolution of error (blue, mean and SEM) and spread of irrelevant weights (red, mean and SEM) in WP proceeds for weak input noise (SNR^−1^ = 0.1) in four phases (solid curve, phases indicated by numerals). Appropriate weight decay stops the dynamics in phase II and induces long-term results as in the noisefree case (dashed curve, cf. also Fig. Fig. 2b.ii). (a,b) use the learning rates that minimize error after 10 000 trials (gray vertical line in (c)) for a given noise level, SM, Fig. S6. This implies that WP’s error in (a) is the one in phase II. Parameters: *M* = *N*_eff_ = 10, *N* = *T* = 100, *α*^2^ = *N/N*_eff_ = 10, *σ*_eff_ = 0.04, *E*_opt_ = 0, *γ*_WD_ = 0.99998.

We find that in presence of input noise the irrelevant weights diffuse in WP and NP (Fig. 2b.iii, Fig. 5b). They settle at a finite spread, which contributes to the error (Fig. 5c). The diffusion stops because WP and NP update irrelevant weights on average towards zero, due to their generation of errors. For WP, the final spread increases with decreasing noise strength and reaches infinity for zero noise. (Note that Fig. 5b does not show the final, stationary spread.) For NP the diffusion of irrelevant weights is caused by their noise-induced updates and is in contrast to the noise-free case. Their final spread is independent of the noise strength and discontinuously drops to zero at zero noise. To explain this we identify (weak) noise input with (weak) deterministic input and apply our findings for noise-free networks with inhomogeneous input strength distribution: For WP, each weight contributes equally to the final error and the final spread scales like one over the square root of the input strength (SM 6, Eq. (S196)). For NP, weights related to small inputs contribute only little to the error, and the final spread is independent of input strength (SM 6, Eq. (S198)).

The simulations (Fig. 5a) and our analytical understanding also show that for finite learning time or when introducing a weight limiting mechanism there is no discontinuity in the error when increasing the input noise from zero to some finite value. The previous error analysis therefore stays valid as the limit of weak input noise. For WP this is because limiting the learning time or the weights limits the final spread of irrelevant weights. This happens in such a way that sufficiently small noise has only a negligible effect on the output (SM 6, Sec. “Small input components”). For NP and weak input noise, the final spread of irrelevant weights is approximately equal to that of the relevant weights and thus also limited.

Finally, we observe that WP learning proceeds for weak input noise in four phases, Fig. 5c. In the first phase, the relevant weights are learned, such that the error decreases. Since the noise is small, the error due to its presence in relevant inputs is small. In the second phase the error remains approximately constant, at a low level. In the third phase, the irrelevant weights, which have been diffusing all the time, become so large that they amplify the input noise sufficiently to influence the output despite the small noise amplitude. The error therefore increases. In the fourth and final phase, this error becomes so large that WP counteracts the further diffusion of weights. The error therefore saturates, at a high level. The four phases can be clearly temporally separated. (Note the logarithmic axis scale chosen in Fig. 5c.) We note that we observe divergence of the error for sufficiently large irrelevant weights if the learning rate is too large or the perturbations are too weak (SM, Fig. S6).

If learning stops in the second phase, the the contributions from irrelevant weights can be neglected. The same holds if a limiting mechanism stops the diffusion of irrelevant weights at the level that they reach in the second phase, while only mildly affecting the relevant weights, because they converge at a shorter timescale (Fig. 5c, dashed line: network with weight decay, cf. also Fig. 2b.ii). Interestingly, WP can then reach a lower (final) error than NP, and is less affected by input noise, cf. Fig. 5a.

### Conclusions from the theoretical analysis and new learning rules

Our theoretical analysis reveals a simple reason for the differences between WP and NP: WP produces better perturbations, while NP better solves the credit assignment problem. Output perturbations caused by WP lie, in contrast to NP, always in the realizable output subspace and do not interfere with unrealizable target components. On the other hand, NP updates only (trial-)relevant weights, while WP updates all weights such that the (trial-)irrelevant weights change randomly. Small input noise does not change the overall picture. When single trials capture only a small part of the full task, WP learning slows down. Training in batches reduces the disadvantage.

Based on these insights, we introduce two novel learning rules, WP0 and Hybrid Perturbation (HP), Fig. 6a,b.

**Figure 6.**
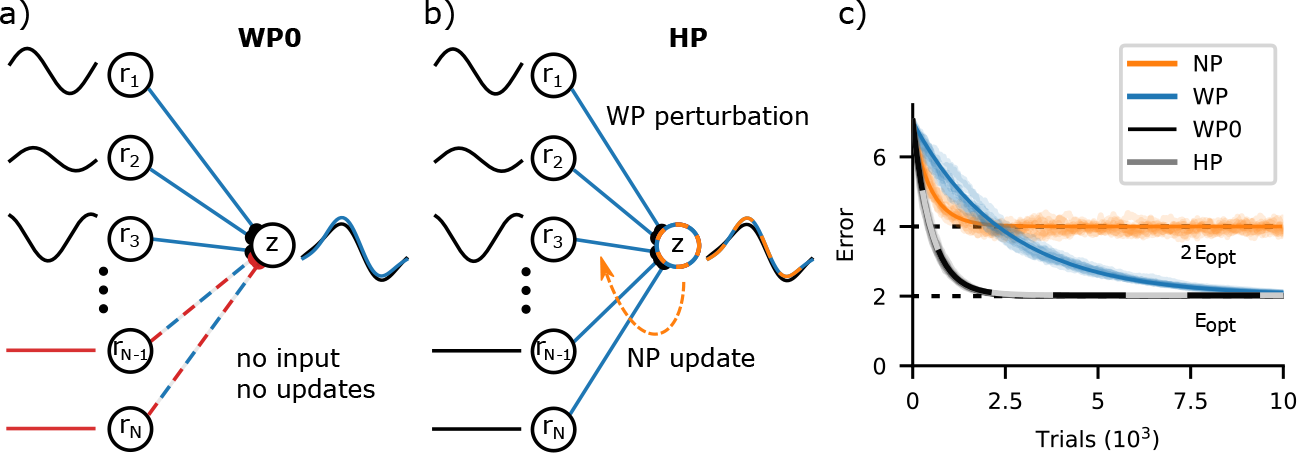
New learning rules and learning of tasks consisting of multiple subtasks. a) WP0 does not update weights that mediate zero input, avoiding their diffusion. b) Hybrid perturbation (HP): NP scheme with output perturbations induced by WP. c) WP converges approximately *P* = 5 (number of subtasks) times slower than NP, but in presence of unrealizable target components (or for finite *σ*_eff_ and 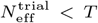, SM 5, Eqs. (S157-S159)) to a lower final error. For the used maximally sparse and equally strong inputs, WP0 and HP combine the higher convergence rate of NP with the low final error of WP. Error curves (solid: theoretical predictions, shaded: 10 exemplary runs) are for *M* = 10, *N, T* = 100, 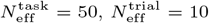, negligible *σ*_eff_ and *E*_opt_ = 2.

WP0 adds a simple modification to WP: not to update currently irrelevant weights, i.e. weights whose inputs are zero (or close to it). This solves part of WP’s credit assignment problem, as changing the weights will not improve the output, and it avoids diffusion of irrelevant weights. The improvement is especially large when inputs are sparse such that many inputs are (close to) zero (Fig. 6a,c), which might be frequently the case in biological neural networks [34–36]. HP aims to combine the advantages of WP and NP by generating the output perturbations like WP, through perturbing the weights, and generating updates like NP, using its eligibility trace. The learning rule performs well when all latent inputs have (approximately) the same strength *α*^2^, Fig. 6b,c.

WP0 and HP perform for the tasks used in the theoretical analysis section as well as WP and NP, or better than both (Fig. 6c). WP0 is, however, benefited by the assumption of rotated inputs (in contrast to WP and NP), as it renders the input maximally sparse. Further, the latent inputs have equal strengths, benefiting HP. We observe only slight improvements of WP0 over WP for the reservoir computing and MNIST task, due to the lack of coding sparseness in our networks. HP performed much worse than WP and NP in the reservoir computing and similar to NP in the MNIST task. We explain this by the relevance of weak inputs (SM 8, Eq. (S217)). Adding appropriately equalizing preprocessing layers may mitigate HP’s problems. Further, weak inputs may be irrelevant for biological learning. Since HP generates perturbations of the same class as the inputs and suppresses the learning of weights related to small inputs, we expect it to also work well for correlated input (as in Fig. 4) and in presence of input noise.

### Simulated learning experiments

In the following, we apply WP and NP to more general networks and temporally extended tasks with nonlinearities. We cover reservoir computing for dynamical pattern generation, learning of recurrent weights in a delayed non-match-to-sample task and a temporally extended, reward-based learning version of MNIST. The results confirm and extend our findings for analytically tractable tasks: they often show similar or superior performance of WP in temporally extended tasks relevant for biology and machine learning.

### Reservoir computing-based drawing task

In reservoir computing schemes an often low-dimensional input is given to a recurrent nonlinear network. The network effectively acts as a nonlinear filter bench: it expands the input and its recent past by applying a set of nonlinear functions to them. Each unit outputs one such function, which depends on the interactions within the recurrent network. Like a “computational reservoir”, the network thereby provides in its current state the results of manifold nonlinear computations on the current and past inputs. A desired result can be extracted by training a simple, often linear readout of the reservoir neurons’ activities. Reservoir computing schemes are widely used as models of neurobiological computations [37– since learning in them is simpler and seems more easily achievable with biological machinery than learning of full recurrent and multilayer networks. Further, the schemes explain the presence of inhomogeneity and apparent randomness in neuron properties and connectivity in biological neural networks as helpful for enriching the computational reservoir. Here we find that when learning temporally extended output patterns with a reservoir computing scheme, WP can learn as well as or better than NP and NPc.

We consider a recurrently connected reservoir of *N* = 500 rate neurons driven by five external inputs of length *T* = 500. Inspired by the behaviorally relevant task of reproducing a movement from memory - here drawing a figure - the task is to generate the *x* and *y* coordinates of a butterfly trajectory [42, 43] at the *M* = 2 outputs by training a linear readout (Fig. 7a). The trajectory is non-trivial in that it is not realizable from the external inputs. In fact, it requires reading out from many reservoir modes (Fig. 7b, dashed gray line).

**Figure 7.**
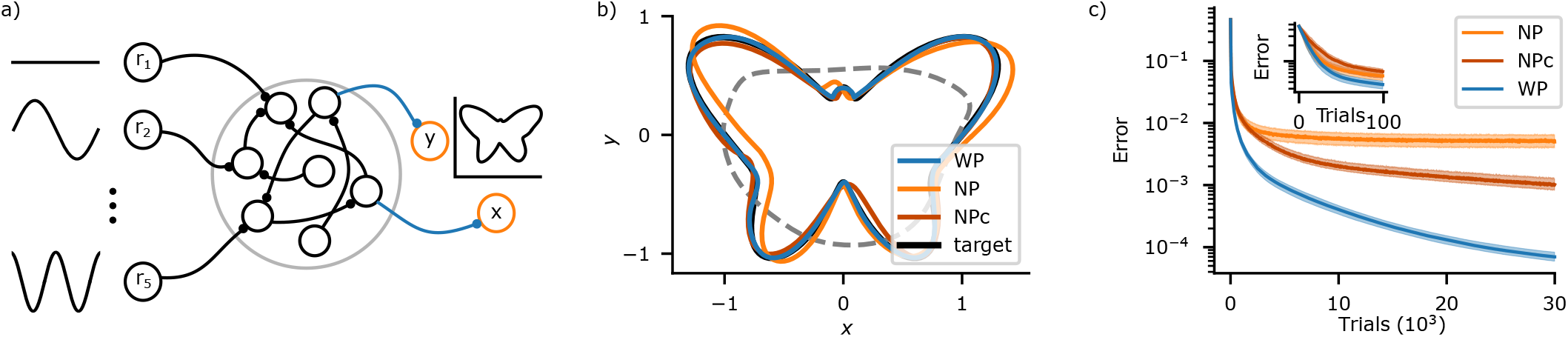
WP outperforms NP on a reservoir computing-based drawing task. a) Schematic of the recurrent, fixed reservoir receiving five external inputs. Only readout weights are learned. b) Target (black) and final outputs of WP (blue), NP (orange) and NPc (red). A least squares fit (gray, dashed) using only the first five principle components of the reservoir dynamics demonstrates that the task critically depends on reading out further, weaker dynamical components. c) Error curves on a logarithmic scale. WP reaches a lower final error than NP and NPc, with NPc improving on NP, cf. also (b). Inset: Early error evolution. There is a considerable improvement already during the first 50 trials. The curves show median (solid) and interquartile range between first and third quartile (shaded) over 1000 runs of the same network with different noise configurations.

Formally, the task is similar to the setting discussed above, with the difference that there is a wide distribution of different, nonzero input strengths 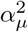. The evolution of expected error is then best described by splitting the error 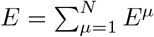 into different error components, each of which is associated with the weights that read out from a latent input *r_μt_* (SM 6, Eq. (S173)). In WP and NP the evolution of the error components follows a matrix exponential where different components decay at different rates and interfere with each other. Components that decay relatively quickly may be the main source of improvements in the beginning of training, whereas more slowly decaying components dominate the error towards the end. This effect can be seen in the approximately piecewise linear error decay in the logarithmic plot in Fig. 7c.

Fig. 7c compares the performance of WP, NP and NPc in the drawing task. Perturbation size is finite, *σ*_eff_ = 5 × 10^−3^. WP converges faster initially, which may be typical for tasks with distributed input strengths (SM 6, Eq. (S187)). It also achieves a lower final error. This is compatible with the observation that the effective dimension of the reservoir dynamics, as measured by the participation ratio (PR ≈ 5), is much smaller than the temporal extent of the task: the resulting smaller effective perturbation dimension of WP (*M*PR vs. *MT* for NP vs. 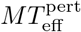 for NPc) yields an advantage for WP (Fig. 1b,Fig. 4a). NPc reaches a lower final error than NP. We have optimized its 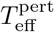 using a parameter scan, SM, Fig. S8. From our simulations we cannot completely exclude that the improvement is (in part) due to an effective decrease in learning rate, resulting from a longer correlation time in the perturbations than in some relevant inputs. For the biologically less relevant case of infinitesimal perturbation sizes, the performances of WP and NP are similar, SM, Fig. S7 (compatible with Fig. 1 with *b* = 0). Towards larger trial numbers the convergence of WP becomes slower: WP has difficulties with adjusting weights mediating weak inputs, since the impact of their perturbation on the error is small; the same effect underlies the weight diffusion in Fig. 2. Simulations indicate that the convergence is only slower by a constant factor of the order of 1 and that the optimal learning rate can be well estimated from the participation ratio (SM, Fig. S9).

### Delayed non-match-to-sample task

To ensure analytical tractability and for simplicity, so far we made a few biologically implausible assumptions. Specifically, only connection weights to linear units were trained, each trial consisted of a perturbed and an unperturbed run and mostly the exact same input was used in each trial. In the following we show that our findings generalize to settings without these assumptions. For this we consider the learning of a delayed non-match-to-sample (DNMS) task (temporal XOR) by nonlinear recurrent networks. DNMS tasks and closely related variants are widely used both in experiment [44] and theory [14, 45], where they serve as simple working memory-reliant, not linearly separable decision making tasks. We use the same setting as [14], which shows that a new variant of NP is able to solve the DNMS task. In particular, the setting is not adjusted to WP. The network consists of 200 nonlinear rate neurons receiving input from two external units *u*_1_ and *u*_2_. One of the network neurons, whose rate we denote with *z*, serves as its output (Fig. 8a). In each trial, the network receives two input pulses, where each pulse is a 200 ms long period with either *u*_1_ or *u*_2_ set to 1, and subsequently has to output 1 for 200 ms if different inputs were presented and −1 if the same inputs were presented (Fig. 8b). There is a 200 ms long delay period after each input pulse.

**Figure 8.**
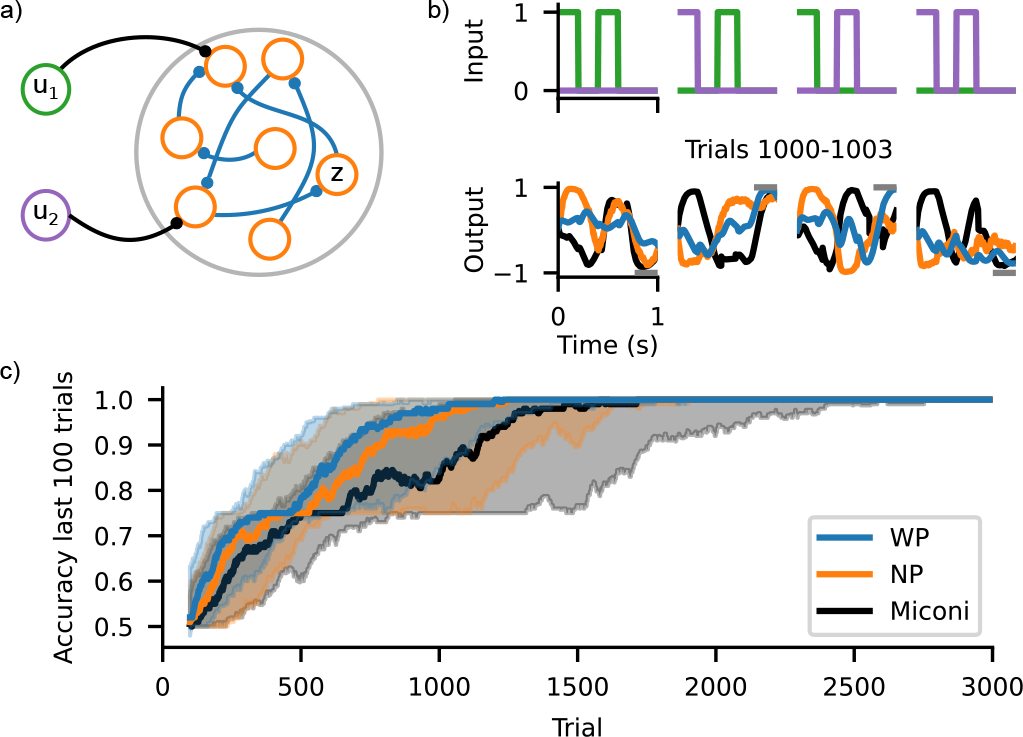
WP performs as well as NP on a DNMS task. a) Schematic of the recurrent network with inputs *u*_1_ and *u*_2_ and output *z*. All network weights are learned, i.e., for WP, all network weights (blue) are perturbed and for NP, all network nodes (orange) are perturbed. b) Inputs and outputs during example trials. Top row: Inputs *u*_1_ (green) and *u*_2_ (purple) for the four different trial types. Bottom row: Outputs for WP (blue), NP (orange) and the version of NP proposed by [14] (black) for trials 1000–1003 for the inputs shown above. Gray bars show target outputs. c) Accuracy during training. WP (blue) performs similarly well as NP (orange) and the version of NP used by [14] (black). There is a noticeable transient slowdown at an accuracy of 75 %, which corresponds to the successful learning of three out of the four different trial types. Solid lines show the median and shaded areas represent the interquartile range between first and third quartile using 100 network instances.

We train all recurrent weights using the usual update rules (Eqs. (3,6)), but replace the error of the unperturbed trial by an exponential average of the errors of the previous trials [12–14]. Hence, each trial now only consists of a perturbed and not additionally an unperturbed run. We first assume that the exact perturbations *ξ* are accessible for the weight update, which seems biologically plausible for WP (cf. Discussion), but less so for NP (cf. Discussion and [14]). Therefore we also compare WP and NP to the biologically plausible version of NP proposed by [14], which avoids this assumption: in the weight update rule, it approximates the exact node perturbations *ξ*^NP^ with a nonlinearly modulated difference between the momentary input to a neuron and its short term temporal average (see Methods for more details).

Fig. 8c shows the performance of the three update rules in terms of their accuracy over the last 100 trials, where a trial is considered successful if the mean absolute difference between *z* and the target output is smaller than 1. We find that all update rules learn the task comparably well and reach perfect accuracy within at most 2000 trials when considering the median of network instances. Thus, our previous findings that WP can perform as well as or better than NP in simplified settings extend to the considered biologically plausible setup. That means WP can perform well for nonlinear neuron models, recurrent connectivity and when the error of the unperturbed network is not available. Furthermore, the results indicate that approximating the perturbation as in [14] only mildly impacts the performance of NP for the considered task. Finally, we applied NPc to the task, which did not yield an improvement over NP (SM, Fig. S12). Together with the similar performance of WP and NP, this indicates that the temporal dimension of the perturbation has little effect on task performance, perhaps because the period in which the target value needs to be assumed is rather short and the output is otherwise unconstrained.

### MNIST

Finally, we apply WP and NP to MNIST classification. We use batches of images to train the networks. Each time step thereby corresponds to the presentation of one image and the networks receive error feedback only at the end of a batch. This allows us to test how well WP and NP work on a more complicated, temporally extended task and in networks with a multi-layer structure. In addition, it allows us to study how our analytical results for the learning of multiple input patterns (Sec. “Multiple subtasks”) extend to real-world tasks.

We use a two-layer feed-forward network with 10 output neurons and 100 neurons in the hidden layer (Fig. 9a). It learns via the rules Eqs. (3,6), where *T* equals the batch size *N*_batch_. Hence, the perturbation is different for each image in the case of NP, while it is the same for WP. We test WP and NP for batch sizes of *N*_batch_ ∈ {1, 10, 100, 1000}. For each batch size we determine the best-performing learning rates *η* and perturbation strengths 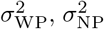 via grid searches. The perturbation strength has, however, little impact on performance, indicating that the final error is not restricted by reward noise due to finite size perturbations (Eqs. (13,14)).

**Figure 9.**
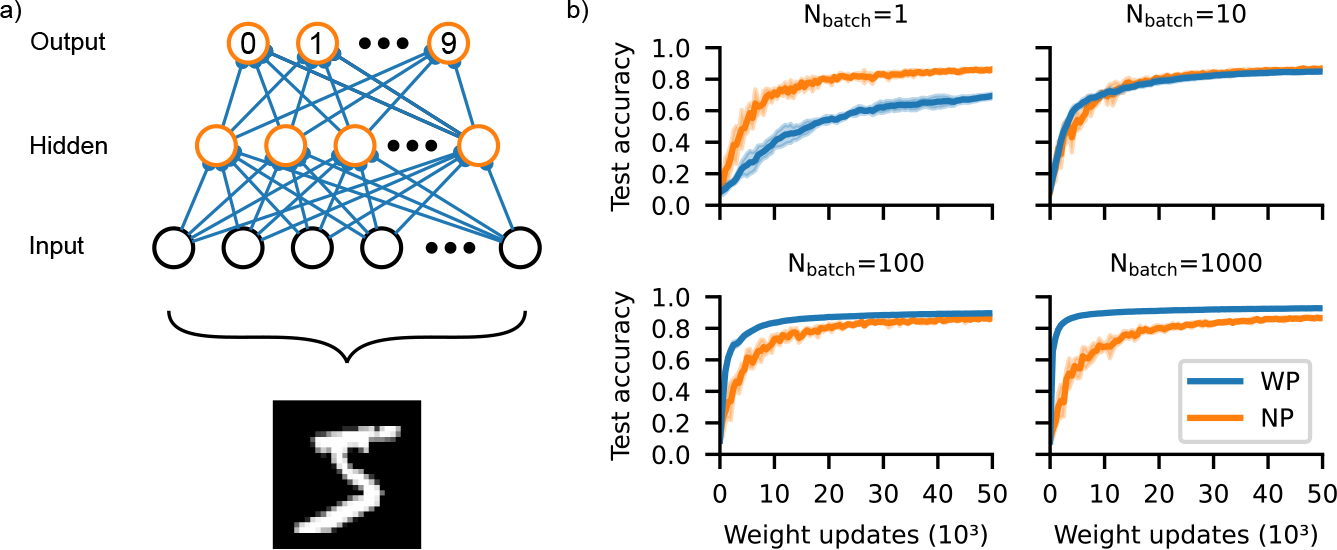
WP can outperform NP on MNIST. a) Schematic of the used fully connected, two-layer network. All network weights are learned, i.e., for WP all network weights (blue) are perturbed and for NP all network nodes (orange) are perturbed. b) Test accuracy as a function of the number of weight updates for WP (blue) and NP (orange) for different batch sizes. NP does not profit from increasing the batch size and always reaches a final accuracy of ≈86 %. WP improves considerably with increasing batch sizes and reaches a final accuracy of ≈92 % for *N*_batch_ = 1000. Solid lines show the mean and shaded areas show the standard deviation using 5 network instances.

We find that for WP the performance improves drastically with increasing batch size, Fig. 9b. The final test accuracy is only about 69 % for a batch size of 1 but reaches 92 % for *N*_batch_ = 1000. Simultaneously the optimal learning rate increases strongly, by a factor of approximately 50 (SM, Fig. S10c and Supplementary Table S2). For comparison: the biologically implausible stochastic gradient descent (SGD) rule reaches accuracies of 95 %–98 % for the considered batch sizes. In contrast, the learning curves of NP appear to be entirely independent of the batch size (Fig. 9b); the final test accuracy is always about 86 % and the optimal learning rate is constant as well. We also applied NPc to the task. The inputs are temporally uncorrelated because the elements of the batches are drawn randomly. Based on our previous observations for uncorrelated input, Fig. 4c, we therefore expect that NPc performs similar to or worse than NP. The numerical experiments confirm this: performance deteriorates with increasing perturbation correlation time; the effect is more pronounced with larger batch size (SM, Fig. S12). To conclude, larger batch sizes, as commonly used in machine learning, favor WP, while smaller batch sizes favor NP (and NPc).

An improvement of WP with batch size and NP’s independence of it are in agreement with our theoretical analysis Sec. “Multiple subtasks”. However, from this analysis we also expected that WP’s learning rate can reach at most that of NP for large batch size. NP’s slower convergence suggests that it is more susceptible to deviations of the network architecture from linear, single-layer networks. Indeed, when using single-layer networks, NP’s performance improves, while the opposite holds for WP and SGD (SM, Fig. S11). In a single layer linear network with realizable targets NP performs better than WP even for large batch sizes (SM, Fig. S11), consistent with our analytical findings that training with different subtasks (here: different batches) harms WP (Sec. “Multiple subtasks”) while the absence of unrealizable targets benefits NP (Sec. “Unrealizable targets”).

The results are particularly remarkable when naively comparing the number of perturbed nodes and weights (“Introduction”): For the network considered here, there are only 110 output and hidden nodes, but 79 510 weights (including biases). Nevertheless WP can clearly outperform NP. Also a comparison of the actual perturbation dimensions cannot explain the better performance of WP in, e.g., Fig. 9b lower left (WP pert. dim.: 79 510, NP pert. dim.: 110 × *T* = 11 000).

## DISCUSSION

Our results show that WP performs better than NP for tasks where long trials capture most of the task’s content. This might seem paradoxical as NP incorporates more structural knowledge on the network, namely the linear summation of inputs. However, WP accounts for the fact that the weights in a neural network are (approximately) static. Further, by perturbing the weights it implicitly accounts for low input dimensionality and generates only realizable output changes. Therefore it generates better tentative perturbations. This leads to less noise in the reward signal and better performance (smaller final error and sometimes faster convergence) in the tasks where WP is superior to NP.

Our theoretical analysis shows that the lower noise in WP firstly results from an effective perturbation dimension that is lower than NP’s if the temporal extent of a task is larger than its input dimensionality, *T* > *N*_eff_. Secondly, factors such as the attempt of NP to realize unrealizable targets contribute. Temporally extended tasks with durations on the order of seconds and low dimensionality occur frequently in biology, for example in motor learning and working memory tasks. In line with perturbation-based learning, biological movements are endowed with noise, which helps their learning and refinement [46]. The associated neuronal dynamics in the brain are confined to a low-dimensional space, a property shared by many types of biological and artificial neural network activity [47–49]. The dynamics for simple movements as investigated in typical experiments are embedded in spaces of dimension of order 10 [19]. This indicates low effective input dimensionality *N*_eff_ at the different processing stages. The effective muscle activation dimensionality is similarly low [20, 50]. Neurons under in vivo conditions can faithfully follow input fluctuations on a timescale of 10ms [51] and significant changes in neuronal trajectories happen on a timescale of 100ms [19, 21, 52]. For the learning of a movement of duration 1s, this suggests an effective temporal dimension of about 10 to 100 similar to the expected input dimension. This implies that both WP and NP/NPc are promising candidates for the learning of simple movements. Our results indicate that WP will be superior if the movements are longer lasting or lower-dimensional.

We have explicitly studied the learning of movement generation (drawing task) and of a working memory task (DNMS). The numerical simulations show that WP performs similarly well or better compared to NP. In a task generally investigating the learning of complicated nonlinear, temporally extended input-output tasks (MNIST), WP outperforms NP as soon as the tasks have sufficient temporal extent.

As another concrete application, consider the learning of the single song in certain birds. A single, stereotypical input sequence in a “conductor area” (HVC) may drive the circuit [35, 53]. The effective input dimension *N*_eff_ is thus at most as large as the temporal dimension *T* of the task. Based on recent experiments, ref. [53] proposed that the output of the tutor/experimenter area (LMAN) is modified by reinforcement learning via NP, such that it guides the motor area (RA) to learn the right dynamics. Our analytical results predict that WP is as well or better suited to achieve this task since *N*_eff_ ≤ *T*. Earlier work suggested that WP [22] or NP [17] may directly mediate the learning of the connections from HVC to RA. Due to HVC’s very sparse activity, WP0 would be highly suitable for such learning. Reward-based learning of mappings between conductor sequences and downstream neural networks may also be important for different kinds of precisely timed motor activity [54, 55] and for sequential memory [56, 57].

Biological neural networks are inherently noisy. We found that WP and NP induce two types of weight update noise, credit assignment and reward noise. We understood their impact analytically; additional feedback or output noise implies additional reward noise with alike effects. We have additionally studied the impact of input noise on the learning of linear networks. Our simulations show that the convergence time and the final error increase with the input noise strength. The increase is smaller in WP than in NP. We further find that our results with zero noise are recovered in the limit of small noise compared to the strength of the relevant latent inputs, if the learning time is finite. The same holds for WP also if the weights are appropriately limited, for example due to weight decay. The fact that animals can learn to perform tasks with high precision, i.e. with small final error, indicates that the case of small noise may be the biologically relevant one. The results of WP and NP learning with noisy inputs can be understood analytically from our findings on inhomogeneous input distributions (SM 6). Also the considered DNMS and MNIST tasks contain input noise: the DNMS task because of the randomly chosen initial conditions, the MNIST task because of the naturally noise-afflicted input images. We conclude that the finding of similar or better performance of WP compared to NP in temporally extended, low-dimensional tasks may readily apply to biologically plausible, noisy networks.

WP and NP have biologically plausible implementations. NP requires that the plastic synapses can keep track of their input and the somatic perturbations (which may arrive from a tutor/experimenter neuron). Biologically plausible mechanisms for this have been proposed both for tasks with immediate reward [12, 13] and reward at a temporally extended trial’s end [14]. Their underlying idea is to assume that the perturbation fluctuates more quickly than the other input. The present fluctuation can then be approximately isolated by subtracting a short-term temporal average of the past overall input from the present one [12, 13]. This difference replaces the injected perturbation in the eligibility trace. For tasks with late reward, the eligibility trace needs to integrate a nonlinearly modulated version of the described difference [14]. This prevents the cancellation of a perturbation’s effect by the subsequent change in the average that it evokes, because the peak in the original perturbation is sharper and higher than the one in the average. We use this learning model of [14] in Fig. 8. The biological implementation of WP may be even simpler. A neural network needs to generate labile random weight changes and keep track of them. They should be approximately constant during a task and enhanced, deleted or reversed by a subsequent reward signal. Experiments on timescales from minutes to days found spontaneous changes in the synaptic weights, which have similar strength as changes due to activity dependent plasticity [31]. Such changes might generate the perturbations required for our WP scheme. Previous work suggested also synaptic unreliability to provide the perturbations for WP [58]. This fits into our scheme of static weight perturbations if neurons spike once during a trial or if they burst once and the synaptic transmission is restricted to a single time bin. Another source of the required randomness may be fluctuations of activity-dependent plasticity, while the deterministic baseline acts as a useful prior. If the baseline is unrelated to the task, it will with high probability be orthogonal to task-relevant directions (due to the high dimensional weight space) and not harm learning, similar to the weight diffusion in WP. In this way, the fluctuations of activity-dependent plasticity, rather than their deterministic part, may be the source of learning.

Modulation of weight changes by reward has been observed in various experiments [59, 60]. As an example, the potentiation of synapses is enhanced or reversed depending on the presence or absence of a temporally close dopamine reward signal [61]. Also other factors play a role; potentiation can for example be reversed within a “grace period” of tens of minutes by a change of environment [62]. The consolidation and amplification of changes may be dependent on plasticity related proteins, which are upregulated by reward and for which the synapses compete (synaptic tagging hypothesis) [60, 63]. A posteriori modifications of tentative synaptic weight changes are also assumed in the reinforcement learning scheme of reward modulated Hebbian plasticity [64, 65], which is closely related to WP.

WP applies a random perturbation vector to the weights, measures the error change to obtain a reinforcement signal and applies as weight update the perturbation modulated by the reinforcement signal. The improvement therefore follows on average the weight gradient. A related approach is to randomly perturb and accept the perturbation if it leads to a better performance [22]. This simple instance of an evolutionary strategy [66, 67] is also applicable if there is no gradient. Our results for WP suggest that this and related evolutionary learning strategies might benefit from tasks that are low-dimensional, as reported previously [68], and not be harmed by their temporal extension. The sketched simple evolutionary learning may in the brain generate structural improvements: Experiments show that synaptic turnover occurs in presence but also spontaneously in absence of neuronal activity [69–72]. This may reflect the random elimination and creation of synapses and their consolidation by activity-dependent plasticity [73–78]. The basis of consolidation is that mainly weak synapses are removed such that strengthening through Hebbian learning causes the long-term presence of a synapse. Furthermore, Hebbian learning counteracts spontaneous synaptic weight changes, which could otherwise weaken useful synapses and ultimately lead to their removal. Network models show that restructuring with subsequent selective consolidation can recruit sparse available connectivity for task learning, prevent catastrophic forgetting and may explain the benefits of dividing learning into several temporally distinct phases [73, 75, 77]. Further, it may explain the experimental observation that there are commonly multiple synapses between connected neurons [74, 76, 78]. The signal for the strengthening of a tentatively established synapse may be interpreted as a reinforcement signal for its presence. This signal is generated if the pre- and postsynaptic neuron are coactive, due to external stimulation or recall mediated by other, already strengthened synapses. In contrast to WP, the reinforcement signal is therefore specific to an individual synapse, which simplifies learning. Specifically, any useful new synapse will be independently consolidated, while in WP tentatively applied useful weight changes can be reverted due to harmful ones in other parts of the weight perturbation vector. In a related model, ref. [79], consolidatory strengthening was implemented with NP instead of Hebbian learning. This allows to learn tasks based on a global reward signal. The synaptic weight fluctuations in the model induce random changes in task-irrelevant directions similar to the weight diffusion that we observe in WP. Our results suggest to directly exploit the synaptic weight fluctuations for consolidatory synaptic strengthening by using WP when learning low-dimensional and temporally extended reward-based tasks.

WP has been proposed in several variants. They differ in: (i) the task setup, for example instantaneous [7, 9] or temporally extended tasks [8, 22, 58], (ii) the implementing network, for example rate [22] or spike-based [58, 80] networks, (iii) the perturbation scheme, where all weights are simultaneously perturbed [7, 8, 58] or only one weight at a time [15], (iv) the computation of the weight update, by correlating reward and perturbation [7–9, 11, 58] or direct estimation of the gradient components (for the single weight perturbation scheme) [9, 15], (v) the estimation of the success of the perturbed network, which may involve a comparison of the obtained reward to an unperturbed baseline [8, 9] or a running average [22, 58] or it may consider the reward only [7, 58] and (vi) the weight update, which may be proportional to the success of the network [7–9, 11, 58] or independent of its size as long as there is an improvement [22]. A similar diversity of NP variants exists [9–14, 16, 17, 24, 79, 81, 82].

The tasks considered in our article are temporally extended. The reward is provided at the end of the trial, but influenced by earlier output states. This is consistent with many tasks in biology [1, 5, 14, 22] and with the learning schemes by [8, 10, 14, 22]. We choose a WP rule that is biologically plausible as it involves simultaneous perturbations to all weights and correlates reward and weight change. The success measure compares the obtained reward to the reward of an unperturbed network in order to reduce the update noise [8, 9]. In particular this avoids that unfavorable perturbations are associated with positive reward feedback. Finally, the weight update is proportional to the measured success in order to ensure that it occurs on average parallel to the reward gradient. The choices are identical to those by [8, 9] for temporally not extended tasks. Specifically the results in ref. [9] appear as special case of our results for multiple input patterns, if the task-dimension is maximal, single trials have no temporal extent and the inputs have fluctuating amplitude (see Sec. “Comparison with ref. [9]”).

We choose the NP scheme such that it matches the WP scheme. It is a discrete-time version of the NP scheme proposed by [10] and an extension of the scheme by [9] to temporally extended tasks. In biologically plausible implementations of WP and NP, the reward should be compared to an intrinsically generated prediction, such as an average of previous rewards [12–14] or the reward of another perturbed trial [82]. In the delayed non-match-to-sample task, we thus replace our standard unperturbed baseline by such an average. This also allows a direct comparison with the NP scheme by [14]. In Sec. “Input noise” the perturbed and unperturbed trials have different input noise, such that *E* is no longer the exact unperturbed counterpart of *E*^pert^.

To exploit correlations in the inputs with a node perturbation learning rule, we introduced NPc, which is identical to standard NP apart from using temporally correlated, smoothed node perturbations. We find that the temporal correlations are usually beneficial if also the inputs are (similarly) correlated. This is in contrast to ref. [13], which observed a detrimental effect already of short correlations for a node perturbation variant that relies on high frequency perturbations. Other previous studies injected white noise perturbations only [10, 12, 16, 17]. Our simulations with linear networks indicate that the perturbation correlation time of NPc should optimally match that of the inputs; NPc then performs similar to NP in a task with reduced temporal extension.

NP is studied in various concrete neurobiological settings. Previous work used feedforward networks with NP to model the learning of coordinate transforms in the visual system [83], birdsong [17, 53] and motor output [12, 84]. Ref. [13] shows that reservoir computers with NP trained, fed back readouts can learn periodic inputs, routing and working memory tasks. [14] uses a fully plastic recurrent network for the learning of a delayed non-match-to-sample, a selective integration and a motor control task. Finally, NP is often employed for reference and comparison [85–91]. WP is considered less in studies of neurobiological learning. It is implemented in early feedforward network models of birdsong [92] and binary output task learning [58, 80]. Further, it is occasionally used for comparison [86, 88]. Very recently ref. [4] has shown that recurrent neural networks can be pretrained with WP and the reservoir computing scheme to thereafter learn with static weights to generate fixed point activity.

The results of our present article using feedforward, reservoir computing and fully plastic recurrent networks suggest that for many tasks WP is at least as suitable as NP, while the implementation may be even simpler. This indicates that WP is a useful benchmark and a similarly plausible model for learning in the brain as NP. Experimentally measurable features of the learning and weight dynamics may allow to distinguish the learning rules in biological neural networks.

## MATERIALS AND METHODS

### Analytical error dynamics

To analytically compute the dynamics of the expected error, we consider an arbitrary perturbation *ξ*. This determines the error change *E*^pert^ − *E* and the resulting weight update Δ*w* via Eqs. (3,6). Δ*w* in turn determines the new weights and via Eq. (8) the error *E*(*n* + 1) after the update. *E*(*n* + 1) is thus a function of *ξ*, the weight mismatch *W* (*n*) before the update and the input correlations *S*,

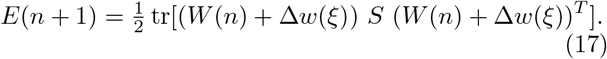

Averaging over perturbations and using Isserlis’ theorem yields an equation for the expected error 〈*E*(*n* + 1)〉. When assuming that all latent inputs have the same strength, 〈*E*(*n* + 1)〉 becomes a function of the error 〈*E*(*n*)〉 before the update and the system parameters, leading to Eq. (9). The detailed derivation is given in SM 2.

### Numerical simulations accompanying the theoretical analysis

In the numerical experiments in Sec. “Theoretical analysis”, the *N*_eff_ nonzero inputs are orthonormal functions, superpositions of sines, scaled by *α*^2^ = *N/N*_eff_ to keep the total input strength *α*^2^*N*_eff_ for different *N*_eff_ constant. Targets *z_it_* are obtained by linearly combining these functions using teacher weights 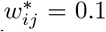 and adding as unrealizable component a further, appropriately scaled, orthonormal function. Learning rates are *η*\*.

### Input and perturbation correlations

NPc is applicable to general network dynamics (cf. Figs. 7,S7,S8,S12). In Fig. 4 we apply it to linear networks with correlated inputs that are constructed similar to the perturbations, by low-pass filtering *N*_eff_ white noise traces (filtering time constant 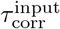, effective temporal dimension 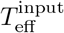). Subsequently we orthonormalize them, which somewhat modifies the correlation times. The realizable components of a target are linear combinations of these correlated inputs weighted by 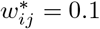. We assume that there is an additional unrealizable component, which, for simplicity, contains all modes orthogonal to the inputs with equal strength, such that *E*_opt_ = 2.

For 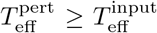, NPc operates at 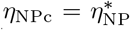, the optimal learning rate for NP. For 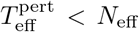, we use the optimal learning rate of NP for a task with reduced 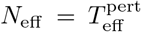, i.e. 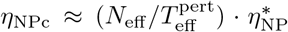. Our intuition is that perturbations with temporal dimension 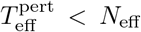 can only improve an 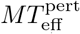-dimensional subspace of the weights. For NPc 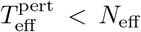 therefore reduces the learnable number of weights from *MN*_eff_ to effectively 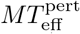 independent ones. This is like in Fig. 1c, gray curves, where *T* restricts *N*_eff_. As effectively fewer weights are learned, a higher learning rate can be chosen. We performed additional simulations with an unadjusted (smaller) learning rate that confirmed our choice, as convergence otherwise became much slower.

For a given 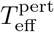, simulations of 20 000 trials are repeated 1000 times (randomly generated inputs change between runs but not within a run). The final error of a run is computed by averaging over the last 500 trials (in which the error is approximately constant, SM, Fig. S4) to determine the mean over runs and its SEM. To determine 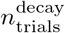, we use 10 samples of 100 runs each. For each sample, we compute the mean error over runs and additionally smooth it with a centered temporal running average of window size 20. 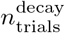 is then the trial for which the described average drops for the first time below *E*_*f*,unr_ + 0.05 · (*E*(0) − *E*_*f*,unr_). Fig. 4b reports the mean and standard error of the mean of 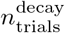 over all samples. Fig. 4c repeats the same analysis for NP with *η* varied from 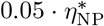 to 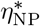. Gray curves in Fig. 4c (NPc with *η* adjusted by a factor of 0.5, 0.6, · · ·, 1.2) used 100 repetitions.

### Input noise

We extend the basic theory task, in which a single mapping from an effectively *N*_eff_-dimensional input signal *r_jt_* onto target outputs 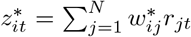 is learned, by adding independent white noise to each input at each time-step,

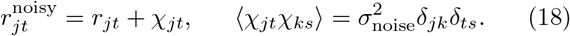

As the added white noise has a rotationally symmetric distribution in the space of input neurons, we can still without loss of generality rotate the input space such that each of the first *N*_eff_ input neurons carries a signal component and additional noise, while the remaining inputs are purely noisy. We note that because the noise in the task-relevant inputs is amplified by the weights, their optimal values are closer to zero than those of the noise-free task.

SNR is defined as the ratio of the total (summed) power in the input signal to that in the noise. Averages are taken over 100 repetitions (a,b,c), the last 1000 of 100 000 trials (a,b) and over irrelevant weights (c).

### Reservoir computing task

The *N* = 500 rate neurons of the fully connected recurrent reservoir network evolve according to

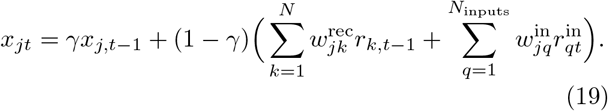

The rate of neuron *k* is *r_kt_* = tanh(*x_kt_*). Their decay time constant is *τ* = 10 time steps, i.e. *γ* = e^−1/*τ*^. Recurrent weights *w*^rec^ are drawn from a centered normal distribution, the weight matrix is thereafter normalized to ensure that the real part of its largest eigenvalue is *g*_rec_ = 1. Input weights *w*^in^ are drawn from 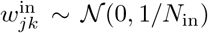. Creating various instances of such random networks showed that performance and participation ratio 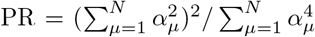 are rather independent of the instance. The participation ratio gives an estimate of the dimensionality of the reservoir dynamics [19, 93]. Generally we observe PR ≈ 5, for example in the network of Fig. 7 PR ≈ 5.3. The *N*_in_ = 5 inputs to the reservoir are orthogonal to each other, 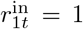, 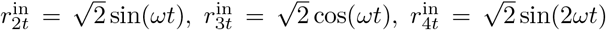, 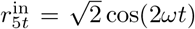, *ω* = 2*π/T*, *T* = 500 timesteps. The trained linear readout produces *M* = 2 outputs 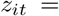 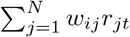. Their target 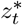 is, up to scaling, the same as in [42]: 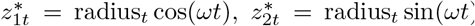, with radius_*t*_ = 0.1 · (9 − sin(*ωt*) + 2 sin(3*ωt*) + 2 sin(5*ωt*) − sin(7*ωt*)+3 cos(2*ωt*)−2 cos(4*ωt*)). Already 100 timesteps before the task starts, the reservoir is initialized and given external input. By the time the task begins, network activity is enslaved by the external input and has settled down to a periodic orbit. Technically, we record the reservoir activity traces *r_jt_* once for the entire training of *w*, because they are the same in each trial. The value of the participation ratio motivated us to construct an optimal readout reading out the largest five principal components via the least squares fit 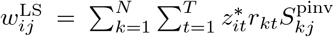 (Fig. 7b, dashed gray). Here *S*^pinv^ is the pseudoinverse of the reduced correlation matrix of the reservoir that is obtained by setting all eigenvalues of *S* except the largest five to zero. Including six principal components did not qualitatively change the result.

From the theoretical analysis Eq. (12), we obtain an estimate 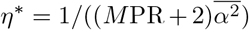 for the optimal learning rate, by setting *N*_eff_ → PR and 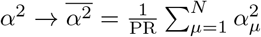. 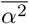 is the strength of each latent input when we assume that the total input strength is generated by PR equally strong ones. We verified by a grid search that this estimated value yields for both WP and NP close to optimal performance, as measured by the error after 10 000 trials with infintesimally small 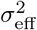, showing that it indeed maximizes convergence speed. We therefore choose it as the learning rate for our task with infinitesimal (SM, Fig. S7) and also with finite perturbation size (Fig. 7), since the theoretical analysis yielded independence of *η*\* from 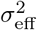 (Eq. (12)). For NPc we used the same learning rate as for NP and WP and determined the optimal 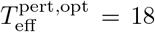 by minimizing the error after 30 000 trials (SM, Fig. S8). Simulations with finite perturbations used *σ*_eff_ = 5 × 10^−3^. A scan over *σ*_eff_ confirmed that the final error depends quadratically on it, as predicted by the theory.

### Delayed non-match-to-sample task

The fully connected recurrent network has *N* = 200 rate neurons. The dynamics of neuron *i*, *i* = 4,…, *N*, are governed by

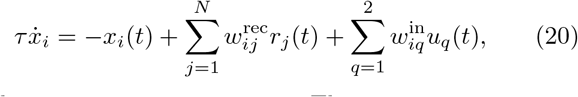

with time constant *τ* = 30 ms. The constant activations *x*_1_(*t*) = *x*_2_(*t*) = 1 and *x*_3_(*t*) = −1 provide biases [14]. The rate of each neuron *i*, *i* = 1,…, *N*, is given by *r_i_*(*t*) = tanh(*x_i_*(*t*)). *z*(*t*) = *r*_4_(*t*) is the network output. We use the forward Euler-method with stepsize *dt* = 1 ms to simulate the dynamics and draw the initial activations from a uniform distribution, 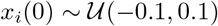 for *i* = 4,…, *N*. Recurrent weights are drawn from a Gaussian distribution, 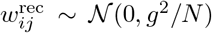, with *g* = 1.5. Input weights are drawn from a uniform distribution, 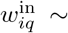 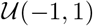.

All recurrent weights 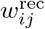 are trained. The error function of WP and NP is the mean squared difference between the output *z* and the target within the last 200 ms of each trial. For each of the different trial types *k*, *k* = 1,…, 4, we use an exponential average of the previous errors *E*^pert^(*n*_k_) for this trial type (*n*_k_ indexes the trials of type *k*) as the error baseline:

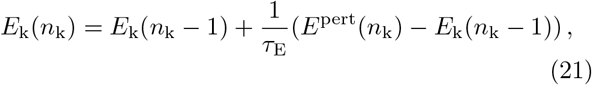

where *τ*_E_ = 4. To get the best performing learning parameters, we performed a grid search, which yielded *η*^WP^ = 1 × 10^−5^, *σ*^WP^ = 4.64 × 10^−3^, *η*^NP^ = 1 × 10^−5^, *σ*^NP^ = 4.64 × 10^−1^.

For the details of the version of NP proposed by [14], see this article. For the convenience of the reader here we briefly mention the main differences to the vanilla NP version Eq. (6): For each network neuron a node perturbation is applied at a simulation time step only with a probability of 0.3 % and is drawn from a uniform distribution, 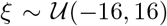. The error is given by the absolute difference between output and target. Weight updates are computed via 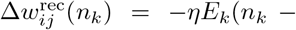 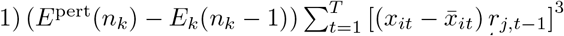, and clipped when they exceed ±3 × 10^−4^ (cf. code accompanying [14]). *t* indexes the simulation time step of each trial, *T* is the total number of simulation time steps per trial and 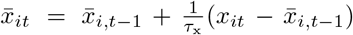 is an exponential average of past activations. Parameter values are *η* = 0.1, 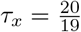.

### MNIST classification task

The input layer of the fully connected feedforward network consists of 784 units encoding the pixel values of the data. The hidden layer consists of 100 neurons with tanh activation function and biases. The output layer consists of 10 neurons, one for each single-digit number, with softmax activation function and biases. We use the standard training and test data set, but split the standard training data set into a training data set of 50 000 images and a validation data set of 10 000 images. No preprocessing is done on the data. We employ vanilla WP Eq. (3), NP Eq. (6) or SGD to train all parameters of the network. The error function is the cross-entropy loss averaged over the batch of length *N*_batch_ = *T*. We also tried to combine the gradient estimates obtained from WP and NP with Momentum, RMSProp or Adam [29], but did not find an improvement of performance compared to the vanilla versions with carefully tuned parameters. The same holds for SGD. This may be because of the rather simple network architecture.

To obtain the best-performing parameters (the learning rate for all three algorithms and the standard deviation for WP and NP), we performed a grid search for each of the considered batch sizes: For each parameter set we trained the network for 50 000 trials (i.e.: weight updates) on the training data set. We then selected the best-performing parameter sets based on the final accuracy on the validation data set and applied them to the test data set. High final accuracy appeared to concur with fast convergence speed, such that a comparison to our analytical results (where learning rate optimizes the convergence speed) seems justified.

## I. ACKNOWLEDGMENTS

We thank the German Federal Ministry of Education and Research (BMBF) for support via the Bernstein Network (Bernstein Award 2014, 01GQ1710).

## Notes

### Competing Interest Statement

The authors have declared no competing interest.

### Summary of Updates

Update author information on bioRxiv; Main and Supplementary PDF remain unaffected

