## Supplementary Material for "Weight Perturbation Learning Performs Similarly or Better than Node Perturbation on Broad Classes of Temporally Extended Tasks"

The supplement follows the structure of the main text; main results and equations referenced in the main text are highlighted for clarity with a yellow background. The Supplementary Material (SM) has eight parts, SM1-SM8, each with several sections. Further there are twelve supplementary figures, Figs. S1–S12 and two supplementary tables Tabs. 1,2. The first table, Tab. 1 below, gives an overview of the assumptions used in the different parts and sections of the supplement.

| Part / Section | Linear networks | Repeated inputs | Same strength inputs |
|---|---|---|---|
| 1: "Learning models and task principles" | | | |
| "Mean updates...", "Dependence of..." | | × | |
| "Task setting", "Effective pert..." | × | × | × |
| 2: "Derivation of error dynamics" | | | |
| "Error curves for gradient descent" – "...for node perturbation" | × | × | |
| "...for equally strong input components", "Optimal learning rate" | × | × | × |
| 3: "Analysis of error dynamics" | × | × | × |
| 4: "Analysis of Weight Diffusion" | × | × | × |
| 5: "Multiple subtasks" | × | | × |
| 6: "Arbitrary input strength distributions" | × | × | |
| 7: "Input and perturbation correlations" | × | × | (×) |
| 8: "Improved learning rules" | | | |
| "WP0: Assign zero credit to zero inputs" | | | |
| "Hybrid perturbation (HP): Using WP..." | × | × | (×) |

**Table 1.** Assumptions used in the different supplementary parts. Crosses, ×, mean that an assumption is used. If different sections of a part use different assumptions, all sections are specified. Parentheses mean that an assumption is used only in part of a section.

## SM 1

### Learning models and task principles

##### Mean updates for small perturbations

To calculate the average update of WP, one uses the linear order approximation valid for small perturbations ***Dembo and Kailath*** (***1990***); ***Cauwenberghs*** (***1993***),

$$E^{\mathrm{pert}} - E \approx \sum_{m=1}^{M}\sum_{k=1}^{N} \frac{\partial E}{\partial w_{mk}} \xi_{mk}^{\mathrm{WP}}, \tag{S1}$$

where the sum is over all weights in the network. With this and $\langle \xi_{ij}^{\mathrm{WP}} \xi_{mk}^{\mathrm{WP}} \rangle = \delta_{im}\delta_{jk}\sigma_{\mathrm{WP}}^2$, where $\delta$ is the Kronecker delta, the WP update rule, main text Eq. (3), yields

$$\langle \Delta w_{ij}^{\mathrm{WP}} \rangle \approx -\frac{\eta}{\sigma_{\mathrm{WP}}^2} \sum_{m=1}^{M}\sum_{k=1}^{N} \frac{\partial E}{\partial w_{mk}} \langle \xi_{mk}^{\mathrm{WP}} \xi_{ij}^{\mathrm{WP}} \rangle = -\eta \frac{\partial E}{\partial w_{ij}}, \tag{S2}$$

i.e. the weight update is along the negative error gradient. The result's independence of $\sigma_{\mathrm{WP}}$ motivates the division by $\sigma_{\mathrm{WP}}^2$ in Eq. (3).

NP perturbs for temporally extended tasks the total inputs $y_{it} = \sum_{k=1}^{N} w_{ik} r_{kt}$ of neurons $i = 1, \cdots, M$ by the node perturbation vectors $\xi_{it}^{\mathrm{NP}}$. Therefore the change in the scalar error is, to linear order, given by the projection of $\xi_{it}^{\mathrm{NP}}$ onto the error gradient $\partial E/\partial y_{it}$ with respect to $y_{it}$ (***Fiete and Seung, 2006***),

$$E^{\mathrm{pert}} - E \approx \sum_{i=1}^{M}\sum_{t=1}^{T} \frac{\partial E}{\partial y_{it}} \xi_{it}^{\mathrm{NP}}. \tag{S3}$$

Since $\partial y_{it}/\partial w_{ij} = r_{jt}$, the gradients with respect to weights and sums of inputs are related via the chain rule by $\partial E/\partial w_{ij} = \sum_t \partial E/\partial y_{it}\, r_{jt}$. This reveals that the NP weight update Eq. (6) is on average along the negative error gradient,

$$\langle \Delta w_{ij}^{\mathrm{NP}} \rangle \approx -\frac{\eta}{\sigma_{\mathrm{NP}}^2} \sum_{m=1}^{M}\sum_{s,t=1}^{T} \frac{\partial E}{\partial y_{mt}} \langle \xi_{mt}^{\mathrm{NP}} \xi_{is}^{\mathrm{NP}} \rangle r_{js} = -\eta \sum_{t=1}^{T} \frac{\partial E}{\partial y_{it}} r_{jt} = -\eta \frac{\partial E}{\partial w_{ij}}. \tag{S4}$$

Eqs. (S2,S4) hold for any error function. For corresponding more specific computations for the quadratic error function Eq. (8) see Sec. "Task setting".

##### Dependence of weight update noise on error baseline

To show that the choice of $E$ as error baseline in Eq. (3) minimizes the update noise for WP, we compute the variance $\langle\langle \Delta w_{ij}^{\mathrm{WP}} \rangle\rangle$ of the WP weight updates when adding a possibly trial- and synapse-dependent baseline term $\tilde{E}_{ij}$ to the update rule, $\Delta w_{ij}^{\mathrm{WP}} = -\frac{\eta}{\sigma_{\mathrm{WP}}^2}(E^{\mathrm{pert}} - E + \tilde{E}_{ij})\xi_{ij}^{\mathrm{WP}}$. The linear approximation for $E^{\mathrm{pert}}$ yields

$$\begin{aligned} \langle\langle \Delta w_{ij}^{\mathrm{WP}} \rangle\rangle &\approx \frac{\eta^2}{\sigma_{\mathrm{WP}}^4} \left\langle \left[ \left( \sum_{m=1}^{M}\sum_{l=1}^{N} \frac{\partial E}{\partial w_{ml}} \xi_{ml}^{\mathrm{WP}} + \tilde{E}_{ij} \right) \xi_{ij}^{\mathrm{WP}} \right]^2 \right\rangle - \eta^2 \left( \frac{\partial E}{\partial w_{ij}} \right)^2 \\ &= \eta^2 \sum_{m=1}^{M}\sum_{l=1}^{N} \left( \frac{\partial E}{\partial w_{ml}} \right)^2 + \eta^2 \left( \frac{\partial E}{\partial w_{ij}} \right)^2 + \frac{\eta^2}{\sigma_{\mathrm{WP}}^2} \tilde{E}_{ij}^2. \end{aligned} \tag{S5}$$

To show that the choice of $E$ minimizes the update noise for NP, we analogously compute the variance $\langle\langle \Delta w_{ij}^{\mathrm{NP}} \rangle\rangle$ of the NP weight updates Eq. (6) when adding a possibly trial- and neuron-dependent baseline term $\tilde{E}_i$, $\Delta w_{ij}^{\mathrm{NP}} = -\frac{\eta}{\sigma_{\mathrm{NP}}^2}(E^{\mathrm{pert}} - E + \tilde{E}_i) \sum_{t=1}^{T} \xi_{it}^{\mathrm{NP}} r_{jt}$,

$$\begin{aligned} \langle\langle \Delta w_{ij}^{\mathrm{NP}} \rangle\rangle &\approx \frac{\eta^2}{\sigma_{\mathrm{NP}}^4} \left\langle \left[ \left( \sum_{m=1}^{M}\sum_{s=1}^{T} \frac{\partial E}{\partial y_{ms}} \xi_{ms}^{\mathrm{NP}} + \tilde{E}_i \right) \sum_{t=1}^{T} \xi_{it}^{\mathrm{NP}} r_{jt} \right]^2 \right\rangle - \eta^2 \left( \frac{\partial E}{\partial w_{ij}} \right)^2 \\ &= \eta^2 \sum_{m=1}^{M}\sum_{s=1}^{T} \left( \frac{\partial E}{\partial y_{ms}} \right)^2 \sum_{t=1}^{T} r_{jt}^2 + \eta^2 \left( \frac{\partial E}{\partial w_{ij}} \right)^2 + \frac{\eta^2}{\sigma_{\mathrm{NP}}^2} \tilde{E}_i^2 \sum_t r_{jt}^2. \end{aligned} \tag{S6}$$

For both algorithms the variance is minimal if $\tilde{E} = 0$, as apparent from its quadratic occurrence with positive prefactor.

#### Task setting

We train a linear readout $w \in \mathbb{R}^{M\times N}$ that maps $N$ input traces $r \in \mathbb{R}^{N\times T}$ of time-dimension $T$ onto $M$ output traces $z \in \mathbb{R}^{M\times T}$ of the same time-dimension. If not stated otherwise, input and target traces do not change between trials. The readout weights $w_{ij}$ are trained to minimize the mean squared deviation $E = \frac{1}{2T}\sum_{i=1}^{M}\sum_{t=1}^{T}(z_{it} - z^*_{it})^2$ of the output $z$ from a target $z^*$. The target reads in general $z^*_{it} = \sum_{j=1}^{N} w^*_{ij} r_{jt} + d_{it}$, where $w^*_{ij}$ are target weights - i.e. the output error is certainly minimal for $w_{ij} = w^*_{ij}$ - and $d_{it}$ is an output component that cannot be generated by the network, since it is orthogonal to its inputs, $\sum_{t=1}^{T} d_{it} r_{jt} = 0\ \forall i, j$. It is useful to define the (symmetric, positive semi-definite) input correlation matrix

$$S_{ij} = \frac{1}{T}\sum_{t=1}^{T} r_{it} r_{jt}. \tag{S7}$$

$S$ can be diagonalized by a rotation $O \in \mathrm{SO}(N)$ into $D = O^T S O$ such that $D_{\mu\nu} = \alpha_\mu^2 \delta_{\mu\nu}$ is the diagonal matrix of eigenvalues $\alpha_\mu^2 \geq 0$ (where $\mu, \nu = 1, \cdots, N$). We refer to the $N$-dimensional (spatial) eigenvectors of $S$ as input directions. Another useful characteristic is the autocorrelation of the inputs. We denote the autocorrelation summed over all inputs and normalized by $T$ by

$$C_{ts} = \frac{1}{T}\sum_{j=1}^{N} r_{jt} r_{js}. \tag{S8}$$

We refer to the $T$-dimensional (temporal) eigenvectors of $C$ as input components. Reading out from an input direction yields a temporal output vector parallel to the related input component. $S$ and $C$ have the same nonzero eigenvalues $\alpha_\mu^2$. (This follows for example from the more general fact that the products $AB$ and $BA$ of an $N\times T$ matrix $A$ and a $T \times N$ matrix $B$ have the same nonzero eigenvalues ***Horn and Johnson*** (***2012***) by setting $A_{it} = r_{it}$, $B_{ti} = r_{it}$.) We call the $\alpha_\mu^2$ input "strengths", since they equal the average strength of the $\mu$th latent input per time step, or equivalently the average strength of all input activity read out from the $\mu$th input direction. The sum of the eigenvalues, the trace

$$\mathrm{tr}[S] \equiv \alpha_{\mathrm{tot}}^2, \tag{S9}$$

equals the total average input strength per time step $\alpha_{\mathrm{tot}}^2$. We call $\alpha_{\mathrm{tot}}^2$ the total input strength for short.

Throughout the following parts SM2-SM5 we will mainly consider inputs with correlation matrices $S$ that have $N_{\mathrm{eff}}$ eigenvalues equal to $\alpha^2$ and all others zero (Tab. 1), although intermediate results can also hold for inputs with general correlations $S$. The correlation matrix then has the useful properties

$$S^2 = \alpha^2 S, \tag{S10}$$

$$\mathrm{tr}[S] \equiv \alpha_{\mathrm{tot}}^2 = \alpha^2 N_{\mathrm{eff}}. \tag{S11}$$

Varying the effective input dimensionality while keeping the total input strength $\alpha_{\mathrm{tot}}^2$ constant thus implies that the strengths of individual input components scale like $\alpha^2 \sim N_{\mathrm{eff}}^{-1}$.

To measure the strength of the unrealizable target component, we define a quantity $\alpha_d^2$ analogous to $\alpha^2$. Since $\alpha^2 = \frac{1}{N_{\mathrm{eff}}}\mathrm{tr}[S] = \frac{1}{N_{\mathrm{eff}} T}\mathrm{tr}[rr^T]$ is, in particular, the average input strength per time and latent input, we set

$$\alpha_d^2 \equiv \frac{1}{MT}\mathrm{tr}[dd^T]; \tag{S12}$$

$\alpha_d^2$ is the average strength of the unrealizable component of the target per time and output.

Using the weight mismatch $W_{ij} = w_{ij} - w^*_{ij}$ and the correlation matrix $S$, the error function can be written as

$$E = \frac{1}{2T}\sum_{i=1}^{M}\sum_{t=1}^{T}(z_{it} - z^*_{it})^2 \tag{S13}$$

$$= \frac{1}{2T}\sum_{i=1}^{M}\sum_{t=1}^{T}\Big(\sum_{j=1}^{N} W_{ij} r_{jt} - d_{it}\Big)^2 = \tfrac{1}{2}\mathrm{tr}[WSW^T] + E_{\mathrm{opt}}. \tag{S14}$$

Here $E_{\mathrm{opt}}$ is the lowest achievable error corresponding to zero weight mismatch,

$$E_{\mathrm{opt}} \equiv \frac{1}{2T}\mathrm{tr}[dd^T] = \tfrac{1}{2}M\alpha_d^2. \tag{S15}$$

We note that due to the division by $T$ both the correlation matrix and the error no longer scale with the task duration, but the error does scale with the number $M$ of outputs.

The choice of a quadratic error function allows to compute the evolution of the expected error analytically. For the quadratic error function Eq. (8,S13), the average WP and NP weight updates Eq. (3) and Eq. (6) follow the gradient exactly, i.e. the relations Eqs. (4,S2,S4) hold exactly for any perturbation size:

$$\begin{aligned}
\langle \Delta w_{ij}^{\mathrm{WP}} \rangle &= -\frac{\eta}{\sigma_{\mathrm{WP}}^2} \langle (E^{\mathrm{pert}} - E) \xi_{ij}^{\mathrm{WP}} \rangle \\
&= -\frac{\eta}{\sigma_{\mathrm{WP}}^2} \sum_{m=1}^{M} \sum_{k,l=1}^{N} \left( W_{mk} S_{kl} \langle \xi_{ml}^{\mathrm{WP}} \xi_{ij}^{\mathrm{WP}} \rangle + \frac{1}{2} S_{kl} \cancel{\langle \xi_{mk}^{\mathrm{WP}} \xi_{ml}^{\mathrm{WP}} \xi_{ij}^{\mathrm{WP}} \rangle} \right) \\
&= -\eta \sum_{k=1}^{N} W_{ik} S_{kj} = -\eta \frac{\partial E}{\partial w_{ij}}
\end{aligned} \tag{S16}$$

$$\begin{aligned}
\langle \Delta w_{ij}^{\mathrm{NP}} \rangle &= -\frac{\eta}{\sigma_{\mathrm{NP}}^2} \sum_{t=1}^{T} \langle (E^{\mathrm{pert}} - E) \xi_{it}^{\mathrm{NP}} \rangle r_{jt} \\
&= -\frac{\eta}{\sigma_{\mathrm{WP}}^2 T} \sum_{m=1}^{M} \sum_{s,t=1}^{T} \left( \sum_{k=1}^{N} W_{mk} r_{ks} \langle \xi_{ms}^{\mathrm{NP}} \xi_{it}^{\mathrm{NP}} \rangle + \frac{1}{2} \cancel{\langle (\xi_{ms}^{\mathrm{NP}})^2 \xi_{it}^{\mathrm{NP}} \rangle} - d_{ms} \langle \xi_{ms}^{\mathrm{NP}} \xi_{it}^{\mathrm{NP}} \rangle \right) r_{jt} \\
&= -\eta \sum_{k=1}^{N} W_{ik} S_{kj} + \frac{\eta}{T} \cancel{\sum_{t=1}^{T} d_{it} r_{jt}} = -\eta \frac{\partial E}{\partial w_{ij}}
\end{aligned} \tag{S17}$$

#### Effective perturbation strength

For a fair comparison of WP and NP we consider the output perturbations $\delta z = z^{\mathrm{pert}} - z$ that they generate. For WP Eqs. (2,7) imply

$$\delta z_{it} = \sum_{j=1}^{N} \xi_{ij}^{\mathrm{WP}} r_{jt}, \tag{S18}$$

for NP Eqs. (5,7) imply

$$\delta z_{it} = \xi_{it}^{\mathrm{NP}}. \tag{S19}$$

We choose $\sigma_{\mathrm{WP}}$ and $\sigma_{\mathrm{NP}}$ such that weight and node perturbations lead to the same output perturbation strength as measured by $\sigma_{\mathrm{eff}}^2 = \frac{1}{MT} \langle \sum_{it} \left(\delta z_{it}\right)^2 \rangle$, the total induced output variance per time step and output neuron:

$$\sigma_{\mathrm{eff,WP}}^2 = \frac{1}{MT} \sum_{i=1}^{M} \sum_{t=1}^{T} \left\langle \left( \sum_{j=1}^{N} \xi_{ij}^{\mathrm{WP}} r_{jt} \right)^2 \right\rangle = \sigma_{\mathrm{WP}}^2 \cdot \alpha^2 N_{\mathrm{eff}}, \tag{S20}$$

$$\sigma_{\mathrm{eff,NP}}^2 = \frac{1}{MT} \sum_{i=1}^{M} \sum_{t=1}^{T} \langle \left(\xi_{it}^{\mathrm{NP}}\right)^2 \rangle = \sigma_{\mathrm{NP}}^2. \tag{S21}$$

Here we used $\langle \xi_{ij}^{\mathrm{WP}} \xi_{mk}^{\mathrm{WP}} \rangle = \sigma_{\mathrm{WP}}^2 \delta_{im} \delta_{jk}$ and $\langle \xi_{it}^{\mathrm{NP}} \xi_{ms}^{\mathrm{NP}} \rangle = \sigma_{\mathrm{NP}}^2 \delta_{im} \delta_{ts}$. Requiring $\sigma_{\mathrm{eff,WP}}^2 \overset{!}{=} \sigma_{\mathrm{eff,NP}}^2 \equiv \sigma_{\mathrm{eff}}^2$ implies

$$\sigma_{\mathrm{NP}}^2 = \sigma_{\mathrm{eff}}^2 \qquad \sigma_{\mathrm{WP}}^2 = \frac{1}{\alpha^2 N_{\mathrm{eff}}} \cdot \sigma_{\mathrm{eff}}^2. \tag{S22}$$

We note that although the induced output perturbations have the same variance, they follow different distributions,

$$\frac{1}{MT} \sum_{it} \left(\delta z_{it}\right)^2 \sim \begin{cases} \frac{\sigma_{\mathrm{eff}}^2}{M N_{\mathrm{eff}}} \chi_{k=M N_{\mathrm{eff}}}^2 & \text{for WP,} \\ \frac{\sigma_{\mathrm{eff}}^2}{MT} \chi_{k=MT}^2 & \text{for NP,} \end{cases} \tag{S23}$$

where $\chi_k^2$ is the chi-square distribution for $k$ degrees of freedom.

## SM 2

### Derivation of error dynamics

This part starts with a derivation of the error dynamics if our tasks are learned with pure gradient descent (GD) learning. These are comparably simple and a useful benchmark. The sections thereafter provide a full derivation of the error dynamics of WP and NP learning, main text, Eqs. (9-14). Their analysis and interpretation follow in SM3.

##### Error curves for gradient descent

In GD the updates directly follow the gradient,

$$\Delta w_{ij}^{\mathrm{GD}} = -\eta \frac{\partial E}{\partial w_{ij}} = -\eta \sum_{k=1}^{N} W_{ik} S_{kj}. \tag{S24}$$

The error after such a deterministic update (which equals the expected error) is

$$\begin{aligned} E(n) &= \tfrac{1}{2}\,\mathrm{tr}[(W(n-1)+\Delta w^{\mathrm{GD}})\tilde{S}(W(n-1)+\Delta w^{\mathrm{GD}})^T] + E_{\mathrm{opt}} \\ &= E(n-1) + \mathrm{tr}[W(n-1)\tilde{S}(\Delta w^{\mathrm{GD}})^T] + \tfrac{1}{2}\,\mathrm{tr}[\Delta w^{\mathrm{GD}}\tilde{S}(\Delta w^{\mathrm{GD}})^T] \\ &= E(n-1) - \eta\,\mathrm{tr}[(W\tilde{S}SW^T] + \tfrac{1}{2}\eta^2\,\mathrm{tr}[WS\tilde{S}SW^T], \end{aligned} \tag{S25}$$

where $W$ stands for $W(n-1)$. To facilitate the tracing of the different terms, we have tagged the correlation matrix that stems from the cost evaluation at trial $n$ by a tilde, which keeps it distinguishable from those arising from the weight update Eq. (S24); the entries $\tilde{S}_{ij}$ are identical to $S_{ij}$. For a correlation matrix $S$ that has $N_{\mathrm{eff}}$ eigenvalues equal to $\alpha^2$ and all others zero, see SM1, Sec. "Task setting", we can use Eq. (S10) (and $\tilde{S} \equiv S$) together with $\frac{1}{2}\,\mathrm{tr}[WSW^T] = E(n-1) - E_{\mathrm{opt}}$ to obtain

$$\begin{aligned} E(n) &= E(n-1) - \eta\alpha^2\,\mathrm{tr}[(WSW^T] + \tfrac{1}{2}\eta^2\alpha^4\,\mathrm{tr}[WSW^T] \\ &= (1-2\eta\alpha^2+\eta^2\alpha^4)\cdot(E(n-1)-E_{\mathrm{opt}}) + E_{\mathrm{opt}}. \end{aligned} \tag{S26}$$

This leads to an exponential decay of the error to $E_{\mathrm{opt}}$,

$$E(n) = (E(0) - E_{\mathrm{opt}})a^n + E_{\mathrm{opt}}, \tag{S27}$$

with

$$a_{\mathrm{GD}} = 1 - 2\eta\alpha^2 + \eta^2\alpha^4, \tag{S28}$$

which becomes zero for the optimal learning rate $\eta^* = \alpha^{-2}$ such that the task is solved in a single trial. As for WP and NP (Eqs. (11,S62)), learning diverges once $\eta > 2\eta^*$ where $a > 1$.

##### Error Curves for Weight Perturbation

Since we only consider WP here, for clarity of notation we omit the specifier "WP" in $\xi^{\mathrm{WP}}$, $\Delta w^{\mathrm{WP}}$ and $\sigma_{\mathrm{WP}}$. The error $E^{\mathrm{pert}}$ as a function of the perturbations $\xi$ applied to the weights then reads

$$E^{\mathrm{pert}} = \tfrac{1}{2}\,\mathrm{tr}[(W+\xi)S(W+\xi)^T] + E_{\mathrm{opt}} = E + \mathrm{tr}[WS\xi^T] + \tfrac{1}{2}\,\mathrm{tr}[\xi S\xi^T], \tag{S29}$$

which yields a weight update (cf. Eq. (3))

$$\begin{aligned} \Delta w_{ij} &= -\frac{\eta}{\sigma^2}(E^{\mathrm{pert}} - E)\xi_{ij} \\ &= -\frac{\eta}{\sigma^2}\left(\mathrm{tr}[WS\xi^T] + \tfrac{1}{2}\,\mathrm{tr}[\xi S\xi^T]\right)\xi_{ij}. \end{aligned} \tag{S30}$$

Averaging over $\xi$ gives the expected error $\langle E(n)\rangle$ after the $n$th update,

$$\begin{aligned} \langle E(n)\rangle &= \tfrac{1}{2}\,\langle \mathrm{tr}[(W(n-1)+\Delta w)\tilde{S}(W(n-1)+\Delta w)^T]\rangle + E_{\mathrm{opt}} \\ &= \langle E(n-1)\rangle + \underbrace{\langle \mathrm{tr}[W\tilde{S}\Delta w^T]\rangle}_{\equiv\,\Delta E^{\mathrm{lin}}_{\mathrm{upd}}} + \underbrace{\tfrac{1}{2}\,\langle \mathrm{tr}[\Delta w\tilde{S}\Delta w^T]\rangle}_{\equiv\,\Delta E^{\mathrm{quad}}_{\mathrm{upd}}}, \end{aligned} \tag{S31}$$

where $W$ stands for $W(n-1)$. Here we again tagged the correlation matrix with a tilde, since it stems from the cost evaluation at trial $n$ (cf. Sec. "Error curves for gradient descent"). Using Eq. (S30) and that odd moments of $\xi$ vanish, the first highlighted term, $\Delta E_{\text{upd}}^{\text{lin}}$, which is linear in the update, gives

$$\begin{aligned}
\Delta E_{\text{upd}}^{\text{lin}} &= \langle \operatorname{tr}[W\tilde{S}\Delta w^T]\rangle \\
&= -\frac{\eta}{\sigma^2}\langle \operatorname{tr}[W\tilde{S}\big(\operatorname{tr}[WS\xi^T] + \tfrac{1}{2}\cancel{\operatorname{tr}[\xi S\xi^T]}\big)\xi^T]\rangle \\
&= -\frac{\eta}{\sigma^2}\langle \operatorname{tr}[W\tilde{S}\xi^T]\operatorname{tr}[WS\xi^T]\rangle \\
&= -\frac{\eta}{\sigma^2}\sum_{im=1}^{M}\sum_{jklp=1}^{N} W_{ij}\tilde{S}_{jk}W_{ml}S_{lp}\cdot\langle\xi_{ik}\xi_{mp}\rangle\,.
\end{aligned} \tag{S32}$$

Using again Eq. (S30) and that odd moments of $\xi$ vanish, the second highlighted term, $\Delta E_{\text{upd}}^{\text{quad}}$, which is quadratic in the updates, can be expanded as

$$\begin{aligned}
\Delta E_{\text{upd}}^{\text{quad}} &= \tfrac{1}{2}\langle \operatorname{tr}[\Delta w\tilde{S}\Delta w^T]\rangle \\
&= \frac{\eta^2}{2\sigma^4}\langle\big(\operatorname{tr}[WS\xi^T] + \tfrac{1}{2}\operatorname{tr}[\xi S\xi^T]\big)\cdot\operatorname{tr}[\xi\tilde{S}\xi^T]\cdot\big(\operatorname{tr}[WS\xi^T] + \tfrac{1}{2}\operatorname{tr}[\xi S\xi^T]\big)\rangle \\
&= \frac{\eta^2}{2\sigma^4}\langle\operatorname{tr}[\xi\tilde{S}\xi^T]\operatorname{tr}[WS\xi^T]^2\rangle + \frac{\eta^2}{8\sigma^4}\langle\operatorname{tr}[\xi\tilde{S}\xi^T]\operatorname{tr}[\xi S\xi^T]^2\rangle \\
&= \underbrace{\frac{\eta^2}{2\sigma^4}\sum_{imn=1}^{M}\sum_{jklpqr=1}^{N}\tilde{S}_{jk}W_{ml}S_{lp}W_{nq}S_{qr}\cdot\langle\xi_{ij}\xi_{ik}\xi_{mp}\xi_{nr}\rangle}_{\equiv\Delta E_{\text{upd}}^{\text{quad,a}}} + \underbrace{\frac{\eta^2}{8\sigma^4}\sum_{imn=1}^{M}\sum_{jklpqr=1}^{N}\tilde{S}_{jk}S_{lp}S_{qr}\cdot\langle\xi_{ij}\xi_{ik}\xi_{ml}\xi_{mp}\xi_{nq}\xi_{nr}\rangle}_{\equiv\Delta E_{\text{upd}}^{\text{quad,b}}}\,.
\end{aligned} \tag{S33}$$

$\Delta E_{\text{upd}}^{\text{quad,a}}$ depends on the weight mismatch $W$ and originates from the gradient related part of the error signal $\Delta E_{\text{pert}} - E$, whereas $\Delta E_{\text{upd}}^{\text{quad,b}}$ originates from quadratic reward noise (SM3). The moments of $\xi$ can be computed using $\langle\xi_{ij}\xi_{mk}\rangle = \sigma^2\delta_{im}\delta_{jk}$ and Isserlis' theorem,

$$\langle\xi_{ik}\xi_{mp}\rangle = \sigma^2\delta_{im}\delta_{kp} \tag{S34}$$

$$\langle\xi_{ij}\xi_{ik}\xi_{mp}\xi_{nr}\rangle = \sigma^4\big(\delta_{jk}\delta_{mn}\delta_{pr} + \delta_{im}\delta_{jp}\delta_{in}\delta_{kr} + \delta_{in}\delta_{jr}\delta_{im}\delta_{kp}\big) \tag{S35}$$

$$\begin{aligned}
\langle\xi_{ij}\xi_{ik}\xi_{ml}\xi_{mp}\xi_{nq}\xi_{nr}\rangle = \sigma^6\big(&\delta_{jk}(\delta_{lp}\delta_{qr} + \delta_{mn}\delta_{lq}\delta_{pr} + \delta_{mn}\delta_{lr}\delta_{pq}) && (= \langle\xi_{ij}\xi_{ik}\rangle\langle\xi_{ml}\xi_{mp}\xi_{nq}\xi_{nr}\rangle)\\
&+ \delta_{im}\delta_{jl}(\delta_{kp}\delta_{qr} + \delta_{in}\delta_{kq}\delta_{pr} + \delta_{in}\delta_{kr}\delta_{pq}) && (= \langle\xi_{ij}\xi_{ml}\rangle\langle\xi_{ik}\xi_{mp}\xi_{nq}\xi_{nr}\rangle)\\
&+ \delta_{im}\delta_{jp}(\delta_{kl}\delta_{qr} + \delta_{in}\delta_{kq}\delta_{lr} + \delta_{in}\delta_{kr}\delta_{lq}) && (= \langle\xi_{ij}\xi_{mp}\rangle\langle\xi_{ik}\xi_{ml}\xi_{nq}\xi_{nr}\rangle)\\
&+ \delta_{in}\delta_{jq}(\delta_{im}\delta_{kl}\delta_{pr} + \delta_{im}\delta_{kp}\delta_{lr} + \delta_{kr}\delta_{lp}) && (= \langle\xi_{ij}\xi_{nq}\rangle\langle\xi_{ik}\xi_{ml}\xi_{mp}\xi_{nr}\rangle)\\
&+ \delta_{in}\delta_{jr}(\delta_{im}\delta_{kl}\delta_{pq} + \delta_{im}\delta_{kp}\delta_{lq} + \delta_{kq}\delta_{lp})\big). && (= \langle\xi_{ij}\xi_{nr}\rangle\langle\xi_{ik}\xi_{ml}\xi_{mp}\xi_{nq}\rangle)
\end{aligned} \tag{S36}$$

Inserting this into Eqs. (S32,S33) and performing the summations is partially lengthy but straightforward. It results in

$$\Delta E_{\text{upd}}^{\text{lin}} = -\eta\sum_{i=1}^{M}\sum_{jkl=1}^{N}W_{ij}\tilde{S}_{jk}S_{lk}W_{il} = -\eta\operatorname{tr}[W\tilde{S}SW^T], \tag{S37}$$

$$\begin{aligned}
\Delta E_{\text{upd}}^{\text{quad,a}} &= \frac{\eta^2}{2}\sum_{imn=1}^{M}\sum_{jklpqr}^{N}\tilde{S}_{jk}W_{ml}S_{lp}W_{nq}S_{qr}\cdot\big(\delta_{jk}\delta_{mn}\delta_{pr} + \delta_{im}\delta_{jp}\delta_{in}\delta_{kr} + \delta_{in}\delta_{jr}\delta_{im}\delta_{kp}\big)\\
&= \frac{\eta^2}{2}M\operatorname{tr}[WS^2W^T]\operatorname{tr}[\tilde{S}] + \eta^2\operatorname{tr}[WS\tilde{S}SW^T],
\end{aligned} \tag{S38}$$

$$\Delta E_{\text{upd}}^{\text{quad,b}} = \frac{\eta^2\sigma^2}{8}\big(M^3\operatorname{tr}[\tilde{S}]\operatorname{tr}[S]^2 + 2M^2\operatorname{tr}[\tilde{S}]\operatorname{tr}[S^2] + 4M^2\operatorname{tr}[\tilde{S}S]\operatorname{tr}[S] + 8M\operatorname{tr}[S\tilde{S}S]\big). \tag{S39}$$

With this, Eq. (S31) for the evolution of expected error becomes

$$\begin{aligned}
\langle E(n)\rangle &\overset{\text{WP}}{=} \langle E(n-1)\rangle + \Delta E_{\text{upd}}^{\text{lin}} + \Delta E_{\text{upd}}^{\text{quad,a}} + \Delta E_{\text{upd}}^{\text{quad,b}}\\
&= \langle E(n-1)\rangle - \eta\operatorname{tr}[W\tilde{S}SW^T] + \frac{\eta^2}{2}M\operatorname{tr}[WS^2W^T]\operatorname{tr}[\tilde{S}] + \eta^2\operatorname{tr}[WS\tilde{S}SW^T]\\
&\quad + \frac{\eta^2\sigma^2}{8}\big(M^3\operatorname{tr}[\tilde{S}]\operatorname{tr}[S]^2 + 2M^2\operatorname{tr}[\tilde{S}]\operatorname{tr}[S^2] + 4M^2\operatorname{tr}[\tilde{S}S]\operatorname{tr}[S] + 8M\operatorname{tr}[S\tilde{S}S]\big).
\end{aligned} \tag{S40}$$

The result holds for a general input correlation matrix $S$. In Sec. "Error curves for equally strong input components" we will assume same strength latent inputs (Tab. 1). The resulting properties of $S$ (Eqs. (S10,S11)) allow to re-express all right hand side terms of Eq. (S40) in terms of $E(n-1)$ instead of $W$, yielding a scalar recurrence relation. We will make the assumption of equally strong latent inputs also in parts SM3-SM5 (Tab. 1). The convergence for general input will be analyzed in SM6.

#### Error curves for node perturbation

Similar to the previous section, we omit the specifier "NP" in $\xi^{\mathrm{NP}}$, $\Delta w^{\mathrm{NP}}$ and $\sigma_{\mathrm{NP}}$ in the following for notational clarity. The error $E^{\mathrm{pert}}$ then reads as a function of the NP $\xi$

$$\begin{aligned} E^{\mathrm{pert}} &= \frac{1}{2T}\sum_{i=1}^{M}\sum_{t=1}^{T}\Big(\sum_{j=1}^{N} W_{ij}r_{jt} + \xi_{it} - d_{it}\Big)^2 \\ &= \frac{1}{2T}\sum_{i=1}^{M}\sum_{t=1}^{T}\Big(\Big(\sum_{j=1}^{N} W_{ij}r_{jt} + \xi_{it}\Big)^2 - 2\sum_{j=1}^{N} W_{ij}\cancel{r_{jt}d_{it}} - 2\xi_{it}d_{it} + d_{it}^2\Big) \\ &= E + \tfrac{1}{T}\,\mathrm{tr}[Wr\xi^T] + \tfrac{1}{2T}\,\mathrm{tr}[\xi\xi^T] - \tfrac{1}{T}\,\mathrm{tr}[d\xi^T]. \end{aligned} \tag{S41}$$

The term $\mathrm{tr}[d\xi^T]$ here reflects the interference of learning and unrealizable targets. The other term including $d$, $\frac{1}{2T}\,\mathrm{tr}[dd^T] = E_{\mathrm{opt}}$, is only an additive constant which also occurs in GD and WP and does not enter the update equation. Eq. (S41) yields a weight update (cf. Eq. (6))

$$\begin{aligned} \Delta w_{ij} &= -\frac{\eta}{\sigma^2}(E^{\mathrm{pert}} - E)\sum_{t=1}^{T}\xi_{it}r_{jt} \\ &= -\frac{\eta}{\sigma^2 T}\Big(\mathrm{tr}[Wr\xi^T] + \tfrac{1}{2}\,\mathrm{tr}[\xi\xi^T] - \mathrm{tr}[d\xi^T]\Big)\sum_{t=1}^{T}\xi_{it}r_{jt}. \end{aligned} \tag{S42}$$

Averaging over $\xi$ gives the expected error after the update,

$$\begin{aligned} \langle E(n)\rangle &= \tfrac{1}{2}\,\langle \mathrm{tr}[(W(n-1)+\Delta w)\tilde{S}(W(n-1)+\Delta w)^T]\rangle + E_{\mathrm{opt}} \\ &= \langle E(n-1)\rangle + \underbrace{\langle \mathrm{tr}[W\tilde{S}\Delta w^T]\rangle}_{\equiv\,\Delta E_{\mathrm{upd}}^{\mathrm{lin}}} + \underbrace{\tfrac{1}{2}\,\langle \mathrm{tr}[\Delta w\tilde{S}\Delta w^T]\rangle}_{\equiv\,\Delta E_{\mathrm{upd}}^{\mathrm{quad}}}, \end{aligned} \tag{S43}$$

where $W$ stands for $W(n-1)$ and the correlation matrix that stems from the error evaluation is tagged with a tilde. Using Eq. (S42) and that odd moments of $\xi$ vanish, $\Delta E_{\mathrm{upd}}^{\mathrm{lin}}$ and $\Delta E_{\mathrm{upd}}^{\mathrm{quad}}$ are

$$\begin{aligned} \Delta E_{\mathrm{upd}}^{\mathrm{lin}} &= \langle \mathrm{tr}[W\tilde{S}\Delta w^T]\rangle \\ &= -\frac{\eta}{\sigma^2 T}\,\langle \mathrm{tr}[W\tilde{S}\big(\mathrm{tr}[Wr\xi^T] + \tfrac{1}{2}\cancel{\mathrm{tr}[\xi\xi^T]} - \mathrm{tr}[d\xi^T]\big)r\xi^T]\rangle \\ &= -\frac{\eta}{\sigma^2 T}\,\langle \mathrm{tr}[W\tilde{S}r\xi^T]\,\mathrm{tr}[Wr\xi^T] - \mathrm{tr}[W\tilde{S}r\xi^T]\,\mathrm{tr}[d\xi^T]\rangle \\ &= -\frac{\eta}{\sigma^2 T}\sum_{im=1}^{M}\sum_{jkl=1}^{N}\sum_{st=1}^{T} W_{ij}\tilde{S}_{jk}W_{ml}r_{lt}r_{ks}\cdot\langle\xi_{is}\xi_{mt}\rangle + \frac{\eta}{\sigma^2 T}\sum_{im=1}^{M}\sum_{jk=1}^{N}\sum_{st=1}^{T} W_{ij}\tilde{S}_{jk}r_{ks}d_{mt}\cdot\langle\xi_{is}\xi_{mt}\rangle, \end{aligned} \tag{S44}$$

$$\begin{aligned} \Delta E_{\mathrm{upd}}^{\mathrm{quad}} &= \tfrac{1}{2}\,\langle \mathrm{tr}[\Delta w\tilde{S}\Delta w^T]\rangle \\ &= \frac{\eta^2}{2\sigma^4 T^2}\,\langle \mathrm{tr}[\xi r^T\tilde{S}r\xi^T]\big(\mathrm{tr}[Wr\xi^T]^2 + \tfrac{1}{4}\,\mathrm{tr}[\xi\xi^T]^2 + \mathrm{tr}[d\xi^T]^2 - 2\,\mathrm{tr}[Wr\xi^T]\,\mathrm{tr}[d\xi^T]\big)\rangle \\ &= \frac{\eta^2}{2\sigma^4 T^2}\sum_{imn=1}^{M}\sum_{jklp=1}^{N}\sum_{stuv=1}^{T} r_{js}\tilde{S}_{jk}r_{kt}W_{ml}r_{lu}W_{np}r_{pv}\cdot\langle\xi_{is}\xi_{it}\xi_{mu}\xi_{nv}\rangle \qquad \big(\Delta E_{\mathrm{upd}}^{\mathrm{quad,a}}\big) \qquad \text{(S45)} \\ &\quad + \frac{\eta^2}{8\sigma^4 T^2}\sum_{imn=1}^{M}\sum_{jk=1}^{N}\sum_{stuv=1}^{T} r_{js}\tilde{S}_{jk}r_{kt}\cdot\langle\xi_{is}\xi_{it}\xi_{mu}\xi_{mu}\xi_{nv}\xi_{nv}\rangle \qquad \big(\Delta E_{\mathrm{upd}}^{\mathrm{quad,b}}\big) \qquad \text{(S46)} \\ &\quad + \frac{\eta^2}{2\sigma^4 T^2}\sum_{imn=1}^{M}\sum_{jk=1}^{N}\sum_{stuv=1}^{T} r_{js}\tilde{S}_{jk}r_{kt}d_{mu}d_{nv}\cdot\langle\xi_{is}\xi_{it}\xi_{mu}\xi_{nv}\rangle \qquad \big(\Delta E_{\mathrm{upd}}^{\mathrm{quad,c}}\big) \qquad \text{(S47)} \\ &\quad - \frac{\eta^2}{\sigma^4 T^2}\sum_{imn=1}^{M}\sum_{jkl=1}^{N}\sum_{stuv=1}^{T} r_{js}\tilde{S}_{jk}r_{kt}W_{ml}r_{lu}d_{nv}\cdot\langle\xi_{is}\xi_{it}\xi_{mu}\xi_{nv}\rangle \qquad \big(\Delta E_{\mathrm{upd}}^{\mathrm{quad,d}}\big). \qquad \text{(S48)} \end{aligned}$$

Like for WP, $\Delta E_{\text{upd}}^{\text{quad,a}}$ depends on the weight mismatch $W$ and originates from the gradient related part of the error signal $\Delta E_{\text{pert}} - E$, whereas $\Delta E_{\text{upd}}^{\text{quad,b}}$ originates from quadratic reward noise (SM3). $\Delta E_{\text{upd}}^{\text{quad,c}}$ and $\Delta E_{\text{upd}}^{\text{quad,d}}$ stem from the reward noise due to coupling to unrealizable target components. The moments of $\xi$ can again be computed from $\langle \xi_{is}\xi_{mt}\rangle = \sigma^2\delta_{im}\delta_{st}$ and Isserlis' theorem,

$$\langle \xi_{is}\xi_{mt}\rangle = \sigma^2\delta_{im}\delta_{st} \tag{S49}$$

$$\langle \xi_{is}\xi_{it}\xi_{mu}\xi_{nv}\rangle = \sigma^4\left(\delta_{st}\delta_{mn}\delta_{uv} + \delta_{im}\delta_{su}\delta_{in}\delta_{tv} + \delta_{in}\delta_{sv}\delta_{im}\delta_{tu}\right) \tag{S50}$$

$$\begin{aligned}\langle \xi_{is}\xi_{it}\xi_{mu}\xi_{mu}\xi_{nv}\xi_{nv}\rangle = \sigma^6\big(&\delta_{st}\left(1 + 2\delta_{mn}\delta_{uv}\right) && (= \langle \xi_{is}\xi_{it}\rangle\langle \xi_{mu}\xi_{mu}\xi_{nv}\xi_{nv}\rangle)\\ &+ 2\delta_{im}\delta_{su}\left(\delta_{im}\delta_{tu} + 2\delta_{in}\delta_{tv}\delta_{mn}\delta_{uv}\right) && (= 2\langle \xi_{is}\xi_{mu}\rangle\langle \xi_{it}\xi_{mu}\xi_{nv}\xi_{nv}\rangle)\\ &+ 2\delta_{in}\delta_{sv}\left(\delta_{in}\delta_{tv} + 2\delta_{im}\delta_{tu}\delta_{mn}\delta_{uv}\right)\big) && (= 2\langle \xi_{is}\xi_{nv}\rangle\langle \xi_{it}\xi_{mu}\xi_{mu}\xi_{nv}\rangle)\\ = \sigma^6\big(&\delta_{st}\left(1 + 2\delta_{mn}\delta_{uv}\right) + 8\delta_{imn}\delta_{stuv}\\ &+ 2\delta_{im}\delta_{su}\delta_{tu} + 2\delta_{in}\delta_{sv}\delta_{tv}\big).\end{aligned} \tag{S51}$$

Inserting this into Eqs. (S44–S48) gives after a partially lengthy but straightforward computation

$$\Delta E_{\text{upd}}^{\text{lin}} = -\eta\,\mathrm{tr}[W\tilde{S}SW^T], \tag{S52}$$

$$\Delta E_{\text{upd}}^{\text{quad,a}} = \frac{\eta^2}{2}M\,\mathrm{tr}[WSW^T]\,\mathrm{tr}[\tilde{S}S] + \eta^2\,\mathrm{tr}[WS\tilde{S}SW^T], \tag{S53}$$

$$\Delta E_{\text{upd}}^{\text{quad,b}} = \frac{\eta^2\sigma^2}{8T}\,\mathrm{tr}[\tilde{S}S]\cdot\left(M^3T^2 + 6M^2T + 8M\right), \tag{S54}$$

$$\Delta E_{\text{upd}}^{\text{quad,c}} = \frac{\eta^2}{2T}M\,\mathrm{tr}[\tilde{S}S]\,\mathrm{tr}[dd^T], \tag{S55}$$

$$\Delta E_{\text{upd}}^{\text{quad,d}} = 0. \tag{S56}$$

With this, Eq. (S43) for the evolution of expected reward becomes

$$\begin{aligned}\langle E(n)\rangle &\overset{\text{NP}}{=} \langle E(n-1)\rangle + \Delta E_{\text{upd}}^{\text{lin}} + \Delta E_{\text{upd}}^{\text{quad,a}} + \Delta E_{\text{upd}}^{\text{quad,b}} + \Delta E_{\text{upd}}^{\text{quad,c}}\\ &= \langle E(n-1)\rangle - \eta\,\mathrm{tr}[W\tilde{S}SW^T] + \frac{\eta^2}{2}M\,\mathrm{tr}[WSW^T]\,\mathrm{tr}[\tilde{S}S] + \eta^2\,\mathrm{tr}[WS\tilde{S}SW^T]\\ &\quad + \frac{\eta^2\sigma_{\text{NP}}^2}{8T}\,\mathrm{tr}[\tilde{S}S]\cdot\left(M^3T^2 + 6M^2T + 8M\right) + \eta^2M\,\mathrm{tr}[\tilde{S}S]\cdot\frac{1}{2T}\,\mathrm{tr}[dd^T].\end{aligned} \tag{S57}$$

As for WP, this result holds for any correlation matrix $S$.

#### Error curves for equally strong input components

In this section we consider the case that there are $N_{\text{eff}}$ latent inputs of equal strength, see SM1, Sec. "Task setting". Using Eqs. (S10,S11) and $\tilde{S} = S$, we first simplify the evolution equation of WP's expected error, Eq. (S40). By identifying occurrences of $\frac{1}{2}\,\mathrm{tr}[WSW^T]$ and replacing them with $\langle E(n-1)\rangle - E_{\text{opt}}$ (Eq. (S14)) we obtain a linear recurrence relation

$$\begin{aligned}\langle E(n)\rangle &\overset{\text{WP}}{=} \langle E(n-1)\rangle - \eta\,\mathrm{tr}[W\tilde{S}SW^T] + \frac{\eta^2}{2}M\,\mathrm{tr}[WS^2W^T]\,\mathrm{tr}[\tilde{S}] + \eta^2\,\mathrm{tr}[WS\tilde{S}SW^T]\\ &\quad + \frac{\eta^2\sigma_{\text{WP}}^2}{8}\left(M^3\,\mathrm{tr}[\tilde{S}]\,\mathrm{tr}[S]^2 + 2M^2\,\mathrm{tr}[\tilde{S}]\,\mathrm{tr}[S^2] + 4M^2\,\mathrm{tr}[\tilde{S}S]\,\mathrm{tr}[S] + 8M\,\mathrm{tr}[S\tilde{S}S]\right)\\ &= \left(1 - 2\eta\alpha^2 + \eta^2\alpha^4(MN_{\text{eff}} + 2)\right)\cdot\left(\langle E(n-1)\rangle - E_{\text{opt}}\right)\\ &\quad + \tfrac{1}{8}\eta^2\sigma_{\text{WP}}^2\alpha^6\cdot\left(M^3N_{\text{eff}}^3 + 6M^2N_{\text{eff}}^2 + 8MN_{\text{eff}}\right) + E_{\text{opt}}.\end{aligned} \tag{S58}$$

The evolution equation of NP's expected error, Eq. (S57), similarly simplifies to a linear recurrence relation

$$\begin{aligned}\langle E(n)\rangle &\overset{\text{NP}}{=} \langle E(n-1)\rangle - \eta\,\mathrm{tr}[W\tilde{S}SW^T] + \frac{\eta^2}{2}M\,\mathrm{tr}[WSW^T]\,\mathrm{tr}[\tilde{S}S] + \eta^2\,\mathrm{tr}[WS\tilde{S}SW^T]\\ &\quad + \frac{\eta^2\sigma_{\text{NP}}^2}{8T}\,\mathrm{tr}[\tilde{S}S]\cdot\left(M^3T^2 + 6M^2T + 8M\right) + \eta^2M\,\mathrm{tr}[\tilde{S}S]\cdot\frac{1}{2T}\,\mathrm{tr}[dd^T]\\ &= \left(1 - 2\eta\alpha^2 + \eta^2\alpha^4(MN_{\text{eff}} + 2)\right)\cdot\left(\langle E(n-1)\rangle - E_{\text{opt}}\right)\\ &\quad + \tfrac{1}{8}\eta^2\sigma_{\text{NP}}^2\alpha^4\cdot\left(M^3N_{\text{eff}}T + 6M^2N_{\text{eff}} + 8M\frac{N_{\text{eff}}}{T}\right) + \eta^2\alpha^4MN_{\text{eff}}\cdot E_{\text{opt}} + E_{\text{opt}}.\end{aligned} \tag{S59}$$

Both Eq. (S58) and Eq. (S59) have the form

$$\langle E(n)\rangle = \big(\langle E(n-1)\rangle - E_{\text{opt}}\big)\cdot a + b + E_{\text{opt}}, \tag{S60}$$

with (different) constant parameters $a$ and $b$. The convergence factor $a$ characterizes the convergence speed, $b$ the per-update increase in error. The recurrence relation can be solved straightforwardly by iteration, by expanding $b$ to $\frac{b}{1-a}(1-a)$, shifting $\frac{b}{1-a}+E_{\text{opt}}$ to the equation's left hand side and considering $\langle E(n)\rangle - E_{\text{opt}} - \frac{b}{1-a}$ as recurrently specified variable,

$$\langle E(n)\rangle = \left(E(0) - \tfrac{b}{1-a} - E_{\text{opt}}\right)\cdot a^n + \frac{b}{1-a} + E_{\text{opt}}. \tag{S61}$$

Another possibility is to consider $\langle E(n)\rangle - E_{\text{opt}}$ as recurrently specified variable and to observe that the iteration gives rise to a finite geometric series that yields $\frac{1-a^n}{1-a}b$. The values of the constants are

$$a = 1 - 2\eta\alpha^2 + \eta^2\alpha^4(MN_{\text{eff}}+2), \tag{S62}$$

$$b_{\text{WP}} = \tfrac{1}{8}\eta^2\sigma_{\text{WP}}^2\alpha^6\cdot\left(M^3N_{\text{eff}}^3 + 6M^2N_{\text{eff}}^2 + 8MN_{\text{eff}}\right), \tag{S63}$$

$$b_{\text{NP}} = \tfrac{1}{8}\eta^2\sigma_{\text{NP}}^2\alpha^4\cdot\left(M^3N_{\text{eff}}T + 6M^2N_{\text{eff}} + 8M\frac{N_{\text{eff}}}{T}\right) + \eta^2\alpha^4MN_{\text{eff}}\cdot E_{\text{opt}}. \tag{S64}$$

Finite $b$ due to finite perturbation size $\sigma^2$ causes a finite residual error $b/(1-a)$ in Eq. (S61) even if $E_{\text{opt}}$ is zero. To enable a fair comparison, $\sigma_{\text{WP}}$ and $\sigma_{\text{NP}}$ are chosen such that they lead to output perturbations $\delta z$ of the same strength, see SM1, Sec. "Effective perturbation strength". Expressing $\sigma_{\text{WP}}^2$ and $\sigma_{\text{NP}}^2$ through the strength $\sigma_{\text{eff}}^2$ of the output perturbation that they generate (Eq. (S22)), the constants $b$ become

$$b_{\text{WP}} = \tfrac{1}{8}\eta^2\sigma_{\text{eff}}^2\alpha^4\cdot\left(M^3N_{\text{eff}}^2 + 6M^2N_{\text{eff}} + 8M\right), \tag{S65}$$

$$b_{\text{NP}} = \underbrace{\tfrac{1}{8}\eta^2\sigma_{\text{eff}}^2\alpha^4\cdot\left(M^3N_{\text{eff}}T + 6M^2N_{\text{eff}} + 8M\frac{N_{\text{eff}}}{T}\right)} + \underbrace{\eta^2\alpha^4MN_{\text{eff}}\cdot E_{\text{opt}}} \tag{S66}$$

$$\equiv \qquad b_{\text{NP}}^{\text{quad}} \qquad + \qquad b_{\text{NP}}^{\text{lin}}, \tag{S67}$$

where the splitting of $b_{\text{NP}}$ into $b_{\text{NP}}^{\text{quad}}$ and $b_{\text{NP}}^{\text{lin}}$ is motivated in the next part, Sec. "Components of the recurrence relation". $a$ is independent of the perturbation size and the same for WP and NP.

#### Optimal learning rate

The optimal learning rate and convergence factor are obtained by minimizing the quadratic function $a$ as a function of $\eta$ with respect to $\eta$,

$$\eta^* = \operatorname*{argmin}_{\eta} a(\eta) = \frac{1}{(MN_{\text{eff}}+2)\alpha^2}, \tag{S68}$$

$$a^* = \min_{\eta} a(\eta) = 1 - \frac{1}{MN_{\text{eff}}+2} = a(\eta^*). \tag{S69}$$

Learning diverges for $\eta \to 2\eta^*$ because then $a \to 1$,

$$a(2\eta^*) = 1 - \frac{4}{MN_{\text{eff}}+2} + 4\frac{MN_{\text{eff}}+2}{(MN_{\text{eff}}+2)^2} = 1. \tag{S70}$$

## SM 3

### Analysis of error dynamics.

In order to interpret and explain the main results of main text Sec. "Error dynamics", it is helpful to make four distinctions: 1) between task relevant and irrelevant weights, 2) between realizable and unrealizable outputs, 3) between informative linear, uninformative linear and uninformative quadratic contributions to the error signal, and 4) between update fluctuations due to a credit assignment problem and those due to reward noise. In the following sections we will work these distinctions out, considering first the error signal and then the thereby informed updates. Finally we use the concepts to explain the components and behavior of the recurrence relation that describes the resulting error evolution. As before, we consider training linear readouts to reduce a quadratic error. Inputs and targets exactly repeat each trial. We focus on networks where the first $N_{\text{eff}}$ eigenvalues of the input correlation matrix $S$ are equal to $\alpha^2$ and all others are zero (SM2, Sec. "Error curves for equally strong input components", main text Sec. "Theoretical analysis").

##### Task relevant and irrelevant weights

For low-dimensional input, weight changes along many directions in synaptic weight space influence neither output nor error. Here we define *relevant* and *irrelevant weight space directions* and, after an input rotation that we show leaves WP and NP learning invariant, *relevant* and *irrelevant weights*. Distinguishing task-relevant and -irrelevant weights will allow us to explain why irrelevant weights diffuse for WP but not NP, and why WP nevertheless attains the same convergence speed.

Because any $S = \frac{1}{T} r r^T$ is symmetric, it can be diagonalized by a $N \times N$ rotation matrix $O$, $D = O^T S O$. We can therefore define a set of rotated inputs $\tilde{r} = O^T r$ ($\tilde{r}_{\mu t} = \sum_{l=1}^{N} (O^T)_{\mu l} r_{lt}$) that are mutually uncorrelated as their correlation matrix is diagonal: $\frac{1}{T} \tilde{r} \tilde{r}^T = \frac{1}{T} O^T r r^T O = O^T S O = D$. The rotation does not affect the output of our networks if the reverse rotation is applied to the readout weights $\tilde{w} = wO$,

$$z_{it} = \sum_{j=1}^{N} w_{ij} r_{jt} = \sum_{j\mu l=1}^{N} w_{ij} \underbrace{O_{j\mu} \cdot (O^T)_{\mu l}}_{\Rightarrow \delta_{jl}} r_{lt} = \sum_{\mu=1}^{N} \tilde{w}_{i\mu} \tilde{r}_{\mu t} = \tilde{z}_{it}. \tag{S71}$$

The rotational invariance holds as long as inputs sum linearly, unaffected by the (possibly nonlinear) activation functions $g$. WP and NP work the same before and after the rotation of inputs because the noise is Gaussian iid and thus in particular isotropic. This is also reflected by the fact that all results only depend on the traces of powers of $S$, see Eqs. (S40, S57), which are invariant to rotations. Up to fluctuations due to different noise realizations, a WP or NP learning network thus behaves the same as its counterpart with rotated inputs, if the initial weights of the latter are transformed with the inverse rotation.

In networks where the first $N_{\text{eff}}$ eigenvalues of the input correlation matrix $S$ are equal to $\alpha^2$ and all others are zero, due to the equivalence of the learning dynamics for rotated and unrotated inputs, we can assume without loss of generality that the first $N_{\text{eff}}$ inputs are orthogonal and have strength $\alpha^2$, while the last $N - N_{\text{eff}}$ inputs are zero. Main text, Fig. 2a illustrates inputs with the assumed correlation matrix before and after the rotation. Because the last $N - N_{\text{eff}}$ inputs are always zero, all the $M(N - N_{\text{eff}})$ weights that connect them to the outputs are irrelevant for the task. Conversely, the $MN_{\text{eff}}$ weights that originate from the first $N_{\text{eff}}$ inputs are task relevant; changes of these weights affect performance. More generally, irrelevant weights occur in a network with rotated inputs whenever the obtained diagonal input correlation matrix $D$ contains zero diagonal entries. These weights correspond to irrelevant weight directions in the network with unrotated inputs.

Considering relevant and irrelevant weights instead of weight space directions, which are linear combinations of individual weights, will simplify our discussion. In all of the following we will therefore without loss of generality assume rotated inputs.

##### Realizable and unrealizable outputs

General targets and the output perturbations of NP can contain components that are not present in the inputs and thus not realizable. Here we define these components and describe their different effects in WP and NP learning. This will be crucial to understand one part of the difference between the final errors that WP and NP can obtain.

If there is a single linear readout, the inputs (i.e. the temporal input vectors) span the space of outputs (of temporal output vectors) that can be realized by adjusting the weights. If the inputs are effectively $N_{\text{eff}}$-dimensional as above, that space is also $N_{\text{eff}}$-dimensional. For $M$ outputs it is $MN_{\text{eff}}$-dimensional. The full output space can thus be split into an $MN_{\text{eff}}$-dimensional subspace of *realizable outputs*, and an orthogonal $M(T - N_{\text{eff}})$-dimensional subspace of *unrealizable outputs*.

Because WP perturbs the weights, the resulting perturbations of the outputs are always reproducible through a weight update. NP, on the other hand, perturbs all $MT$ dimensions of the full output space equally, such that only a fraction of $N_{\mathrm{eff}}/T$ of its perturbation's variance falls into the realizable subspace. However, only the realizable components of an output perturbation can be used to inform weight updates. If, for example, an unrealizable component improves performance, there is no way to capitalize on this by reproducing it through a weight update. As a consequence, the unrealizable output perturbation components of NP are a source of *reward noise*, which partially obscures the informative error changes due to better or worse generation of the realizable components. In particular, this lowers NP's performance if the target has an unrealizable component.

The worse performance of NP can be understood in more detail as follows: In presence of unrealizable target components, also the output error gradient will have a component in the unrealizable subspace. Unrealizable node perturbation components can "couple" to this component, i.e. they can have a nonzero projection onto it, contributing to the error $E^{\mathrm{pert}}$ already in linear order. To translate output perturbations into appropriate weight updates, the eligibility trace Eq. (6) projects them onto the inputs, which deletes all unrealizable perturbation components. Their contribution to $E^{\mathrm{pert}} - E$, however, still enters the update. From the perspective of the weight updates the unrealizable output perturbation components therefore just contribute noise of unknown origin to the error change $E^{\mathrm{pert}} - E$. This noise adds to the informative error change that results from realizable perturbations, which are reflected in the eligibility trace and translated into weight changes. These different contributions of the perturbations to the error signal and their effects on updates and error evolution will be quantitatively analyzed in the following sections.

#### Linear informative, linear uninformative and quadratic uninformative contributions to the error signal

In this section we distinguish the different contributions to the error signal and discuss their magnitude and scaling. This will allow us to understand the origin of the different contributions to the updates as well as their effects on the error evolution and to compare their impact for WP and NP.

The error change $\Delta E_{\mathrm{pert}} = E^{\mathrm{pert}} - E$ due to a perturbation can be split into a linear and a quadratic part,

$$\Delta E_{\mathrm{pert}} = \Delta E^{\mathrm{lin}}_{\mathrm{pert}} + \Delta E^{\mathrm{quad}}_{\mathrm{pert}}. \tag{S72}$$

Higher orders do not occur due to the use of a quadratic error function (Eq. (8)).

We first consider WP, where

$$\Delta E^{\mathrm{WP}}_{\mathrm{pert}} \;=\; \overbrace{\mathrm{tr}[W S(\xi^{\mathrm{WP}})^T]}^{\equiv \Delta E^{\mathrm{lin,WP}}_{\mathrm{pert}}} + \overbrace{\tfrac{1}{2}\,\mathrm{tr}[\xi^{\mathrm{WP}} S (\xi^{\mathrm{WP}})^T]}^{\equiv \Delta E^{\mathrm{quad,WP}}_{\mathrm{pert}}} \tag{S73}$$

(Eq. (S29)). The linear part of an error change due to weight perturbations can generally also be written as

$$\Delta E^{\mathrm{lin,WP}}_{\mathrm{pert}} = \sum_{mk} \frac{\partial E}{\partial w_{mk}} \cdot \xi^{\mathrm{WP}}_{mk}, \tag{S74}$$

cf. Eq. (S1). This shows that $\Delta E^{\mathrm{lin,WP}}_{\mathrm{pert}}$ contains all the information about the error gradient employed in weight perturbation learning, namely the size of the tried perturbation's projection onto it: The update equations Eqs. (4,S2,S16) use that if we average all normalized perturbation vectors weighted by their projections onto the gradient, the result is the gradient. $\Delta E^{\mathrm{lin}}_{\mathrm{pert}}$ provides the employed projections. In contrast, the quadratic part $\Delta E^{\mathrm{quad}}_{\mathrm{pert}}$ of $\Delta E_{\mathrm{pert}}$ does not contain such information: $\Delta E^{\mathrm{quad}}_{\mathrm{pert}}$ is uncorrelated with the size of the projection of $\xi^{\mathrm{WP}}$ onto the gradient, $\langle \Delta E^{\mathrm{lin}}_{\mathrm{pert}} \Delta E^{\mathrm{quad}}_{\mathrm{pert}} \rangle = 0$. $\Delta E^{\mathrm{quad}}_{\mathrm{pert}}$ therefore only adds reward noise to the learning process. When averaging over perturbations, this does not bias the resulting update, because $\Delta E^{\mathrm{quad}}_{\mathrm{pert}}$ is also uncorrelated with $\xi^{\mathrm{WP}}$, $\langle \Delta E^{\mathrm{quad}}_{\mathrm{pert}} \xi^{\mathrm{WP}} \rangle = 0$ (Eq. (S16)). The noise, however, entails that a larger number of perturbations needs to be tried to obtain a faithful gradient estimate and thus a good weight update.

In NP, the error change is given by

$$\Delta E^{\mathrm{NP}}_{\mathrm{pert}} \;=\; \overbrace{\tfrac{1}{T}\,\mathrm{tr}[(Wr - d)(\xi^{\mathrm{NP}})^T]}^{\Delta E^{\mathrm{lin,NP}}_{\mathrm{pert}}} + \overbrace{\tfrac{1}{2T}\,\mathrm{tr}[\xi^{\mathrm{NP}} (\xi^{\mathrm{NP}})^T]}^{\equiv \Delta E^{\mathrm{quad,NP}}_{\mathrm{pert}}} \tag{S75}$$

(Eq. (S41)). The linear term is the projection of the node (i.e. output) perturbation onto the output gradient. As discussed in Sec. "Realizable and unrealizable outputs", this gradient is composed of two orthogonal parts, laying in the realizable and in the unrealizable subspace,

$$\frac{\partial E}{\partial z} = \underbrace{\frac{1}{T}Wr}\quad \underbrace{-\frac{d}{T}} \tag{S76}$$

$$= \left[\frac{\partial E}{\partial z}\right]^{\text{real}} + \left[\frac{\partial E}{\partial z}\right]^{\text{unr}}. \tag{S77}$$

$\Delta E_{\text{pert}}^{\text{lin,NP}}$ thus consists of two components, resulting from the projection of $\xi^{\text{NP}}$ onto the gradient parts in the realizable and in the unrealizable output subspace,

$$\Delta E_{\text{pert}}^{\text{lin}} = \Delta E_{\text{pert}}^{\text{lin,real}} + \Delta E_{\text{pert}}^{\text{lin,unr}}, \tag{S78}$$

where

$$\Delta E_{\text{pert}}^{\text{lin,real}} \overset{\text{NP}}{=} \frac{1}{T}\sum_{m=1}^{M}\sum_{t=1}^{T}\sum_{j=1}^{N} W_{mj} r_{jt} \xi_{mt}^{\text{NP}} = \sum_{m=1}^{M}\sum_{t=1}^{T}\left[\frac{\partial E}{\partial z}\right]_{mt}^{\text{real}} \xi_{mt}^{\text{NP}}, \tag{S79}$$

$$\Delta E_{\text{pert}}^{\text{lin,unr}} \overset{\text{NP}}{=} -\frac{1}{T}\sum_{m=1}^{M}\sum_{t=1}^{T} d_{mt}\xi_{mt}^{\text{NP}} = \sum_{m=1}^{M}\sum_{t=1}^{T}\left[\frac{\partial E}{\partial z}\right]_{mt}^{\text{unr}} \xi_{mt}^{\text{NP}}. \tag{S80}$$

$\Delta E_{\text{pert}}^{\text{lin,unr}}$ only generates reward noise, since it is uncorrelated with the projection of the noise onto the realizable part of the gradient (which is used for learning), $\langle \Delta E_{\text{pert}}^{\text{lin,real}} \Delta E_{\text{pert}}^{\text{lin,unr}} \rangle = 0$. $\Delta E_{\text{pert}}^{\text{lin,unr}}$ increases the number of perturbations that are necessary to obtain a good weight update from averaging over them, but does not bias the resulting update (Eq. (S17)). Since $\Delta E_{\text{pert}}^{\text{lin,unr}}$ is linear in the perturbations, its effect is independent of the perturbation size (due to our division by $\sigma_{\text{NP}}$ in the NP update equation Eq. (6)). For WP $\Delta E_{\text{pert}}^{\text{lin,unr}}$ is zero. The quadratic part of $\Delta E_{\text{pert}}^{\text{NP}}$ does not contain information on the gradient either and is therefore only a source of reward noise, like the quadratic part of the error change in WP.

Only a fraction of $N_{\text{eff}}/T$ of NP's node perturbation strength (as measured by the perturbation variance), lies in the subspace of realizable outputs and can couple to the realizable part of the gradient. Thus for NP the variance of $\Delta E_{\text{pert}}^{\text{lin,real}}$ is smaller than for WP by a factor of $N_{\text{eff}}/T$,

$$\begin{aligned}
\langle\langle \Delta E_{\text{pert}}^{\text{lin,WP}} \rangle\rangle &= \left\langle \left( \sum_{i=1}^{M}\sum_{j=1}^{N} \frac{\partial E}{\partial w_{ij}} \xi_{ij}^{\text{WP}} \right)^2 \right\rangle \\
&= \sum_{im=1}^{M}\sum_{jklp=1}^{N} W_{il} S_{lj} W_{mp} S_{pk} \langle \xi_{ij}^{\text{WP}} \xi_{mk}^{\text{WP}} \rangle \\
&= \sigma_{\text{WP}}^2 \sum_{i=1}^{M}\sum_{jlp=1}^{N} W_{il} S_{lj} S_{pj} W_{ip} \\
&= \sigma_{\text{eff}}^2 (\alpha^2 N_{\text{eff}})^{-1} \operatorname{tr}[W S^2 W^T] \\
&= 2\frac{\sigma_{\text{eff}}^2}{N_{\text{eff}}}(E - E_{\text{opt}}),
\end{aligned}
\qquad
\begin{aligned}
\langle\langle \Delta E_{\text{pert}}^{\text{lin,real,NP}} \rangle\rangle &= \left\langle \left( \sum_{i=1}^{M}\sum_{t=1}^{T} \left[\frac{\partial E}{\partial z}\right]_{it}^{\text{real}} \xi_{it}^{\text{NP}} \right)^2 \right\rangle \\
&= \sum_{im=1}^{M}\sum_{jk=1}^{N}\sum_{ts}^{T} \frac{1}{T^2} W_{ij} r_{jt} W_{mk} r_{ks} \langle \xi_{it}^{\text{NP}} \xi_{ms}^{\text{NP}} \rangle \\
&= \sigma_{\text{eff}}^2 \sum_{i=1}^{M}\sum_{jk=1}^{N} \frac{1}{T} W_{ij} \sum_{t}^{T} \frac{1}{T} r_{jt} r_{kt} W_{ik} \\
&= \sigma_{\text{eff}}^2 T^{-1} \operatorname{tr}[W S W^T] \\
&= 2\frac{\sigma_{\text{eff}}^2}{T}(E - E_{\text{opt}}) = \frac{N_{\text{eff}}}{T} \cdot \langle\langle \Delta E_{\text{pert}}^{\text{lin,WP}} \rangle\rangle.
\end{aligned} \tag{S81}$$

This leads to a smaller "signal-to-noise ratio" in NP, when measuring the projection of the perturbations onto the gradient using $\Delta E_{\text{pert}}^{\text{lin,real}}$. NP has to compensate this by amplifying the learning signal in $\Delta E_{\text{pert}}^{\text{lin,real}}$ more strongly by a factor of $\sqrt{T/N_{\text{eff}}}$ to achieve the same mean update $\langle w \rangle$ as WP (Eqs. (S2,S4)). Since the learning signal and the related weight update cannot be selectively increased, the entire weight update is larger. This amplification becomes apparent when comparing the size (as measured by the standard deviation, since the mean is zero) of the different factors that $\Delta E_{\text{pert}}$ is multiplied with in the update rules Eqs. (3,6),

$$\xi_{ij}^{\text{WP}}/\sigma_{\text{WP}}^2 \sim \sigma_{\text{eff}}^{-1}\sqrt{\alpha^2 N_{\text{eff}}}, \qquad \sum_{t=1}^{T} \xi_{it}^{\text{NP}} r_{jt}/\sigma_{\text{eff}}^2 \sim \sigma_{\text{eff}}^{-1}\sqrt{\alpha^2 T} \tag{S82}$$

Here we used Eq. (S22) and assumed for NP that input $j$ is a relevant input, which has strength $\alpha^2$ (SM1, Sec. "Task setting"). For an irrelevant input, $r_j t = 0$ and $\Delta E_{\text{pert}}$ is multiplied with zero. Sec. "Strength of weight update fluctuations due to reward noise" explains how the larger weight update leads for NP to larger update fluctuations due to reward noise.

For both WP and NP, the quadratic contribution to the error change equals a normalized sum over the squared output perturbations $\delta z_{it}$,

$$\Delta E_{\text{pert}}^{\text{quad}} = \frac{1}{2T}\sum_{i=1}^{M}\sum_{t=1}^{T} \delta z_{it}^2, \tag{S83}$$

due to Eqs. (S73,S7,S18) for WP and Eqs. (S75,S19) for NP. Since the output perturbations produced by WP and NP have the same summed variance (Eqs. (S20-S22)), also the quadratic contributions to the error have to leading order the same size, which is given by the non-vanishing mean $\langle \Delta E^{\text{quad}}_{\text{pert}} \rangle$,

$$\langle \Delta E^{\text{quad}}_{\text{pert}} \rangle = \frac{1}{2T} \sum_{i=1}^{M} \sum_{t=1}^{T} \langle \delta z_{it}^2 \rangle = \tfrac{1}{2} M \sigma_{\text{eff}}^2, \tag{S84}$$

$$\begin{aligned}
\langle\langle \Delta E^{\text{quad}}_{\text{pert}} \rangle\rangle &\overset{\text{WP}}{=} \frac{1}{4T^2} \sum_{im=1}^{M} \sum_{ts=1}^{T} \sum_{jkpq=1}^{N} T^2 S_{jk} S_{pq} \langle \xi_{ij}^{\text{WP}} \xi_{ik}^{\text{WP}} \xi_{mp}^{\text{WP}} \xi_{mq}^{\text{WP}} \rangle - \langle \Delta E^{\text{quad}}_{\text{pert}} \rangle^2 \\
&= \frac{1}{4} \sigma_{\text{WP}}^4 \sum_{im=1}^{M} \sum_{ts=1}^{T} \sum_{jkpq=1}^{N} S_{jk} S_{pq} \big( \delta_{jk} \delta_{pq} + \delta_{im} (\delta_{jp} \delta_{kq} + \delta_{jq} \delta_{kp}) \big) - \frac{1}{4} M^2 \sigma_{\text{eff}}^4 \\
&= \frac{1}{4} \sigma_{\text{WP}}^4 \big( M^2 \operatorname{tr}[S]^2 + 2M \operatorname{tr}[S^2] \big) - \frac{1}{4} M^2 \sigma_{\text{eff}}^4 = \frac{1}{2} \sigma_{\text{eff}}^4 \frac{M}{N_{\text{eff}}},
\end{aligned} \tag{S85}$$

$$\begin{aligned}
\langle\langle \Delta E^{\text{quad}}_{\text{pert}} \rangle\rangle &\overset{\text{NP}}{=} \frac{1}{4T^2} \sum_{im=1}^{M} \sum_{ts=1}^{T} \langle \xi_{it}^{\text{NP}} \xi_{it}^{\text{NP}} \xi_{ms}^{\text{NP}} \xi_{ms}^{\text{NP}} \rangle - \langle \Delta E^{\text{quad}}_{\text{pert}} \rangle^2 \\
&= \frac{1}{4T^2} \sigma_{\text{eff}}^4 \sum_{im=1}^{M} \sum_{ts=1}^{T} \big( 1 + 2 \delta_{im} \delta_{ts} \big) - \frac{1}{4} M^2 \sigma_{\text{eff}}^4 = \frac{1}{2} \sigma_{\text{eff}}^4 \frac{M}{T}.
\end{aligned} \tag{S86}$$

To leading order, the size of $\Delta E^{\text{quad}}_{\text{pert}}$ scales with the perturbation strength $\sigma_{\text{eff}}^2$ and with $M$ because the $MT$-dimensional output is perturbed with a per-dimension variance of $\sigma_{\text{eff}}^2$ and the definition Eq. (8) of the error contains a factor of $1/T$ that cancels the $T$-dependence.

#### Contributions to the weight update: gradient following, credit assignment-related noise and reward noise

The mean updates of WP and NP align with the gradient and are equal to those of GD (Eqs. (S16,S17, S24)). The updates of WP and NP, however, fluctuate. This slows learning down and, if the perturbations $\xi$ are finite, also limits the final performance. In this section we show that there are two sources of update fluctuations: a credit assignment problem and reward noise. Their impact will be quantified in the subsequent two sections. Thereafter we describe how they influence the different aspects of the error evolution.

Inserting the different contributions to the error change, $\Delta E^{\text{lin,real}}_{\text{pert}}$, $\Delta E^{\text{lin,unr}}_{\text{pert}}$ and $\Delta E^{\text{quad}}_{\text{pert}}$, into the update equations Eqs. (3) and (6), allows us to split the updates into different components,

$$\Delta w_{ij}^{\text{WP}} = -\frac{\eta}{\sigma_{\text{WP}}^2} \big( \overbrace{\Delta E^{\text{lin,real}}_{\text{pert}} \xi_{ij}^{\text{WP}}}^{\text{I+II}} + \overbrace{\Delta E^{\text{quad}}_{\text{pert}} \xi_{ij}^{\text{WP}}}^{\text{IV}} \big), \tag{S87}$$

$$\Delta w_{ij}^{\text{NP}} = -\frac{\eta}{\sigma_{\text{NP}}^2} \Big( \underbrace{\Delta E^{\text{lin,real}}_{\text{pert}} \sum_{t=1}^{T} \xi_{it}^{\text{NP}} r_{jt}}_{\text{I+II}} + \underbrace{\Delta E^{\text{lin,unr}}_{\text{pert}} \sum_{t=1}^{T} \xi_{it}^{\text{NP}} r_{jt}}_{\text{III}} + \underbrace{\Delta E^{\text{quad}}_{\text{pert}} \sum_{t=1}^{T} \xi_{it}^{\text{NP}} r_{jt}}_{\text{IV}} \Big), \tag{S88}$$

which may be written as

| | | |
|---|---|---|
| $\Delta w_{ij} = \langle \Delta w_{ij} \rangle$ | (I, from $\Delta E^{\text{lin,real}}_{\text{pert}}$ - mean update) | (S89) |
| $+\, \delta w_{ij}^{\text{cr.as}}$ | (II, from $\Delta E^{\text{lin,real}}_{\text{pert}}$ - fluctuations due to credit assignment problem) | (S90) |
| $+\, \delta w_{ij}^{\text{rew.noise,lin}}$ | (III, from $\Delta E^{\text{lin,unr}}_{\text{pert}}$ - fluctuations due to linear reward noise) | (S91) |
| $+\, \delta w_{ij}^{\text{rew.noise,quad}}$ | (IV, from $\Delta E^{\text{quad}}_{\text{pert}}$ - fluctuations due to quadratic reward noise). | (S92) |

The mean update is proportional to the gradient (Eqs. (S16,S17)),

$$\langle \Delta w_{ij} \rangle = -\eta \frac{\partial E}{\partial w_{ij}}. \tag{S93}$$

The first part of the fluctuations explicitly reads (using Eqs. (S87,S87,S74,S79,S93))

$$\delta w_{ij}^{\mathrm{cr.as}} \stackrel{\mathrm{WP}}{=} -\frac{\eta}{\sigma_{\mathrm{WP}}^2}\sum_{m=1}^{M}\sum_{k=1}^{N}\frac{\partial E}{\partial w_{mk}}\xi_{mk}^{\mathrm{WP}}\cdot\xi_{ij}^{\mathrm{WP}}+\eta\frac{\partial E}{\partial w_{ij}}, \tag{S94}$$

$$\delta w_{ij}^{\mathrm{cr.as}} \stackrel{\mathrm{NP}}{=} -\frac{\eta}{\sigma_{\mathrm{NP}}^2}\sum_{m=1}^{M}\sum_{s=1}^{T}\left[\frac{\partial E}{\partial z}\right]_{ms}^{\mathrm{real}}\xi_{ms}^{\mathrm{NP}}\cdot\sum_{t=1}^{T}\xi_{it}^{\mathrm{NP}}r_{jt}+\eta\frac{\partial E}{\partial w_{ij}} \tag{S95}$$

It can be attributed to the *credit assignment* problem of finding, out of the $MN$ weights or $MT$ outputs, the single gradient-parallel component of the perturbation that was responsible for causing $\Delta E_{\mathrm{pert}}^{\mathrm{lin,real}}$, i.e. the linear, instructive part of the error signal. More specifically: Because from the scalar error signal alone it is impossible for WP to determine which of the $MN$ sampled directions was the one parallel to the gradient, WP has to amplify the perturbations $\xi_{ij}^{\mathrm{WP}}$ of all weights equally during the constructions of their updates (Eqs. (3,S94)). This implies that all weights fluctuate. The single backpropagation step of NP, reflected in its use of eligibility traces (Eqs. (6,S95)), allows it to solve the credit assignment problem at least partially. Therefore for NP only the $MN_{\mathrm{eff}}$ relevant weights are updated. The convergence speed is, however, only limited by the update noise of relevant weights, which is the same for NP and WP, see Eqs. (S102,S103) and compare the identical convergence factors of WP and NP, Eq. (S62).

The second kind of update fluctuations, $\delta w_{ij}^{\mathrm{rew.noise,lin}}$ and $\delta w_{ij}^{\mathrm{rew.noise,quad}}$, read (Eqs. (S91,S80))

$$\delta w_{ij}^{\mathrm{rew.noise,lin}} \stackrel{\mathrm{WP}}{=} 0, \qquad \delta w_{ij}^{\mathrm{rew.noise,lin}} \stackrel{\mathrm{NP}}{=} \frac{\eta}{\sigma_{\mathrm{NP}}^2}\sum_{m=1}^{M}\sum_{s=1}^{T}\frac{1}{T}d_{ms}\xi_{mt}^{\mathrm{NP}}\cdot\sum_{t=1}^{T}\xi_{it}^{\mathrm{NP}}r_{jt} \tag{S96}$$

and (Eqs. (S92,S88,S83))

$$\delta w_{ij}^{\mathrm{rew.noise,quad}} \stackrel{\mathrm{WP}}{=} -\frac{\eta}{\sigma_{\mathrm{WP}}^2}\frac{1}{2T}\sum_{m=1}^{M}\sum_{s=1}^{T}\left(\sum_{k=1}^{N}\xi_{mk}^{\mathrm{WP}}r_{ks}\right)^2\cdot\xi_{ij}^{\mathrm{WP}}, \tag{S97}$$

$$\delta w_{ij}^{\mathrm{rew.noise,quad}} \stackrel{\mathrm{NP}}{=} -\frac{\eta}{\sigma_{\mathrm{NP}}^2}\frac{1}{2T}\sum_{m=1}^{M}\sum_{s=1}^{T}\left(\xi_{ms}^{\mathrm{NP}}\right)^2\cdot\sum_{t=1}^{T}\xi_{it}^{\mathrm{NP}}r_{jt}. \tag{S98}$$

They are caused by reward noise ($\Delta E_{\mathrm{pert}}^{\mathrm{lin,unr}}$ and $\Delta E_{\mathrm{pert}}^{\mathrm{quad}}$, respectively). Multiplying the reward noise in $E^{\mathrm{pert}}-E$ with the applied perturbations or eligibility traces in the construction of updates, Eqs. (3,6), results in random update contributions that are unrelated to the gradient (see Eqs. (S97,S98) in contrast to Eqs. (S94,S95))). Because the reward noise is independent of the weight mismatch, the amplitudes of these fluctuations do not change over the course of training, which prevents learning with arbitrary precision. In contrast, the informative $\Delta E_{\mathrm{pert}}^{\mathrm{lin,real}}$ diminishes with training.

#### Strength of credit assignment-related weight update fluctuations and the dimensionality argument for our task

This section computes and compares the fluctuation strengths due to the credit-assignment problem in WP and NP learning. Thereafter it quantitatively states a dimensionality argument, which adapts the one that is commonly used to compare WP, NP and GD learning (main text, introduction) to our type of task.

The strength of the update noise due to the credit assignment problem, i.e. the variance of $\delta w_{ij}^{\mathrm{cr.as}}$, depends only on the contribution $\Delta E_{\mathrm{pert}}^{\mathrm{lin,real}}$ of the error change (Line (S90)) and is thus independent of $d$ and $\sigma_{\mathrm{eff}}$ (Eqs. (S87,S88,S73,S79)). For WP it is

$$\begin{aligned}\langle\langle\delta w_{ij}^{\mathrm{cr.as}}\rangle\rangle &\stackrel{\mathrm{WP}}{=} \left\langle\left(-\tfrac{\eta}{\sigma_{\mathrm{WP}}^2}\Delta E_{\mathrm{pert}}^{\mathrm{lin}}\xi_{ij}^{\mathrm{WP}}\right)^2\right\rangle-\langle\Delta w_{ij}\rangle^2\\ &= \tfrac{\eta^2}{\sigma_{\mathrm{WP}}^4}\sum_{mn=1}^{M}\sum_{kl=1}^{N}\frac{\partial E}{\partial w_{mk}}\frac{\partial E}{\partial w_{nl}}\underbrace{\left\langle\xi_{mk}^{\mathrm{WP}}\xi_{nl}^{\mathrm{WP}}\xi_{ij}^{\mathrm{WP}}\xi_{ij}^{\mathrm{WP}}\right\rangle}_{=\sigma_{\mathrm{WP}}^4(\delta_{mn}\delta_{kl}+2\delta_{mni}\delta_{jkl})}-\eta^2\left(\frac{\partial E}{\partial w_{ij}}\right)^2\\ &= \eta^2\left|\frac{\partial E}{\partial w}\right|^2+\eta^2\left(\frac{\partial E}{\partial w_{ij}}\right)^2.\end{aligned} \tag{S99}$$

These credit assignment-related random fluctuations in the update of $w_{ij}$ thus grow quadratically with the overall length of the error gradient and with the size of its component in $w_{ij}$-direction. The first dependency arises because any weight perturbation is multiplied with the global error change, which is in linear order proportional to the gradient length. The second dependency arises because weight changes parallel to the gradient fluctuate twice as strongly as in perpendicular directions due to their correlation with error changes.

For NP, the credit assignment-dependent weight update noise strength is

$$\begin{aligned}
\langle\langle \delta w_{ij}^{\mathrm{cr.as}}\rangle\rangle &\overset{\mathrm{NP}}{=} \left\langle \left( -\frac{\eta}{\sigma_{\mathrm{NP}}^2} \Delta E_{\mathrm{pert}}^{\mathrm{lin,real}} \sum_{t=1}^{T} \xi_{it}^{\mathrm{NP}} r_{jt} \right)^2 \right\rangle - |\langle \Delta w_{ij} \rangle|^2 \\
&= \frac{\eta^2}{\sigma_{\mathrm{NP}}^4} \sum_{mn=1}^{M} \sum_{stuv=1}^{T} \left[\frac{\partial E}{\partial z}\right]_{ms}^{\mathrm{real}} \left[\frac{\partial E}{\partial z}\right]_{nt}^{\mathrm{real}} \underbrace{\left\langle \xi_{ms}^{\mathrm{NP}} \xi_{nt}^{\mathrm{NP}} \xi_{iu}^{\mathrm{NP}} \xi_{iv}^{\mathrm{NP}} \right\rangle}_{=\sigma_{\mathrm{NP}}^4 \left(\delta_{mn}\delta_{st}\delta_{uv} + \delta_{imn}(\delta_{su}\delta_{tv} + \delta_{sv}\delta_{tu})\right)} r_{ju} r_{jv} - \eta^2 \left(\frac{\partial E}{\partial w_{ij}}\right)^2 \\
&= \eta^2 \left| \left[\frac{\partial E}{\partial z}\right]^{\mathrm{real}} \right|^2 T S_{jj} + \eta^2 \left(\frac{\partial E}{\partial w_{ij}}\right)^2 .
\end{aligned} \tag{S100}$$

The first term is proportional to the squared length of the realizable part of the output gradient and to the strength of the $j$th input. This is because in the weight update rule node perturbations are multiplied with the global error change (which is in turn proportional to the output gradient length) and with the $j$th input. The second term arises again because weight changes in the direction of the gradient fluctuate twice as strongly due to their correlation with the error change. The first term is generally different from that of WP, which can cause differences in convergence speeds - compare SM6 and main text, Sec. “Reservoir computing-based drawing task”. For the case of equally strong latent inputs, which we focus on in our analytical computations, however, noise variances are equal for WP and NP: in the rotated space of input components $r_{\mu t}$ introduced in Sec. “Task relevant and irrelevant weights”, the squared norm of the weight error gradient is

$$\begin{aligned}
\left|\frac{\partial E}{\partial w}\right|^2 &= \sum_{m=1}^{M} \sum_{\mu=1}^{N} \left(\frac{\partial E}{\partial w_{m\mu}}\right)^2 = \sum_{m=1}^{M} \sum_{\mu=1}^{N} \left( \sum_{t=1}^{T} \frac{\partial E}{\partial z_{mt}} r_{\mu t} \right)^2 \\
&= \sum_{m=1}^{M} \sum_{\mu=1}^{N_{\mathrm{eff}}} \left( \frac{\partial E}{\partial z_m}, \hat{r}_\mu \right)^2 T\alpha^2 \overset{(*)}{=} \left| \left[\frac{\partial E}{\partial z}\right]^{\mathrm{real}} \right|^2 T\alpha^2 .
\end{aligned} \tag{S101}$$

Here we introduced the temporal unit vectors $\hat{r}_\mu = \frac{1}{\sqrt{T\alpha}} r_\mu$ and used in $(*)$ that the $\hat{r}_\mu$ with $\mu = 1, ..., N_{\mathrm{eff}}$ form a basis for the subspace in which the realizable part of the node gradient lies. In the space of rotated inputs we thus have

$$\langle\langle \delta w_{i\mu}^{\mathrm{cr.as}} \rangle\rangle \overset{\mathrm{WP}}{=} \eta^2 \left|\frac{\partial E}{\partial w}\right|^2 + \eta^2 \left(\frac{\partial E}{\partial w_{i\mu}}\right)^2 \qquad \text{for every input } \mu, \tag{S102}$$

$$\langle\langle \delta w_{i\mu}^{\mathrm{cr.as}} \rangle\rangle \overset{\mathrm{NP}}{=} \begin{cases} \eta^2 \left|\frac{\partial E}{\partial w}\right|^2 + \eta^2 \left(\frac{\partial E}{\partial w_{i\mu}}\right)^2 & \text{for relevant inputs } \mu = 1, \cdots, N_{\mathrm{eff}}, \\ 0 & \text{for irrelevant inputs } \mu = N_{\mathrm{eff}} + 1, \cdots, N. \end{cases} \tag{S103}$$

Eq. (S102) reduces for an irrelevant input $\mu$ to $\langle\langle \delta w_{i\mu}^{\mathrm{cr.as}} \rangle\rangle \overset{\mathrm{WP}}{=} \eta^2 \left|\frac{\partial E}{\partial w}\right|^2$, i.e. for WP there are credit assignment-related update fluctuations also in irrelevant weight space directions, in contrast to NP. Eqs. (S102,S103) imply that the average change of a single relevant weight due to credit assignment noise, $\sqrt{\langle\langle \delta w_{i\mu}^{\mathrm{cr.as}} \rangle\rangle}$, is at least as large as the size of the deterministic improvement of the entire weight vector along the gradient, $\eta \left|\frac{\partial E}{\partial w}\right| = |\langle \Delta w \rangle|$. The average size of the entire weight vector change due to credit assignment noise can be computed by summing over Eqs. (S102,S103): Using $\sum_{i,\mu} \eta^2 \left(\frac{\partial E}{\partial w_{i\mu}}\right)^2 = \eta^2 \left|\frac{\partial E}{\partial w}\right|^2 = |\langle \Delta w \rangle|^2$ yields

$$\langle |\delta w^{\mathrm{cr.as}}|^2 \rangle = \begin{cases} (MN + 1) \cdot |\langle \Delta w \rangle|^2 & \text{for WP,} \\ (MN_{\mathrm{eff}} + 1) \cdot |\langle \Delta w \rangle|^2 & \text{for NP.} \end{cases} \tag{S104}$$

This means that the weight change due to credit assignment noise is much larger than that due to the deterministic update $\langle \Delta w \rangle$. The contributions of noise and deterministic gradient following simply add up to the total average square weight change,

$$\langle |\Delta w|^2 \rangle = |\langle \Delta w \rangle|^2 + \langle |\delta w^{\mathrm{cr.as}}|^2 \rangle, \tag{S105}$$

in absence of reward noise. Since update noise will influence and often increase the error in a nonlinear way (Eq. (8)), it might seem as if learning is impossible. However, for sufficiently small updates also with a nonlinear error function the deterministic update parts add up, as they always point into approximately the same direction, while the fluctuations partly cancel each other. For infinitesimally small update size, after $n$ updates, the mean weight change is $n \cdot \langle \Delta w \rangle$, while the standard deviation of the summed fluctuations of a single weight scales only as $\sqrt{n} \cdot \sqrt{\langle |\delta w^{\mathrm{cr.as}}|^2 \rangle}$. This way WP and NP can still learn by adopting a smaller learning rate and averaging out fluctuations over more updates.

Eq. (S104) seems to furthermore imply that NP is much more efficient in learning than WP, as its noise is much smaller. This is the classical dimensionality argument cited in the Introduction section of the main text. However, for a single input-output learning task noise-related changes of only the $MN_{\mathrm{eff}}$ task relevant weights influence the error. Thus, the relevant amount of credit assignment noise is actually the same for WP and NP, leading to the same maximal speed of convergence (Fig. 1).

#### Strength of weight update fluctuations due to reward noise

Here we give the scaling of reward noise-related update fluctuations and explain why these are larger for NP than for WP, which will explain the larger final error of NP.

The scaling of $\delta w_{ij}^{\mathrm{rew.noise,lin}}$ for NP follows from Eqs. (S91,S88,S80). Eq. (S80) yields $\Delta E_{\mathrm{pert}}^{\mathrm{lin,unr}} \overset{\mathrm{NP}}{\sim} \sigma_{\mathrm{eff}} \alpha_d \sqrt{\frac{M}{T}}$, since the "size" of the sum of centered noise terms scales with the square root of the number of summands. We here consider as the size of $\delta w_{ij}^{\mathrm{rew.noise,lin}}$ the standard deviation of the sum (Eq. (S88), term III), as its average vanishes. The formal reason that this common approach works is that we compute the strengths (variances) of noise terms $\delta w_{ij}^{\mathrm{rew.noise,lin}}$ from a product of two independent random variables $X = \Delta E_{\mathrm{pert}}^{\mathrm{lin,unr}}$ and $Y = \sum_{t=1}^{T} \xi_{it}^{\mathrm{NP}} r_{jt}$, which both have zero mean. The well known general formula $\mathrm{Var}(XY) = \mathrm{E}(X)^2 \mathrm{Var}(Y) + \mathrm{E}(Y)^2 \mathrm{Var}(X) + \mathrm{Var}(X)\mathrm{Var}(Y)$ for independent random variables $X, Y$, thus tells us that the leading order scaling in each random variable arises from its squared average or from its variance, i.e. its squared standard deviation. For our computations we therefore consider as characteristic size of a term its average or its standard deviation, depending on which has the leading order scaling, and use its square to compute the final noise variance scaling. Thus the variance of $\delta w^{\mathrm{rew.noise,lin}}$ can be obtained by simply multiplying the variances of $\Delta E_{\mathrm{pert}}^{\mathrm{lin,unr}}$ and $\sum_{t=1}^{T} \xi_{it}^{\mathrm{NP}} r_{jt}$,

$$\langle\langle \delta w_{ij}^{\mathrm{rew.noise,lin}} \rangle\rangle \overset{\mathrm{WP}}{=} 0, \tag{S106}$$

$$\begin{aligned} \langle\langle \delta w_{ij}^{\mathrm{rew.noise,lin}} \rangle\rangle &\overset{\mathrm{NP}}{=} \frac{\eta^2}{\sigma_{\mathrm{eff}}^4} \cdot \langle\langle \Delta E_{\mathrm{pert}}^{\mathrm{lin,unr}} \rangle\rangle \cdot \Big\langle\Big\langle \sum_{t=1}^{T} \xi_{it}^{\mathrm{NP}} r_{jt} \Big\rangle\Big\rangle \\ &= \eta^2 \alpha_d^2 \alpha^2 M = 2\eta^2 \alpha^2 E_{\mathrm{opt}} \qquad \text{for relevant weights only.} \end{aligned} \tag{S107}$$

(We have thus obtained the size of the product from the product of sizes, as for deterministic quantities.) We conclude that the size of $\delta w^{\mathrm{rew.noise,lin}}$ is $0$ for WP and scales for NP as

$$\delta w_{ij}^{\mathrm{rew.noise,lin}} \overset{\mathrm{NP}}{\sim} \eta \alpha_d \sqrt{\alpha^2 M} \qquad \text{for relevant weights only.} \tag{S108}$$

The scaling of $\delta w_{ij}^{\mathrm{rew.noise,quad}}$ can be computed likewise: $\Delta E_{\mathrm{pert}}^{\mathrm{quad}}$ is a sum of independent positive random variables (Eq. (S83)) such that its size has a contribution from the summed means (Eq. (S84)), which scales with $M$, and a contribution from the summed fluctuations (Eqs. (S85,S86)), which scales with $\sqrt{M}$. As $\xi_{ij}^{\mathrm{WP}}$ and $\sum_{t=1}^{T} \xi_{it}^{\mathrm{NP}} r_{jt}$ have zero mean, we obtain the full expressions

$$\begin{aligned} \langle\langle \delta w_{ij}^{\mathrm{rew.noise,quad}} \rangle\rangle &\overset{\mathrm{WP}}{=} \frac{\eta^2}{\sigma_{\mathrm{WP}}^4} \cdot \left( \langle \Delta E_{\mathrm{pert}}^{\mathrm{quad}} \rangle^2 + \langle\langle \Delta E_{\mathrm{pert}}^{\mathrm{quad}} \rangle\rangle \right) \cdot \langle\langle \xi_{ij}^{\mathrm{WP}} \rangle\rangle \\ &= \frac{\eta^2}{\sigma_{\mathrm{WP}}^2} \cdot \left( \frac{1}{4} M^2 \sigma_{\mathrm{eff}}^4 + \frac{1}{2} \sigma_{\mathrm{eff}}^4 \frac{M}{N_{\mathrm{eff}}} \right) = \frac{1}{4} \eta^2 \sigma_{\mathrm{eff}}^2 \alpha^2 \left( M^2 N_{\mathrm{eff}} + 2M \right) \end{aligned} \tag{S109}$$

$$\begin{aligned} \langle\langle \delta w_{ij}^{\mathrm{rew.noise,quad}} \rangle\rangle &\overset{\mathrm{NP}}{=} \frac{\eta^2}{\sigma_{\mathrm{eff}}^4} \cdot \left( \langle \Delta E_{\mathrm{pert}}^{\mathrm{quad}} \rangle^2 + \langle\langle \Delta E_{\mathrm{pert}}^{\mathrm{quad}} \rangle\rangle \right) \cdot \langle\langle \sum_{t=1}^{T} \xi_{it}^{\mathrm{NP}} r_{jt} \rangle\rangle \\ &= \frac{\eta^2}{\sigma_{\mathrm{eff}}^4} \cdot \left( \frac{1}{4} M^2 \sigma_{\mathrm{eff}}^4 + \frac{1}{2} \sigma_{\mathrm{eff}}^4 \frac{M}{T} \right) \cdot \sigma_{\mathrm{eff}}^2 \sum_{t=1}^{T} r_{jt}^2 \\ &= \begin{cases} \frac{1}{4} \eta^2 \sigma_{\mathrm{eff}}^2 \alpha^2 \left( M^2 T + 2M \right) & \text{for relevant weights} \\ 0 & \text{for irrelevant weights} \end{cases} \end{aligned} \tag{S110}$$

for the noise variances. To leading order, the size of update fluctuations induced by quadratic reward noise is thus

$$\delta w_{ij}^{\text{rew.noise,quad}} \overset{\text{WP}}{\approx} \frac{1}{2}\eta\sigma_{\text{eff}}\alpha M\sqrt{N_{\text{eff}}} \qquad \text{for all weights,} \tag{S111}$$

$$\delta w_{ij}^{\text{rew.noise,quad}} \overset{\text{NP}}{\approx} \frac{1}{2}\eta\sigma_{\text{eff}}\alpha M\sqrt{T} \qquad \text{for relevant weights only.} \tag{S112}$$

The different scaling ultimately results from the fact that the output variability of NP in the realizable output subspace is by a factor of $N_{\text{eff}}/T$ smaller than for WP (Eq. (S81)). NP has to compensate this by amplifying the learning signal in $\Delta E_{\text{pert}}^{\text{lin,real}}$ more strongly by a factor of $\sqrt{T/N_{\text{eff}}}$ (Eq. (S82)) to achieve the same beneficial mean update $\langle\Delta w\rangle$. It is, however, unavoidable to apply the stronger amplification to the entire $\Delta E_{\text{pert}}$. Because its part $\Delta E_{\text{pert}}^{\text{quad}}$ is – in contrast to $\Delta E_{\text{pert}}^{\text{lin,real}}$ – to leading order the same for WP and NP (Eq. (S84)), the stronger overall amplification leads to larger fluctuations $\delta w_{\text{NP}}^{\text{rew.noise,quad}} \approx \sqrt{T/N_{\text{eff}}} \cdot \delta w_{\text{WP}}^{\text{rew.noise,quad}}$ in NP.

#### Components of the recurrence relation

Here we leverage the concepts developed in the preceding sections to explain the origin and scaling of each component of the recurrence relation. Together with the next section, in which the recurrence relation is solved and the error dynamics characterized, this provides a rigorous understanding of how different task properties affect specific aspects of the error evolution.

Distinguishing the contributions $\langle\Delta w_{ij}\rangle$, $\delta w_{ij}^{\text{cr.as}}$, $\delta w_{ij}^{\text{rew.noise,lin}}$ and $\delta w_{ij}^{\text{rew.noise,quad}}$ (Lines (S89-S92)) to the updates $\Delta w$ allows to examine their different effects on the evolution of the expected error. The fluctuations are independent of each other and have zero expectation value. The expected error after an update has the following components:

$$\langle E(n)\rangle = \tfrac{1}{2}\langle \text{tr}[(W(n-1)+\Delta w)S(W(n-1)+\Delta w)^T]\rangle + E_{\text{opt}} \tag{S113}$$

$$= \langle E(n-1)\rangle \qquad \text{(error before update)} \tag{S114}$$

$$+\,\text{tr}[WS\langle\Delta w\rangle^T] + \tfrac{1}{2}\text{tr}[\langle\Delta w\rangle S\langle\Delta w\rangle^T] \qquad \text{(mean updates follow gradient as for GD)} \tag{S115}$$

$$+\,\tfrac{1}{2}\langle\text{tr}[\delta w^{\text{cr.as}}S(\delta w^{\text{cr.as}})^T]\rangle \qquad \text{(update noise from credit assignment)} \tag{S116}$$

$$+\,\tfrac{1}{2}\langle\text{tr}[\delta w^{\text{rew.noise,lin}}S(\delta w^{\text{rew.noise,lin}})^T]\rangle \qquad \text{(update noise from linear reward noise)} \tag{S117}$$

$$+\,\tfrac{1}{2}\langle\text{tr}[\delta w^{\text{rew.noise,quad}}S(\delta w^{\text{rew.noise,quad}})^T]\rangle \qquad \text{(update noise from quadratic reward noise).} \tag{S118}$$

Because the mean update is equal to that of GD, the first two contributions to $\langle E(n)\rangle$ would lead to

$$E(n) = (1-2\eta\alpha^2+\eta^2\alpha^4)\cdot\left(E(n-1)-E_{\text{opt}}\right)+E_{\text{opt}} \qquad \text{(for (S114) and (S115)),} \tag{S119}$$

cf. Eqs. (S25) to (S28). $-2\eta\alpha^2$ results from the beneficial term linear in $\langle\Delta w\rangle$, while $\eta^2\alpha^4$ results from the quadratic effect of an update perfectly parallel to the gradient on the error.

The third contribution, (S116), occurs because WP and NP solve the credit assignment problem by random search. The resulting update fluctuations $\delta w^{\text{cr.as}}$ add a diffusive part to the evolution of relevant weights (or, for non-rotated input space: in the relevant weight subspace), which on average leads to an increase in error. Fluctuations of irrelevant weights present in WP (Eq. (S102)) do not contribute; this is reflected by the multiplication of $\delta w^{\text{cr.as}}$ with $S$. Eq. (S104) shows that (S116) yields a detrimental quadratic term that is $(MN_{\text{eff}}+1)$ larger than the detrimental quadratic term in (S115). We therefore have

$$E(n) = \underbrace{(1-2\eta\alpha^2+\eta^2\alpha^4(MN_{\text{eff}}+2))}_{=a}\cdot\left(E(n-1)-E_{\text{opt}}\right)+E_{\text{opt}} \qquad \text{(for (S114) to (S116)),} \tag{S120}$$

which implicates a $(MN_{\text{eff}}+2)$ times smaller optimal learning rate than with GD (Eqs. (S68,S28)).

The increase in error resulting from the last two contributions, (S117) and (S118) stems from the update fluctuations $\delta w^{\text{rew.noise}}$ due to reward noise. Because these fluctuations are independent of gradient and weights (Eqs. (S109,S110)), the magnitude of their additive contribution does not change over the course of training. The contribution therefore yields the per-update error increase $b_{\text{WP|NP}}$ of the recurrence relation (Eqs. (8,S60)). $b$ can be split into two parts $b = b^{\text{lin}} + b^{\text{quad}}$ (Eq. (S67)) according to the reward noise source: $b^{\text{lin}}$ is only present for NP and arises from the linear reward noise due to unrealizable parts of the target (S117). $b^{\text{quad}}$ arises from quadratic reward noise due to the quadratic nonlinearity of the error function (S118). Thus, including all five contributions results in

$$E(n) = (1-2\eta\alpha^2+\eta^2\alpha^4(MN_{\text{eff}}+2))\cdot\left(E(n-1)-E_{\text{opt}}\right)+b^{\text{lin}}+b^{\text{quad}}+E_{\text{opt}} \qquad \text{(for (S114) to (S118)).} \tag{S121}$$

$b_{\text{WP}}$, $b_{\text{NP}}^{\text{lin}}$ and $b_{\text{NP}}^{\text{quad}}$ are given in Eqs. (S65–S67).

#### Characteristics of the average error dynamics

With all components of the recurrence relation explained and quantified, here we characterize the resulting dynamics of the average error. In particular, we derive the scaling of the final error.

The recurrence relation Eqs. (S60,S121),

$$\left(\langle E(n)\rangle - E_{\text{opt}}\right) = \left(\langle E(n-1)\rangle - E_{\text{opt}}\right)\cdot a + b, \tag{S122}$$

leads, provided that $a < 1$, to an exponential decay (cf. Eq. (S61))

$$\langle E(n)\rangle = \left(E(0) - E_{f,\text{unr}}\right)\cdot a^n + E_{f,\text{unr}} \tag{S123}$$

to a final error

$$E_{f,\text{unr}} = E_f + E_{f,d} + E_{\text{opt}}, \qquad E_f = \frac{b^{\text{quad}}}{1-a}, \qquad E_{f,d} = \frac{b^{\text{lin}}}{1-a}. \tag{S124}$$

$E_{f,\text{unr}}$ is the total final error including contributions due to unrealizable target compoents. $E_f$ captures the final error for the case of realizable targets. For NP unrealizable target components $d$ increase the final error by $E_{f,d}$ (which is zero for WP). Further, unrealizable target components increase the final error by $E_{\text{opt}}$, i.e. by the error that necessarily remains even for optimal weights (Eq. (S15)), because the target is not realizable by the network. Since $E_{\text{opt}}$ is known beforehand and for any learning rule represents an inevitable shift in error, it could be absorbed into a redefined $E$. The final error is then still limited by the accumulation of error due to reward noise, which leads to update noise entering the recurrence relation through $b$ (Eq. (S121) and lines (S117, S118)).

In the beginning of learning, when the weight mismatch is large, following the large gradient leads to fast improvements. As the weights approach their target values, the gradient becomes smaller, but the error contributions that result from reward noise stay constant. Gradient-related improvements and deterioration due to reward noise on average balance if the error $E$ has the size of the final error of the average error dynamics, $E = E_{f,\text{unr}}$. Around this point learning yields hardly any or no improvement. If $E$ happens to become smaller than $E_{f,\text{unr}}$, learning even has a deteriorating effect on average, because of the reward noise.

The deteriorating effect of quadratic reward noise due to finite perturbations is present in both WP and NP and unaffected by unrealizable parts of targets. It may be best illustrated for the case where all relevant weights have already assumed their target values: Then the weight error gradient is zero. Still, a finite random perturbation of the weights or outputs leads with probability one to an increase in error due to the quadratic error function. Therefore the update rules induce a finite weight change in the direction opposite to the perturbation. This prevents the weights from staying at or reaching optimal values and leads to a final error larger than $E_{\text{opt}}$.

We note that the recurrence relation Eq. (S60) may also be understood as a leaky integration of $b$. For slow convergence, the factor $1-a$ in the denominator (which arises from the discrete updating process) can be interpreted as the leak rate $-\ln(a) \approx 1-a$ of a corresponding continuous exponential decay $\sim \exp(-\ln(a)t)$ (Fig. 1) that equals the actual decay at the points where $t \in \mathbb{N}_0$. In this picture, the contribution $b/(1-a)$ to the final error results from integrating the per-update error increase $b$ over $1/(1-a)$ updates. This determines the error once the contribution due to the initialization, $\left(E(0) - E_{f,\text{unr}}\right)\cdot a^n$, has faded away.

The leading order contributions in $M$, $N_{\text{eff}}$ and $T$ to $E_f$ as given in Eqs. (13),(14) can be computed using Eqs. (S65,S66,S62) or line (S118) and Eqs. (S111,S112,S62) (note that $E_f = \frac{b^{\text{quad}}}{1-a}$ does not incorporate $b^{\text{lin}}$ and that line (S118) equals $b^{\text{quad}}$),

$$E_f^{\text{WP}} = \frac{b_{\text{WP}}(\eta)}{1-a(\eta)} \approx \frac{\sigma_{\text{eff}}^2}{8}\cdot \eta^2\alpha^4 \frac{MN_{\text{eff}}}{1-a(\eta)}\cdot M^2 N_{\text{eff}}, \tag{S125}$$

$$E_f^{\text{NP}} = \frac{b_{\text{NP}}^{\text{quad}}(\eta)}{1-a(\eta)} \approx \frac{\sigma_{\text{eff}}^2}{8}\cdot \underbrace{\eta^2\alpha^4 \frac{MN_{\text{eff}}}{1-a(\eta)}}_{\equiv c(\eta)}\cdot M^2 T. \tag{S126}$$

For our discussion, we have split the two results into three corresponding factors: (i) The first factor, $\sigma_{\text{eff}}^2/8$, reflects that $E_f^{\text{WP|NP}}$ will be of significant size only when the noise strength $\sigma_{\text{eff}}^2$ is sufficiently large. In this case the quadratic reward noise becomes sizeable and corrupts learning. (ii) The second factor contains the dependence of the final error on the learning rate, via a factor that we abbreviate by $c(\eta)$,

$$c(\eta) \equiv \eta^2\alpha^4 \frac{MN_{\text{eff}}}{1-a(\eta)}, \quad c(\eta^*) \approx 1. \tag{S127}$$

Here it is most important that this factor is approximately $1$ at the optimal learning rate $\eta^*$ (Eq. (12)), which we usually consider throughout the paper. $c(\eta)$ goes to zero for small learning rates (Eq. (S62)) and diverges for $\eta \to 2\eta^*$, since then $a \to 1$ (Eq. (S70)). We will furthermore see below that it describes the $\eta$-dependence of the ratio between diffusion of irrelevant and improvements of relevant weights (Eq. (S134)), as well as the additional increase in final error of NP caused by unrealizable target components (Eq. (S128)). (iii) The last factor originates from the scaling of the update fluctuations $\delta w^{\text{rew.noise,quad}}$, Eqs. (S111,S112), which enter quadratically. It can be further refactored as $M$ times the effective perturbation dimension $MN_{\text{eff}}$ or $MT$. The latter reflects the fact that WP and NP generate perturbations in spaces of dimensions $MN_{\text{eff}}$ and $MT$, but only the projection onto the output gradient is useful for learning. The other factor $M$ originates from the overall scaling of the error $E$ with $M$, as the error function Eq. (8) contains a sum over $M$ outputs. In other words, if $E$ was fully normalized, by $1/(MT)$, $E_f$ would scale with the effective perturbation dimension.

The additional contribution to the final error of NP due to the presence of unrealizable target components, $E_{f,d}^{\text{NP}}$ (Eq. (S124)), can be obtained from Eqs. (S66,S62) or line (S117) and Eq. (S62),

$$E_{f,d}^{\text{WP}} = 0, \qquad E_{f,d}^{\text{NP}} = \frac{b^{\text{lin}}}{1-a} = c(\eta) \cdot E_{\text{opt}}, \tag{S128}$$

with the factor $c(\eta)$ from Eq. (S127). Since $c(\eta^*) \approx 1$, learning at $\eta^*$ means that unrealizable target components increase NP's final error by approximately $2E_{\text{opt}}$ instead of $E_{\text{opt}}$ as for WP (Fig. 3). As $E_{\text{opt}}$ is independent of $\sigma_{\text{eff}}$, $E_{f,d}^{\text{NP}}$ remains finite even in the limit of infinitesimally small perturbations where $E_f$ vanishes. For small perturbations, $E_{f,d}^{\text{NP}}$ and the unavoidable $E_{\text{opt}}$ can therefore become the dominant contributions to the final error.

**SM 4**

### Analysis of Weight Diffusion

This part provides a quantitative mathematical analysis of the weight changes in irrelevant directions of the weight space.

#### Weight diffusion due to credit assignment- and reward noise-related update fluctuations

As described in SM3, Sec. "Task relevant and irrelevant weights", we can assume without loss of generality that the first $N_{\mathrm{eff}}$ inputs are orthogonal and of strength $\alpha^2$, while the remaining $N - N_{\mathrm{eff}}$ inputs are zero. The first $N_{\mathrm{eff}}$ synaptic weights are thus relevant in the sense that their value influences the output while the remaining ones are irrelevant. How do irrelevant weights change during the modeled learning? For NP the update $\Delta w_{ij}^{\mathrm{NP}}$ is proportional to the eligibility trace $\sum_t \xi_{it}^{\mathrm{NP}} r_{jt}$ (Eq. (6)), which is zero when the $j$th input is zero, $r_{jt} = 0$, for all $t$. Therefore NP does not update irrelevant weights. WP, on the other hand, perturbs and updates also the irrelevant weights (Eq. (3)). In the following we therefore focus on WP.

The evolution of the relevant weights is independent of the evolution of the irrelevant weights. This is because only the relevant weights influence the reward and they are perturbed independently of the irrelevant ones. The independence is echoed in the fact that the learning characteristics of WP only depend on $N_{\mathrm{eff}}$ but not on $N$ (Eqs. (S62,S65)). The perturbations of the irrelevant weights are also independent of each other, such that it suffices to consider the evolution of a single one. We call it $w_{\mathrm{irrel}}$ and its perturbation $\xi_{\mathrm{irrel}}^{\mathrm{WP}}$. The expectation value of its update $\Delta w_{\mathrm{irrel}}^{\mathrm{WP}}$ is zero as $\xi_{\mathrm{irrel}}^{\mathrm{WP}}$ does not influence the reward,

$$\langle \Delta w_{\mathrm{irrel}}^{\mathrm{WP}} \rangle = -\frac{\eta}{\sigma_{\mathrm{WP}}^2} \langle \underbrace{\left(E^{\mathrm{pert}} - E\right)}_{\text{indep. of } \xi_{\mathrm{irrel}}^{\mathrm{WP}}} \xi_{\mathrm{irrel}}^{\mathrm{WP}} \rangle = -\frac{\eta}{\sigma_{\mathrm{WP}}^2} \langle E^{\mathrm{pert}} - E \rangle \langle \xi_{\mathrm{irrel}}^{\mathrm{WP}} \rangle = 0. \tag{S129}$$

This implies that the expectation value of the irrelevant weight at step $n$ taken with respect to the noise applied at all times, $\langle w_{\mathrm{irrel}}(n) \rangle_{\mathrm{all}}$, stays identical to the initial weight,

$$\langle w_{\mathrm{irrel}}(n) \rangle_{\mathrm{all}} = w_{\mathrm{irrel}}(0). \tag{S130}$$

As a consequence the empirical ensemble mean, obtained by averaging over the irrelevant weights at a step $n$ in simulations, does not drift, cf. main text, Fig. 2, and Fig. S1. In contrast, the variance of the irrelevant weight, $\langle\langle w_{\mathrm{irrel}}(n) \rangle\rangle_{\mathrm{all}}$, increases from one update to the next. This is because the variance of a weight update,

$$\begin{aligned} \langle\langle \Delta w_{\mathrm{irrel}} \rangle\rangle &= \frac{\eta^2}{\sigma_{\mathrm{WP}}^4} \left\langle \left(E^{\mathrm{pert}} - E\right)^2 \cdot (\xi_{\mathrm{irrel}}^{\mathrm{WP}})^2 \right\rangle \\ &= 2\eta^2\alpha^2(E - E_{\mathrm{opt}}) + \tfrac{1}{4}\eta^2\sigma_{\mathrm{eff}}^2\alpha^2\left(M^2 N_{\mathrm{eff}} + 2M\right), \end{aligned} \tag{S131}$$

(derived below, Eq. (S132)) is nonzero and this variance adds to the variance present before the update. Therefore the empirical standard deviation of the sample of irrelevant weights increases with each learning step $n$, cf. Fig. 2.

We have seen in SM3, Sec. "Contributions to the weight update: gradient following, credit assignment-related noise and reward noise" that the WP weight update can be split into three parts, originating from gradient following (Line (S89)), the specific way of solving the credit assignment problem (Line (S90)) and reward noise (Line (S92)). Since the variance increase in the irrelevant weight, Eq. (S131), stems from the weight update, we can attribute its different terms to the different weight update contributions. For this we first note that the gradient following contribution line Eq. (S89) does not influence $\Delta w_{\mathrm{irrel}}$, since it is parallel to the error gradient (Eq. (S93)), which lies in the subspace of relevant weights. Since the variances due to independent noise sources add up, the variance of an irrelevant weight's update (Eq. (S131)) is the sum of the variances of credit assignment-related (Eq. (S102)) and reward noise induced fluctuations (Eq. (S109))

$$\begin{aligned} \langle\langle \Delta w_{\mathrm{irrel}} \rangle\rangle &= \langle\langle \delta w_{\mathrm{irrel}}^{\mathrm{cr.as}} \rangle\rangle + \langle\langle \delta w_{\mathrm{irrel}}^{\mathrm{rew.noise,quad}} \rangle\rangle \\ &= \eta^2 \left|\frac{\partial E}{\partial w}\right|^2 + \frac{1}{4}\eta^2\sigma_{\mathrm{eff}}^2\alpha^2\left(M^2 N_{\mathrm{eff}} + 2M\right). \end{aligned} \tag{S132}$$

Noticing that $\left|\frac{\partial E}{\partial w}\right|^2 = \mathrm{tr}[W S^2 W^T] = 2\alpha^2(E - E_{\mathrm{opt}})$ yields Eq. (S131).

The derivation shows that the first term in Eq. (S131) originates from credit assignment noise while the second originates from reward noise. Consequently, the first term decreases with the learning progress (as it depends on $E$) and does not depend on the perturbation strength $\sigma^2_{\mathrm{WP}}$. It dominates at the beginning of the training, when the error is large. The second term is, in contrast, constant during learning and depends on the perturbation strength $\sigma^2_{\mathrm{WP}}$. For finite $\sigma^2_{\mathrm{WP}}$ it becomes influential at the end of training when the error is small. The first term in Eq. (S131) converges for finite $\sigma^2_{\mathrm{WP}}$ to a finite value since the second term implies that the task performance error $E$ never converges to zero (Eq. (S58)). While the relevant weights then do not improve further and just fluctuate around their optimal values, the irrelevant weights keep diffusing (Fig. S1b, left hand side, and Sec. "Weight diffusion for stationary error in presence of reward noise" below). For infinitesimal perturbation strength, the second term in Eq. (S131) is zero and the error $E$ as well as the first term converge to zero. The relevant weights converge and the diffusion of irrelevant weights is only transient (Fig. S1a, right hand side, and the next Sec. "Transient weight diffusion due to credit assignment-related update fluctuations").

#### Transient weight diffusion due to credit assignment-related update fluctuations

In the following we quantitatively analyze the transient growth of weights for infinitesimal perturbation size $\sigma^2_{\mathrm{WP}} \to 0^+$. This will explain the asymptotic agreement of the standard deviation of the ensemble of irrelevant weights and the target weight size in Fig. S1a, left hand side.

Summing the variances that arise at every step from Eq. (S131) yields after $n$ steps a total additional variance

$$\langle\langle \Delta w^{\mathrm{tot}}_{\mathrm{irrel}}(n)\rangle\rangle_{\mathrm{all}} = 2\eta^2\alpha^2 \cdot (E(0) - E_{\mathrm{opt}}) \sum_{m=0}^{n-1} a^m = \frac{2\eta^2\alpha^2}{1-a}(E(0) - E_{\mathrm{opt}}) \cdot (1 - a^n). \tag{S133}$$

Here $\Delta w^{\mathrm{tot}}_{\mathrm{irrel}}(n)$ denotes the total change in the irrelevant weight up to the $n$th step and we used that $\langle E(m)\rangle_{\mathrm{all}} - E_{\mathrm{opt}} = (E(0) - E_{\mathrm{opt}})a^m$ for $\sigma^2_{\mathrm{WP}} \to 0^+$ (Eqs. (S61,S63)).

We can now compare the standard deviation of the irrelevant weights to the relevant weights' drift towards their targets. For this we first note that $E(0)(1 - a^n) = E(0) - \langle E(n)\rangle_{\mathrm{all}}$ and that the error at a learning step (measured against $E_{\mathrm{opt}}$) is proportional to the 2-norm of the relevant weight mismatch, $E - E_{\mathrm{opt}} = \frac{1}{2}\alpha^2 \sum_{i=1}^{M} \sum_{j=1}^{N_{\mathrm{eff}}} W^2_{\mathrm{rel},ij}$, due to our assumption on the $S$-matrix (SM1, Sec. "Task setting"). Introducing the average of squared mismatch of the relevant weights, $\overline{W^2_{\mathrm{rel}}} = (MN_{\mathrm{eff}})^{-1} \sum_{i=1}^{M} \sum_{j=1}^{N_{\mathrm{eff}}} W^2_{\mathrm{rel},ij}$, we have $E(0) - \langle E(n)\rangle_{\mathrm{all}} = \frac{1}{2}\alpha^2 MN_{\mathrm{eff}} \left(\overline{W^2_{\mathrm{rel}}}(0) - \langle\overline{W^2_{\mathrm{rel}}}(n)\rangle_{\mathrm{all}}\right)$ and Eq. (S133) becomes

$$\langle\langle \Delta w^{\mathrm{tot}}_{\mathrm{irrel}}(n)\rangle\rangle_{\mathrm{all}} = c(\eta) \cdot \left(\overline{W^2_{\mathrm{rel}}}(0) - \langle\overline{W^2_{\mathrm{rel}}}(n)\rangle_{\mathrm{all}}\right). \tag{S134}$$

The proportionality factor $c(\eta)$ is approximately 1 at the optimal learning rate (Eq. (S127)) such that reductions in the mean square mismatch of the relevant weights co-occur with equally strong increases in the variance of each irrelevant weight; in particular the standard deviation of an irrelevant weight converges to the initial root mean squared error of the weights,

$$\sqrt{\langle\langle \Delta w^{\mathrm{tot}}_{\mathrm{irrel}}(n)\rangle\rangle_{\mathrm{all}}} \to \left(\overline{W^2_{\mathrm{rel}}}(0)\right)^{1/2} \quad \text{for } n \to \infty. \tag{S135}$$

The same then holds for the empirical standard deviation of an ensemble of irrelevant weights in a simulation.

In Fig. S1a left hand side, the learning rate is optimal, $\eta = \eta^*$; further we have $w_{\mathrm{rel},ij}(0) = 0$, $w_{\mathrm{irrel},ij}(0) = 0$ and the teacher weight strengths are all identical. Thus $\left(\overline{W^2_{\mathrm{rel}}}(0)\right)^{1/2}$ equals the teacher weight strengths. Eq. (S135) then implies that the standard deviation of the ensemble of irrelevant weights converges to the teacher weights, like the relevant weights $w_{\mathrm{rel},ij}(n)$ do. In Fig. S1a, right hand side, the learning rate is decreased by a factor of 0.1, which slows the convergence of the relevant weights down. It also decreases the proportionality factor $c(\eta)$ in Eq. (S134) (Eq. (S127)) and leads to a by a factor $\sqrt{c(\eta)} < 1$ smaller spread of irrelevant weights.

#### Weight diffusion for stationary error in presence of reward noise

In Fig. S1b we consider WP's weight diffusion for finite perturbation size when the final steady state error $E_{f,\mathrm{unr}} = E_f + E_{\mathrm{opt}}$ (Eqs. (S125,S61) has been reached. This minimizes the influence of the first term in Eq. (S131), since the error is minimal, and it renders the strength of the credit assignment-related update fluctuations approximately constant, since the error is approximately constant. Because the reward noise-related update fluctuations are constant as well, we have a diffusion of irrelevant weights with constant strength: The variance of the weight distribution increases in each trial by the constant specified by Eq. (S131) with $E - E_{\mathrm{opt}} = E_f$ (Eq. (S125)),

$$\begin{aligned}\langle\langle \Delta w_{\mathrm{irrel}}\rangle\rangle &\approx 2\eta^2\alpha^2 \cdot \tfrac{1}{8}\eta^2\sigma^2_{\mathrm{WP}}\alpha^6 \frac{M^3N^3_{\mathrm{eff}}}{1-a} + \tfrac{1}{4}\eta^2\sigma^2_{\mathrm{WP}}\alpha^4 \cdot M^2N^2_{\mathrm{eff}} \\ &= \left(c(\eta) + 1\right) \cdot \tfrac{1}{4}\eta^2\sigma^2_{\mathrm{eff}}\alpha^2 \cdot M^2N_{\mathrm{eff}}.\end{aligned} \tag{S136}$$

The (with respect to trial number and weight strength) homogeneous random walk or diffusion is reflected in a square root growth of the standard deviation of the sampled weight distribution in Fig. S1b, left hand side.

#### Networks with weight decay

In biological neural networks synaptic strengths do not diverge but stay finite. As a proof of principle, we show that multiplicative weight decay can achieve this: After each update all weights are scaled down by a factor $\gamma_{\mathrm{wd}} < 1$,

$$w_{ij}(n) = \gamma_{\mathrm{wd}} \cdot \left(w_{ij}(n-1) + \Delta w_{ij}(n-1)\right). \tag{S137}$$

Fig. S1b, right hand side, illustrates weight evolution incorporating this weight decay, after the relevant weights have reached their steady state distribution (now shifted towards smaller weights). The irrelevant weights still initially diffuse; the multiplicative weight decay, like a restoring force, counteracts and finally balances the diffusion, which leads to a steady state. We note that multiplicative weight decay is invariant with respect to rotations of the input space and thus compatible with our assumption that only the first $n_{\mathrm{eff}}$ inputs are nonzero (SM3, Sec. "Task relevant and irrelevant weights"). Additive weight decay, i.e. reducing (the amplitude of) each weight by the same amount, would non-isotropically bias the weight evolution.

We further note that multiplicative weight decay is on average equivalent to L2 regularization. To implement such a regularization, the error function may be modified to $E \rightarrow E + \beta \frac{1}{2} \operatorname{tr}[ww^T]$ with a parameter $\beta$ specifying the regularization strength. This adds $-\beta w$ to the negative error gradient, which WP follows on average. Additive weight decay would on average be equivalent to an L1 regularization.

## SM 5

### Multiple subtasks

In this part we first describe the detailed setup of our tasks that are composed of multiple subtasks. Thereafter we derive the convergence factors for the learning of multiple subtasks, which were introduced in the main text using intuitive arguments, and the final error due to finite perturbations and unrealizable target components. This allows us to explain why batch learning improves the convergence speed of WP but not of NP learning. The subsequent section shows that batch learning also reduces the contribution of unrealizable target components to the final error of WP but not NP. Further we find that splitting the task into subtasks decreases the final error for learning with finite size perturbations for both WP and NP learning, while batch learning increases it.

##### Task setting and construction of subtasks

We adapt the framework of the analytically tractable tasks (main text, Sec. "Theoretical analysis") to tasks that consist of several subtasks. A task has input dimensionality $N_{\text{eff}}^{\text{task}}$, i.e. there are $N_{\text{eff}}^{\text{task}}$ latent inputs. The latent inputs have the same strength. In each trial a subtask of the task with dimensionality $N_{\text{eff}}^{\text{trial}}$ is presented. The error is then computed according to Eq. (8) and the updates according to Eq. (3) or Eq. (6).

Without loss of generality we assume rotated inputs. This allows to assign to each of the first $N_{\text{eff}}^{\text{task}}$ input neurons a different $T$-dimensional basis function $e_{jt}$; the $N - N_{\text{eff}}^{\text{task}}$ remaining inputs are zero. Different basis functions are orthogonal, $\frac{1}{T}\sum_{t=1}^{T} e_{jt}e_{kt} = \delta_{jk}$. In a given trial, $N_{\text{eff}}^{\text{trial}}$ of the first $N_{\text{eff}}^{\text{task}}$ inputs are chosen randomly with equal probability to be active,

$$r_{jt} = \begin{cases} \alpha \cdot e_{jt} & \text{if input } j \text{ is active} \\ 0 & \text{if input } j \text{ is inactive.} \end{cases} \tag{S138}$$

Active inputs therefore have strength $\alpha^2$. For simplicity we assume that the targets are fully realizable. The inputs that are active in a subtask $p$ therefore determine the targets via target weights $w^*$,

$$z_{it}^{p,*} = \sum_{j\in\Omega_p} w_{ij}^{*} r_{jt}^{p}, \tag{S139}$$

where $\Omega_p$ is the set of inputs that are active during subtask $p$. We refer to the inputs $r_{jt}^{p}$ of subtask $p$ as an "input pattern".

We define the error of the task as the average error over all subtasks, $E^{\text{task}} \equiv \langle E\rangle_{\text{subtasks}}$. Introducing the subtask-averaged correlation matrix $S^{\text{task}} \equiv \langle S\rangle_{\text{subtasks}}$, $E^{\text{task}}$ can be expressed by

$$E^{\text{task}} = \langle E\rangle_{\text{subtasks}} = \langle \tfrac{1}{2}\,\mathrm{tr}\left[W S W^{T}\right]\rangle_{\text{subtasks}} = \tfrac{1}{2}\,\mathrm{tr}\left[W S^{\text{task}} W^{T}\right]. \tag{S140}$$

An input $j$ is in a given trial active with constant probability

$$\frac{N_{\text{eff}}^{\text{trial}}}{N_{\text{eff}}^{\text{task}}} \equiv \frac{1}{P}, \tag{S141}$$

where $P$ can be interpreted as the effective number of subtasks or patterns in the task and as the number of trials needed to gather information on all relevant weights. The first $N_{\text{eff}}^{\text{task}}$ diagonal elements of $S^{\text{task}}$, which are nonzero, are therefore given by $\frac{1}{P}\alpha^2$, that is for each task-relevant input by the product of its strength with its probability of being active. This stays true if the subtasks are not random but fixed such that their sets of active inputs are non-overlapping, i.e. such that each input $j \in 1, \dots, N_{\text{eff}}^{\text{task}}$ is nonzero in exactly one subtask. $P$ is then the number of these subtasks.

The typical timescale of learning, $1/(1-a)$ (motivated after Eq. (S124)), is larger than $P$ by a factor of $MN_{\text{eff}}^{\text{task}} + 2$ for WP and $MN_{\text{eff}}^{\text{trial}} + 2$ for NP, Eq. (15) and (16). Thus for typical task dimensions all inputs are often sampled before the error significantly changes. It is therefore not important whether random or fixed input patterns are presented and whether the presentation is in a specific order. Further, similar to the case without subtasks, it is not necessary to assume a fixed set of basis functions $e_{jt}$ for our results to hold: The error computation and the update rules Eqs. (8,S30,S42) imply that only the weight mismatch and the correlation structure $S(n)$ of the inputs at a trial $n$ matter for the error between student and teacher output as well as for the weight updates. In the present part we could thus also construct each trial from completely different basis functions $\tilde{e}_{jt}(n+1) \neq \tilde{e}_{jt}(n)$, as long as their correlations remain unchanged, $\frac{1}{T}\sum_{t=1}^{T} \tilde{e}_{jt}\tilde{e}_{kt} = \delta_{jk}$ (and the targets are adapted according to Eq. (S139)).

#### Derivation of the convergence factor

Here we derive the convergence factor of WP by considering its improvements on the current subtask and the simultaneous deterioration on all other subtasks; a slightly altered derivation holds for NP. For simplicity and without loss of generality (see the related argument at the end of the previous section) we assume that there are $P$ fixed subtasks that are orthogonal in the sense that their sets of active inputs do not overlap. This implies that also their sets of trial-relevant weights do not overlap. As here we derive only the convergence factor, we also assume infinitesimally small perturbations and realizable targets.

The task error at trial $n$, Eq. (S140), is the average of the errors $E_q$ of the subtasks $q$,

$$E^{\text{task}} = \frac{1}{P}\sum_{q=1}^{P} E_q(n). \tag{S142}$$

Let pattern $p$ be presented at trial $n$. Then the error signal $E_p^{\text{pert}}(n) - E_p(n)$ and the update $\Delta w(n)$ depend only on pattern $p$, while the effect of the weight update on the task error depends on all subtasks $q$. After the update, the expected error for subtask $p$ decreases by the same factor as for the single pattern case,

$$\langle E_p(n+1)\rangle = \left(1 - 2\eta\alpha^2 + \eta^2\alpha^4(MN_{\text{eff}}^{\text{trial}} + 2)\right)\cdot\langle E_p(n)\rangle \tag{S143}$$

(Eq. (S62)), while each weight that is irrelevant to the current subtask receives fluctuations with mean zero and variance

$$\langle\langle \Delta w_{\text{tr.irrel.}}(n)\rangle\rangle = \langle\langle \delta w_{\text{tr.irrel.}}^{\text{cr.as}}(n)\rangle\rangle = 2\eta^2\alpha^2\cdot E_p(n) \tag{S144}$$

(Eq. (S131) in the small $\sigma_{\text{NP}}^2$ limit). Correspondingly, the expected error for subtasks $q \neq p$ increases as

$$\begin{aligned}
\langle E_q(n+1)\rangle &= \tfrac{1}{2}\langle \text{tr}[(W + \delta w^{\text{cr.as}})S(q)(W + \delta w^{\text{cr.as}})^T]\rangle \\
&= \langle E_q(n)\rangle + \tfrac{1}{2}\langle \text{tr}[\delta w^{\text{cr.as}}S(q)(\delta w^{\text{cr.as}})^T]\rangle \\
&= \langle E_q(n)\rangle + \tfrac{1}{2}MN_{\text{eff}}^{\text{trial}}\alpha^2\cdot\langle\langle \delta w_{\text{irrel}}^{\text{cr.as}}(n)\rangle\rangle \\
&= \langle E_q(n)\rangle + MN_{\text{eff}}^{\text{trial}}\cdot\eta^2\alpha^4\cdot\langle E_p(n)\rangle.
\end{aligned} \tag{S145}$$

Inserting Eqs. (S143,S145) into Eq. (S142) yields the evolution of the task error,

$$\begin{aligned}
\langle E^{\text{task}}(n+1)\rangle &= \frac{1}{P}\cdot\left(1 - 2\eta\alpha^2 + \eta^2\alpha^4(MN_{\text{eff}}^{\text{trial}} + 2)\right)\cdot\langle E_p(n)\rangle \\
&\quad + \frac{1}{P}\sum_{q\neq p}\left(\langle E_q(n)\rangle + MN_{\text{eff}}^{\text{trial}}\cdot\eta^2\alpha^4\cdot\langle E_p(n)\rangle\right) \\
&= E^{\text{task}}(n) + \left(\frac{-2\eta\alpha^2}{P} + \frac{\eta^2\alpha^4(PMN_{\text{eff}}^{\text{trial}} + 2)}{P}\right)\cdot\langle E_p(n)\rangle.
\end{aligned} \tag{S146}$$

Assuming that $\langle E_p(n)\rangle \approx E^{\text{task}}$, i.e. that the error on all subtasks is sufficiently similar, and using $PN_{\text{eff}}^{\text{trial}} = N_{\text{eff}}^{\text{task}}$ (Eq. (S141)) finally yields

$$\langle E^{\text{task}}(n+1)\rangle = \underbrace{\left(1 - \frac{2\eta\alpha^2}{P} + \frac{\eta^2\alpha^4(MN_{\text{eff}}^{\text{task}} + 2)}{P}\right)}_{=a_{\text{WP}}}\cdot\langle E^{\text{task}}(n)\rangle. \tag{S147}$$

The factor $1/P$ arises because the changes to weights relevant in any subtask affect the error only in $1/P$ of the trials. Apart from that factor, the beneficial part of the convergence factor due to gradient following, $-\frac{1}{P}\cdot 2\eta\alpha^2$, stays the same as for a single subtask. The last part due to update fluctuations in all task-relevant weights, $\frac{1}{P}\cdot\eta^2\alpha^4(MN_{\text{eff}}^{\text{task}} + 2)$, however, increases approximately by a factor of $P$ in relation to the beneficial part, as the number of fluctuating weights relevant for the task is $P$ times larger than if only subtask $p$ had to be learnt. These are the arguments given in the main text.

Minimizing $a_{\text{WP}}$ with respect to $\eta$ yields Eq. (16),

$$\eta_{\text{WP}}^* = \frac{1}{(MN_{\text{eff}}^{\text{task}} + 2)\alpha^2}, \qquad a_{\text{WP}}^* = 1 - \frac{1}{P}\frac{1}{MN_{\text{eff}}^{\text{task}} + 2}. \tag{S148}$$

For NP, the same derivation with $\langle E_q(n+1)\rangle = \langle E_q(n)\rangle$ for $q \neq p$ instead of Eq. (S145) produces Eq. (15),

$$\eta_{\text{NP}}^* = \frac{1}{(MN_{\text{eff}}^{\text{trial}} + 2)\alpha^2}, \qquad a_{\text{NP}}^* = 1 - \frac{1}{P}\frac{1}{MN_{\text{eff}}^{\text{trial}} + 2}. \tag{S149}$$

#### Derivation of the final error due to finite perturbations

In this section we quantify the final error in a task with multiple subtasks that results from quadratic reward noise-induced update fluctuations. The contribution of unrealizable target components will be investigated in the next section.

We consider the already described setting with a task that has effective input dimension $N_{\text{eff}}^{\text{task}}$ and is split into $P$ non-overlapping subtasks. These have effectively $N_{\text{eff}}^{\text{trial}}$-dimensional inputs and are presented in the different learning trials. The final task error due to finite perturbation sizes may be defined as

$$E_f^{\text{task}} = \frac{1}{P}\sum_{q=1}^{P} E_{f,q}, \tag{S150}$$

where $E_{f,q}$ are the final errors of the individual subtasks, which arise due to quadratic reward noise (Eq. (S124)). We investigate how $E_f^{\text{task}}$ depends on the number $P$ of subtasks in which the task is split, where $P = 1$ corresponds to the single-pattern case Eqs. (S125,S126).

Combining Eqs. (S150,S124) and line (S118), the contribution of (quadratic) reward noise-induced update fluctuations to the final error is

$$E_f = \frac{1}{1-a}\cdot\frac{1}{P}\sum_{q=1}^{P}\tfrac{1}{2}\left\langle \mathrm{tr}[\delta w^{\text{rew.noise,quad}} S[q] (\delta w^{\text{rew.noise,quad}})^T]\right\rangle . \tag{S151}$$

Because all subtasks have identical properties, we can consider the effect that an update following the presentation of an arbitrary subtask $p$ has on the errors $E_q$ of all subtasks $q$.

For WP, all weights are updated and fluctuations of the weights relevant for any subtask $q$ (including $q = p$) are equal in size and statistics. Inserting the expressions for $\langle\langle \delta w_{ij}^{\text{rew.noise,quad}}\rangle\rangle$, $\eta^*$ and $a^*$ (Eqs. (S109,S148)) into Eq. (S151) yields

$$\begin{aligned}
E_f^{\text{WP}} &= \frac{1}{1-a^*}\cdot\frac{\alpha^2}{2P}\sum_{i=1}^{M}\sum_{j=1}^{N_{\text{eff}}^{\text{task}}}\left\langle\left(\delta w_{ij}^{\text{rew.noise,quad}}\right)^2\right\rangle \\
&\approx PMN_{\text{eff}}^{\text{task}}\cdot\frac{\alpha^2}{2P}MN_{\text{eff}}^{\text{task}}\cdot\frac{1}{4}(\eta^*)^2\sigma_{\text{eff}}^2\alpha^2M^2N_{\text{eff}}^{\text{trial}} \\
&\approx PMN_{\text{eff}}^{\text{task}}\cdot\frac{1}{(MN_{\text{eff}}^{\text{task}}\alpha^2)^2}\cdot\frac{1}{8}\sigma_{\text{eff}}^2\alpha^4\frac{M^3(N_{\text{eff}}^{\text{task}})^2}{P^2} \\
&= \frac{1}{8}\sigma_{\text{eff}}^2M^2N_{\text{eff}}^{\text{trial}} = \frac{1}{8}\sigma_{\text{eff}}^2M^2\frac{N_{\text{eff}}^{\text{task}}}{P}.
\end{aligned} \tag{S152}$$

The final error of WP thus becomes smaller if the task is split into more subtasks, even though that means that WP converges more slowly (Eq. (S148)). The reason is that WP has to operate at a roughly $P$ times smaller learning rate, which means that update fluctuations average out over more trials, ultimately lowering the final error.

For NP, only weights relevant for the current subtask are updated such that only the term with $q = p$ in Eq. (S151) contributes. Inserting the expressions for $\langle\langle \delta w_{ij}^{\text{rew.noise,quad}}\rangle\rangle$, $\eta^*$ and $a^*$ (Eqs. (S110,S149)) into Eq. (S151) yields

$$\begin{aligned}
E_f^{\text{NP}} &= \frac{1}{1-a}\cdot\frac{\alpha^2}{2P}\sum_{i=1}^{M}\sum_{j\in\Omega_p}\left\langle\left(\delta w_{ij}^{\text{rew.noise,quad}}\right)^2\right\rangle \\
&\approx PMN_{\text{eff}}^{\text{trial}}\cdot\frac{\alpha^2}{2P}MN_{\text{eff}}^{\text{trial}}\cdot\frac{1}{4}\eta^2\sigma_{\text{eff}}^2\alpha^2M^2T \\
&\approx MN_{\text{eff}}^{\text{task}}\cdot\frac{1}{(MN_{\text{eff}}^{\text{trial}}\alpha^2)^2}\cdot\frac{1}{8}\sigma_{\text{eff}}^2\alpha^4M^3\frac{N_{\text{eff}}^{\text{task}}T}{P^2} \\
&= \frac{1}{8}\sigma_{\text{eff}}^2M^2T = \frac{1}{8}\sigma_{\text{eff}}^2M^2\frac{T^{\text{task}}}{P},
\end{aligned} \tag{S153}$$

where $\Omega_p$ is the set of inputs that are active during subtask $p$, $T$ is the subtask duration and $T^{\text{task}}$ the duration of the full task with all subtasks concatenated. We conclude that if subtasks are obtained by splitting an original, full task of duration $T^{\text{task}}$ into $P$ subtasks such that $T = T^{\text{task}}/P$, $E_f^{\text{NP}}$ scales like $E_f^{\text{WP}}$ with $1/P$. This holds despite the fact that the optimal learning rate of NP stays approximately constant when changing $P$. Since for any type of split $T \geq N_{\text{eff}}^{\text{trial}}$, comparison of Eq. (S153) and Eq. (S152) yields $E_f^{\text{NP}} \geq E_f^{\text{WP}}$.

#### Derivation of the final error due to unrealizable target components

Here we obtain the final error $E_{f,d}$ that results from the target components of a trial that are perpendicular to any of the inputs, i.e. from the *trial-unrealizable* target components. This error only occurs for NP.

We assume that in each trial the trial-unrealizable components have the same strength, leading to an unavoidable limiting error $E_{\mathrm{opt}}^{\mathrm{tr.unr.}}$, which is the same for each subtask. Since the error definition Eq. (S150) is normalized by $P$, $E_{\mathrm{opt}}^{\mathrm{tr.unr.}}$ is independent of $P$. For NP but not WP, trial-unrealizable target components add reward noise $\Delta E_{\mathrm{pert}}^{\mathrm{lin,unr}}$ to the error signal (Eq. (S80)), which gets translated into update fluctuations with $\langle\langle \delta w_{ij}^{\mathrm{rew.noise,lin}} \rangle\rangle = 2\eta^2\alpha^2 E_{\mathrm{opt}}^{\mathrm{tr.unr.}}$ (Eq. (S107)). These results are still valid for the case considered here, with $\delta w_{ij}^{\mathrm{rew.noise,lin}} \neq 0$ only for the $MN_{\mathrm{eff}}^{\mathrm{trial}}$ trial-relevant weights of the current subtask $p$. The update fluctuations lead to an expected increase in error (Eqs. (S117,S142)) of

$$
\begin{aligned}
b_{\mathrm{NP}}^{\mathrm{lin}} &= \frac{1}{P}\sum_{q=1}^{P} \tfrac{1}{2}\langle \mathrm{tr}[\overbrace{\delta w^{\mathrm{rew.noise,lin}} S_q (\delta w^{\mathrm{rew.noise,lin}})^T}^{\neq 0 \text{ only for } q=p}]\rangle \\
&= \frac{\alpha^2}{2P}\sum_{q=1}^{P}\delta_{qp}\sum_{i=1}^{M}\sum_{j=1}^{N_{\mathrm{eff}}^{\mathrm{trial}}}\langle\langle \delta w_{ij}^{\mathrm{rew.noise,lin}}\rangle\rangle \\
&= \frac{\alpha^2}{2P} MN_{\mathrm{eff}}^{\mathrm{trial}} \cdot 2\eta^2\alpha^2 E_{\mathrm{opt}}^{\mathrm{tr.unr.}} = \frac{1}{P}\eta^2\alpha^4 MN_{\mathrm{eff}}^{\mathrm{trial}} \cdot E_{\mathrm{opt}}^{\mathrm{tr.unr.}}. 
\end{aligned} \tag{S154}
$$

The result is very similar to $b_{\mathrm{NP}}^{\mathrm{lin}}$ in Eq. (S67) and reproduces it for $P = 1$. When keeping $N_{\mathrm{eff}}^{\mathrm{task}}$ fixed and regarding $P$ as a free parameter, both $\eta$ (relative to $\eta^*$) and $N_{\mathrm{eff}}^{\mathrm{trial}}$ contain implicit dependencies on $P$. The contribution of $b_{\mathrm{NP}}^{\mathrm{lin}}$ to the final error (Eq. (S124)), at the optimal learning rate (Eq. (S149)), is

$$
\begin{aligned}
E_{f,d}(\eta^*) &\overset{\mathrm{NP}}{=} \frac{b_{\mathrm{NP}}^{\mathrm{lin}}(\eta^*)}{1 - a_{\mathrm{NP}}(\eta^*)} = \frac{1}{P}(\eta^*)^2\alpha^4 \frac{MN_{\mathrm{eff}}^{\mathrm{trial}}}{1 - a_{\mathrm{NP}}(\eta^*)} \cdot E_{\mathrm{opt}}^{\mathrm{tr.unr.}} \\
&= \frac{1}{P}\frac{\alpha^4}{(MN_{\mathrm{eff}}^{\mathrm{trial}} + 2)^2\alpha^4}(MN_{\mathrm{eff}}^{\mathrm{trial}})P(MN_{\mathrm{eff}}^{\mathrm{trial}} + 2) \cdot E_{\mathrm{opt}}^{\mathrm{tr.unr.}} \\
&\approx E_{\mathrm{opt}}^{\mathrm{tr.unr.}}.
\end{aligned} \tag{S155}
$$

The final error contribution $E_{f,d}$ due to trial-unrealizable target components is thus (approximately) independent of the number of subtasks $P$ that the full task is split into. Also, at the optimal learning rate, the additional error contribution for NP learning again equals the unavoidable limiting error $E_{\mathrm{opt}}^{\mathrm{tr.unr.}}$ (cf. Eq. (S128)).

The results of this section only hold for trial-unrealizable target components. In Sec. “Batch learning improves WP’s but not NP’s performance for overlapping subtasks and unrealizable target components” we make the distinction between “trial-unrealizable” and “task-unrealizable” components. There we show that trial-realizable but task-unrealizable components also harm WP learning, but that combining trials into batches renders some task-unrealizable components also trial-unrealizable, reducing the effect.

#### Error curves when learning multiple subtasks

Combining Eqs. (S148,S149,S152,S153,S155,S124), the errors of WP and NP evolve as

$$
\langle E(n)\rangle = \left(E(0) - E_{f,\mathrm{unr}}\right)\cdot a^n + E_{f,\mathrm{unr}} \tag{S156}
$$

to a final error $E_{f,\mathrm{unr}} = E_f + E_{f,d} + E_{\mathrm{opt}}^{\mathrm{tr.unr.}}$ with

$$
a_{\mathrm{WP}}^* = 1 - \frac{1}{P}\frac{1}{MN_{\mathrm{eff}}^{\mathrm{task}} + 2}, \qquad a_{\mathrm{NP}}^* = 1 - \frac{1}{P}\frac{1}{MN_{\mathrm{eff}}^{\mathrm{trial}} + 2}, \tag{S157}
$$

$$
E_f^{\mathrm{WP}}(\eta^*) \approx \frac{1}{8}\sigma_{\mathrm{eff}}^2 M^2 N_{\mathrm{eff}}^{\mathrm{trial}}, \qquad E_f^{\mathrm{NP}}(\eta^*) \approx \frac{1}{8}\sigma_{\mathrm{eff}}^2 M^2 T, \tag{S158}
$$

$$
E_{f,d}^{\mathrm{WP}}(\eta^*) = 0, \qquad E_{f,d}^{\mathrm{NP}}(\eta^*) \approx \frac{1}{P}\cdot E_{\mathrm{opt}}^{\mathrm{tr.unr.}}. \tag{S159}
$$

#### Batch learning improves WP's but not NP's performance

Why does batch learning improve WP's but not NP's performance? In the following we obtain intuitive explanations from observing how concatenating trials into batches affects which weights are relevant and which target components are realizable in a trial.

Concatenating $K \leq P$ subtasks with non-overlapping inputs into a single batch changes the subtask parameters as follows

$$N_{\text{eff}}^{\text{trial}} \to K N_{\text{eff}}^{\text{trial}}, \qquad P \to \frac{P}{K}, \qquad \alpha^2 \to \frac{\alpha^2}{K}, \tag{S160}$$

$$N_{\text{eff}}^{\text{task}} \to N_{\text{eff}}^{\text{task}}, \qquad T \to KT. \tag{S161}$$

Here the change Eq. (S160) left reflects that the input dimensionality of $K$ non-overlapping inputs with dimension $N_{\text{eff}}^{\text{trial}}$ is $K N_{\text{eff}}^{\text{trial}}$; consequently, the number of trial-relevant weights becomes $KMN_{\text{eff}}^{\text{trial}}$. The number of trials needed to gather information on all task-relevant weights, $P$, therefore decreases, by a factor of $K$ (Eq. (S160) middle). The temporal extent of the subtask increases by a factor $K$ (Eq. (S161) middle). The individual input vectors keep their nonzero entries and are padded by zero entries from length $T$ to length $KT$. Because the entries of $S$ have as normalizing prefactor the new total trial duration $KT$ instead of $T$, the nonzero entries read $\alpha^2/K$ (Eq. (S160) right). Since there are $K$ times more nonzero input vectors, the total input strength per time step, $\alpha_{\text{tot}}^2$ (Eq. (S11)), remains unchanged, as we expect it from a concatenation operation. The task dimensionality is unaffected by changes of the subtasks (Eq. (S161) left).

Using these scalings and assuming $MN_{\text{eff}}^{\text{trial}} \gg 2$ in Eqs. (S148,S149) shows that, approximately, the optimal learning rate and the convergence rate ($\approx 1-a$) of WP increase linearly with $K$, while the performance of NP stays unaffected,

$$\eta_{\text{WP}}^* \to \frac{1}{(MN_{\text{eff}}^{\text{task}}+2)\frac{\alpha^2}{K}} = K \cdot \eta_{\text{WP}}^*, \qquad 1-a_{\text{WP}}^* \to \frac{K}{P}\frac{1}{MN_{\text{eff}}^{\text{task}}+2} = K \cdot (1-a_{\text{WP}}^*), \tag{S162}$$

$$\eta_{\text{NP}}^* \to \frac{1}{(MKN_{\text{eff}}^{\text{trial}}+2)\frac{\alpha^2}{K}} \approx \eta_{\text{NP}}^*, \qquad 1-a_{\text{NP}}^* \to \frac{K}{P}\frac{1}{MKN_{\text{eff}}^{\text{trial}}+2} \approx 1-a_{\text{NP}}^*. \tag{S163}$$

The result can be understood as follows: In a batch of $K$ subtasks, the error feedback of a single trial contains information on $K$ times more weights. In WP this increases the number of weights that receive beneficial updates (Eq. (S143)) and do not simply fluctuate (Eq. (S145)), by a factor $K$. Therefore learning is $K$ times faster. Since for NP trial-irrelevant weights do not fluctuate, it does not matter whether the weight update information is presented in one subtask or distributed over different ones. Therefore NP learning does not benefit from forming batches.

The scaling of the final error due to finite perturbation sizes when learning at the optimal learning rate can be obtained using Eqs. (S152,S153). If we concatenate $K$ non-overlapping subtasks of duration $T$ into batches, $N_{\text{eff}}^{\text{trial}} \to KN_{\text{eff}}^{\text{trial}}$ (Eq. (S160)) and $T \to KT$ (Eq. (S161)) imply

$$E_f^{\text{WP}} = \frac{1}{8}\sigma_{\text{eff}}^2 M^2 N_{\text{eff}}^{\text{trial}} \cdot K, \qquad E_f^{\text{NP}} = \frac{1}{8}\sigma_{\text{eff}}^2 M^2 T \cdot K, \tag{S164}$$

i.e. the final error increases proportional to the batch size. Fig. S5 shows WP and NP learning for different batch sizes along with predicted error curves from the results of this section. The independence of the final error in the MNIST task from the perturbation strength (Fig. S10) and for NP from the batch size (Fig. 7) suggests that final performance is not limited by the reward noise caused by finite perturbations.

#### Batch learning improves WP's but not NP's performance for overlapping subtasks and unrealizable targets

Are there specific effects of batch learning benefiting WP and/or NP when the subtask input patterns are overlapping and the targets contain unrealizable components? To address this questions we need to distinguish *trial-unrealizable* and *task-unrealizable* target components. We will show that trial-realizable but task-unrealizable components are a source of *gradient noise* for WP and NP, that larger batch sizes render some of these components trial-unrealizable, and that this reduces the gradient noise for WP but not NP.

Let $w_{ij}^*$ be an optimal weight matrix that minimizes the task error Eq. (S142), such that in a subtask $p$ and for *task-unrealizable* targets $d^p$ the targets $z_{it}^{p,*}$ are given by

$$z_{it}^{p,*} = \sum_{j=1}^{N} w_{ij}^* r_{jt}^p + d_{it}^p, \qquad d_{it}^p = d_{it}^{p,\text{tr.real.}} + d_{it}^{p,\text{tr.unr.}}. \tag{S165}$$

Here $d^{p,\text{tr.unr.}}$ is the *trial-unrealizable* component that is orthogonal to all inputs of subtask $p$. $d^{p,\text{tr.real.}}$ is the target component that cannot be realized without simultaneously increasing the task error due to worse performance on other (overlapping) subtasks, although it could be realized in trial $p$. We note that $d_{it}^{p,\text{tr.real.}}$ can only be nonzero if subtasks overlap. Expressing the desired output of the subtask using this distinction,

$$z_{it}^{p,*} = \sum_{j=1}^{N} w_{ij}^{*} r_{jt}^{p} + d_{it}^{p,\text{tr.real.}} + d_{it}^{p,\text{tr.unr.}}, \tag{S166}$$

the error for subtask $p$ is

$$\begin{aligned} E_p &= \frac{1}{2T}\sum_{i=1}^{M}\sum_{t=1}^{T}\left(z_{it}-z_{it}^{*}\right)^2 = \frac{1}{2T}\sum_{i=1}^{M}\sum_{t=1}^{T}\Big(\sum_{j=1}^{N} W_{ij} r_{jt}^{p} - d_{it}^{p,\text{tr.real.}} - d_{it}^{p,\text{tr.unr.}}\Big)^2 \\ &= \tfrac{1}{2}\operatorname{tr}[W S^{p} W^{T}] - \frac{1}{T}\operatorname{tr}\left[W r^{p}\left(d^{p,\text{tr.real.}}\right)^{T}\right] + \frac{1}{2T}\operatorname{tr}\left[d^{p}\left(d^{p}\right)^{T}\right]. \end{aligned} \tag{S167}$$

Here the weight mismatch $W = w - w^{*}$ is defined relative to the weights $w^{*}$ that are optimal for the full task. Perturbations of the weights can couple to $d^{p,\text{tr.real.}}$ but not $d^{p,\text{tr.unr.}}$, while node perturbations project equally onto trial- and task-unrealizable target components. $d_{it}^{p,\text{tr.real.}}$ endows the output- and weight error gradients with noise,

$$\frac{\partial E_p}{\partial z_{it}} = \underbrace{\frac{1}{T}\sum_{j=1}^{N} W_{ij} r_{jt}^{p}}_{\text{stoch. est. of } \frac{\partial E^{[\text{task}]_{\text{real}}}}{\partial z_{it}}} - \underbrace{\frac{1}{T} d_{it}^{p,\text{tr.real.}}}_{\text{noise affecting WP and NP}} - \underbrace{\frac{1}{T} d_{it}^{p,\text{tr.unr.}}}_{\text{noise affecting only NP}}, \tag{S168}$$

$$\frac{\partial E_p}{\partial w_{ij}} = \underbrace{\sum_{k=1}^{N} W_{ik} S_{kj}^{p}}_{\text{stoch. est. of } \frac{\partial E^{[\text{task}]_{\text{real}}}}{\partial w_{ij}}} - \underbrace{\frac{1}{T}\sum_{t=1} d_{it}^{p,\text{tr.real.}} r_{jt}^{p}}_{\text{additional gradient noise}}. \tag{S169}$$

The first terms are stochastic estimates of the gradient of the full task with all unrealizable target components removed. The realizable part of the task is solved by the same weight configurations as the full, unrealizable task, as the unrealizable components by definition cannot be improved. Thus their contribution to the error is the same regardless of the weights, and following the gradient of the realizable task already solves the task. The other terms in Eqs. (S168,S169) therefore cause noise. $d^{p,\text{tr.unr.}}$ causes reward noise and harms only NP (Eq. (S155)). $d^{p,\text{tr.real.}}$ adds *gradient noise* to the weight gradient: Even if WP and NP could average over all possible perturbations and measure $\partial E_p/\partial w_{ij}$ with arbitrarily high precision, the computed weight update would not be linearly optimal. The gradient is wrong in the sense that for nonzero $d_{it}^{p,\text{tr.real.}}$ the trial error gradient $\partial E_p/\partial w_{ij}$ does not provide an optimal approximation to the error gradient of the entire task. Because $d^{p,\text{tr.real.}}$ is realizable within the single trial, both WP and NP try to reduce it. It therefore causes alike noise for both WP and NP.

As for tasks without subtasks (cf. SM3, Sec. "Realizable and unrealizable outputs"), the trial-unrealizable part $d^{p,\text{tr.unr.}}$ affects NP by inducing reward noise. WP cannot induce output perturbation along this output component and is thus not affected. Since output perturbations directly change $E_{\text{pert}} - E$ without distinguishing between realizable and unrealizable target components, $E_{\text{pert}} - E$ is equally affected by $d^{p,\text{tr.real.}}$ and $d^{p,\text{tr.unr.}}$. Therefore basically their sum $d^{p}$ matters for NP.

An increasing batch size renders some task-unrealizable but trial-realizable components also trial-unrealizable. This becomes especially apparent for the case where the batch contains all subtasks: Because there is only one (sub-)task or trial, task-unrealizable components are also trial-unrealizable, $d^{1,\text{tr.unr.}} = d^{1}$ and $d^{1,\text{tr.real.}} = 0$.

Due to the decrease of $d^{\text{tr.real.}}$ for increasing batch size, the gradient noise affecting WP decreases, as it no longer induces output perturbations along these components. In contrast, the strength $\frac{1}{T}\sum_{t=1}^{T}\sum_{i=1}^{M}(d_{it}^{p})^2$ of the sum $d^{p,\text{tr.real.}} + d^{p,\text{tr.unr.}} = d^{p}$, which determines NP's gradient noise, is independent of the batch size. Therefore WP but not NP benefits from increasing the batch size.

## SM 6

### Arbitrary input strength distributions

##### Error curves for input components with different strength

In the main text, Sec. "Theoretical analysis" and in SM2, Sec. "Error curves for equally strong input components" (as well as in the SM parts thereafter), we focused on the error dynamics in the special case where all latent inputs have the same strength. To understand the error dynamics in the reservoir computing simulation experiment and for completeness, here we also give the general solution for the error dynamics. We aim at expressions analogous to those of SM2, Sec. "Error curves for equally strong input components". To obtain them we split the error $E$ into components $E^\mu$ and replace the convergence factor $a$ by a matrix $(A+B)_{\mu\nu}$ and the per-update error increase $b$ by a vector $b_\mu$.

To define the error components, we split the correlation matrix $S$ into a sum of matrices $S^\mu$, where each matrix contains the contribution of one eigenvalue to $S$,

$$S^\mu = O D^\mu O^T, \qquad D^\mu_{jk} = \alpha_\mu^2 \delta_{\mu jk}, \qquad S = \sum_{\mu=1}^{N} S^\mu, \tag{S170}$$

where $O$ diagonalizes $S$ and $D^\mu$ is a matrix with only one nonzero element $D^\mu_{\mu\mu} = \alpha_\mu^2$. We observe that the powers of $S$ can be written as sums of powers of $S^\mu$,

$$S^2 = \sum_{\mu=1}^{N} O (D^\mu)^2 O^T = \sum_{\mu=1}^{N} (S^\mu)^2, \qquad S S^\mu = S^\mu S^\mu = \alpha_\mu^2 S^\mu, \qquad \mathrm{tr}[S] = \sum_{\mu=1}^{N} \alpha_\mu^2. \tag{S171}$$

The error $E$ then reads

$$E = \tfrac{1}{2}\,\mathrm{tr}[W \tilde{S} W^T] + E_{\mathrm{opt}} = \tfrac{1}{2}\,\mathrm{tr}[W \sum_{\mu=1}^{N} \tilde{S}^\mu W^T] + E_{\mathrm{opt}}. \tag{S172}$$

Here we again use the tilde symbol to distinguish the correlation matrix that stems from the cost evaluation at trial $n$ from the correlation matrix that determines the error change due to perturbations in trial $n-1$ (SM2). Because the trace is linear, we can split the input strength-dependent part of the error into $N$ summands $E^\mu$ as

$$E^\mu = \tfrac{1}{2}\,\mathrm{tr}[W \tilde{S}^\mu W^T] \qquad E = \sum_{\mu=1}^{N} E^\mu + E_{\mathrm{opt}}. \tag{S173}$$

$E^\mu$ only depends on the strength of the $\mu$th input component and the weight mismatch projected onto the corresponding input direction (Eq. (S170)). In other words, for each input component $\mu$ we can define related weights that read out from it, and a related error component $E^\mu$ that depends on the mismatch of these weights. We now obtain a recurrence equation that relates the $N$ error components $\langle E^\mu(n)\rangle$ at trial $n$ to those at trial $n-1$. For this we first split $\langle E^\mu(n)\rangle$ and $\langle E^\mu(n-1)\rangle$ in Eqs. (S40, S57) up using Eq. (S173). It then suffices to also split $\tilde{S}$ up using Eq. (S170) and to explicitly evaluate the powers and traces of $S$ using Eq. (S171). For WP we obtain

$$\begin{aligned}
\langle E^\mu(n)\rangle &\overset{\mathrm{WP}}{=} \langle E^\mu(n-1)\rangle - \eta\,\mathrm{tr}[W S^\mu S W^T] + \frac{\eta^2}{2} M\,\mathrm{tr}[W S^2 W^T]\,\mathrm{tr}[S^\mu] + \eta^2\,\mathrm{tr}[W S S^\mu S W^T] \\
&\quad + \frac{\eta^2 \sigma_{\mathrm{WP}}^2}{8}\big(M^3\,\mathrm{tr}[S^\mu]\,\mathrm{tr}[S]^2 + 2M^2\,\mathrm{tr}[S^\mu]\,\mathrm{tr}[S^2] + 4M^2\,\mathrm{tr}[S^\mu S]\,\mathrm{tr}[S] + 8M\,\mathrm{tr}[S S^\mu S]\big) \\
&= \langle E^\mu(n-1)\rangle - \eta\alpha_\mu^2\,\mathrm{tr}[W S^\mu W^T] + \frac{\eta^2}{2} M \sum_{\nu=1}^{N} \alpha_\mu^2 \alpha_\nu^2\,\mathrm{tr}[W S^\nu W^T] + \eta^2 \alpha_\mu^4\,\mathrm{tr}[W S^\mu W^T] \\
&\quad + \frac{\eta^2 \sigma_{\mathrm{WP}}^2}{8}\Big(M^3 \alpha_\mu^2 \Big(\sum_{\nu=1}^{N} \alpha_\nu^2\Big)^2 + 2M^2 \alpha_\mu^2 \sum_{\nu=1}^{N} \alpha_\nu^4 + 4M^2 \alpha_\mu^4 \sum_{\nu=1}^{N} \alpha_\nu^2 + 8M\alpha_\mu^6\Big) \\
&= \sum_{\nu=1}^{N} \big((1 - 2\eta\alpha_\mu^2 + 2\eta^2\alpha_\mu^4)\,\mathbb{1}_{\mu\nu} + \eta^2 \alpha_\mu^2 \alpha_\nu^2 M\big) \cdot \langle E^\nu(n-1)\rangle \\
&\quad + \tfrac{1}{8}\eta^2 \sigma_{\mathrm{WP}}^2 \cdot \Big(M^3 \alpha_\mu^2 \Big(\sum_{\nu=1}^{N} \alpha_\nu^2\Big)^2 + 2M^2 \alpha_\mu^2 \sum_{\nu=1}^{N} \alpha_\nu^4 + 4M^2 \alpha_\mu^4 \sum_{\nu=1}^{N} \alpha_\nu^2 + 8M\alpha_\mu^6\Big).
\end{aligned} \tag{S174}$$

For NP we obtain

$$
\begin{aligned}
\langle E^\mu(n)\rangle &\overset{\text{NP}}{=} \langle E^\mu(n-1)\rangle - \eta\,\mathrm{tr}[WS^\mu SW^T] + \frac{\eta^2}{2} M\,\mathrm{tr}[WSW^T]\,\mathrm{tr}[S^\mu S] + \eta^2\,\mathrm{tr}[WSS^\mu SW^T] \\
&\quad + \frac{\eta^2\sigma_{\text{NP}}^2}{8T}\mathrm{tr}[S^\mu S]\cdot\left(M^3T^2 + 6M^2T + 8M\right) + \eta^2 M\,\mathrm{tr}[S^\mu S]\cdot\frac{1}{2T}\mathrm{tr}[dd^T] \\
&= \langle E^\mu(n-1)\rangle - \eta\alpha_\mu^2\,\mathrm{tr}[WS^\mu W^T] + \frac{\eta^2}{2}\alpha_\mu^4 M\sum_{\nu=1}^N \mathrm{tr}[WS^\nu W^T] + \eta^2\alpha_\mu^4\,\mathrm{tr}[WS^\mu W^T] \\
&\quad + \frac{\eta^2\sigma_{\text{NP}}^2}{8T}\alpha_\mu^4\cdot\left(M^3T^2 + 6M^2T + 8M\right) + \eta^2 M\alpha_\mu^2\,\mathrm{tr}[S^\mu]\cdot E_{\text{opt}} \\
&= \sum_{\nu=1}^N \left((1-2\eta\alpha_\mu^2 + 2\eta^2\alpha_\mu^4)\,\mathbb{1}_{\mu\nu} + \eta^2\alpha_\mu^4 M\right)\cdot\langle E^\nu(n-1)\rangle \\
&\quad + \tfrac{1}{8}\eta^2\sigma_{\text{NP}}^2\alpha_\mu^4\cdot\left(M^3T + 6M^2 + 8\frac{M}{T}\right) + \eta^2\alpha_\mu^4 M\cdot E_{\text{opt}}.
\end{aligned} \tag{S175}
$$

Both relations can be written as

$$
\langle E^\mu(n)\rangle = \sum_{\nu=1}^N (A+B)_{\mu\nu}\cdot\langle E^\nu(n-1)\rangle + b_\mu, \tag{S176}
$$

where $A$ is the same for WP and NP but $B$ and $b$ differ,

$$A_{\mu\nu} = (1-2\eta\alpha_\mu^2 + \eta^2\alpha_\mu^4)\,\mathbb{1}_{\mu\nu}, \tag{S177}$$

$$B_{\mu\nu}^{\text{WP}} = \eta^2\alpha_\mu^2\alpha_\nu^2 M + \eta^2\alpha_\mu^4\delta_{\mu\nu}, \tag{S178}$$

$$B_{\mu\nu}^{\text{NP}} = \eta^2\alpha_\mu^4 M + \eta^2\alpha_\mu^4\delta_{\mu\nu}, \tag{S179}$$

$$b_\mu^{\text{WP}} = \tfrac{1}{8}\eta^2\sigma_{\text{WP}}^2\cdot\left(M^3\alpha_\mu^2\Big(\sum_{\nu=1}^N\alpha_\nu^2\Big)^2 + 2M^2\alpha_\mu^2\sum_{\nu=1}^N\alpha_\nu^4 + 4M^2\alpha_\mu^4\sum_{\nu=1}^N\alpha_\nu^2 + 8M\alpha_\mu^6\right), \tag{S180}$$

$$b_\mu^{\text{NP}} = \tfrac{1}{8}\eta^2\sigma_{\text{NP}}^2\alpha_\mu^4\cdot\left(M^3T + 6M^2 + 8\frac{M}{T}\right) + \eta^2\alpha_\mu^4 M\cdot E_{\text{opt}}. \tag{S181}$$

The matrix $A$ is diagonal, but $B$ is not and mixes the different error components. $A$ originates from the effect of the mean update (cf. Eqs. (S89,S115)), and $B$ from that of credit-assignment-related update fluctuations (Eqs. (S90,S116)). $B_{\mu\nu}$ measures the increase in error component $E^\mu$ due to update fluctuations $\delta w^{\text{cr.as}}$ caused by output perturbations parallel to the $\nu$th input component. The diagonal entries $\eta^2\alpha_\mu^4\delta_{\mu\nu}$ in Eqs. (S178,S179) arise due to correlations between the corresponding update and error signal components. The different form of $B_{\mu\nu}$ for WP and NP can be understood by considering three factors: First, in WP the strength (variance) of the induced output perturbation along the $\nu$th input component is proportional to the strength $\alpha_\nu^2$ of the $\nu$th input component, while for NP output perturbations have the same size along all directions. Second, in NP the update of the weights that read out from the $\mu$th input component are constructed by projecting the output perturbations onto that input component in the eligibility trace (Eq. (6)). The update of weights that read out from the $\mu$th input component thus scales with its strength $\alpha_\mu^2$. WP, on the other hand, constructs updates by multiplying weight perturbations with the same error signal for all weights. Third, the effect of update noise on error component $E_\mu$ scales with the related input strength $\alpha_\mu^2$ for both WP and NP. Together, $B_{\mu\nu}$ thus scales with $\alpha_\mu^2\alpha_\nu^2$ for WP and with $\alpha_\mu^4$ for NP.

Eq. (S176) is solved by

$$
\begin{aligned}
\langle E^\mu(n)\rangle &= \sum_{\nu=1}^N \left(A+B\right)_{\mu\nu}^n\langle E^\nu(0)\rangle + \sum_{\nu=1}^N\sum_{s=0}^{n-1}\left(A+B\right)_{\mu\nu}^s b_\nu \\
&= \sum_{\nu=1}^N \left(A+B\right)_{\mu\nu}^n\langle E^\nu(0)\rangle + \sum_{\nu=1}^N\left(\left(\mathbb{1}-(A+B)\right)^{-1}\left(\mathbb{1}-(A+B)^n\right)\right)_{\mu\nu} b_\nu.
\end{aligned} \tag{S182}
$$

For WP, $A+B$ is symmetric (Eq. (S178)). We can therefore diagonalize it and express the error evolution in terms of error modes that decay independently of each other. This is not possible for NP where $A+B$ is in general non-normal (Eq. (S179)). Therefore, even for the considered convex error function, the error of NP can initially increase due to transient non-normal amplification. The different properties of $A+B$ suggest to perform a diagonalization of $A+B$ for WP and a Schur decomposition for NP to analyze the error evolution of specific networks. We will, however, continue to work in the space of error components and not error modes (the eigen- or Schur vectors of $A+B$) because of the simpler expressions and mechanistic interpretations.

For a fair comparison we set $\sigma^2_{\text{NP}} = \sigma^2_{\text{eff}}$ and $\sigma^2_{\text{WP}} = \sigma^2_{\text{eff}} / \sum_{\nu=1}^{N} \alpha^2_\nu$ (SM1, Sec. "Effective perturbation strength") and express the recursion in terms of the effective perturbation strength. This affects only $b$,

$$b^{\text{WP}}_\mu = \tfrac{1}{8}\eta^2\sigma^2_{\text{eff}} \cdot \big(M^3\alpha^2_\mu \sum_{\nu=1}^{N} \alpha^2_\nu + 2M^2\alpha^2_\mu \frac{\sum_{\nu=1}^{N}\alpha^4_\nu}{\sum_{\nu=1}^{N}\alpha^2_\nu} + 4M^2\alpha^4_\mu + 8M\frac{\alpha^6_\mu}{\sum_{\nu=1}^{N}\alpha^2_\nu}\big), \tag{S183}$$

$$b^{\text{NP}}_\mu = \tfrac{1}{8}\eta^2\sigma^2_{\text{eff}}\alpha^4_\mu \cdot \big(M^3T + 6M^2 + 8\frac{M}{T}\big) + \eta^2\alpha^4_\mu M \cdot E_{\text{opt}}. \tag{S184}$$

#### Evolution of error components related to strong and weak inputs

For infinitesimal perturbation strength and realizable targets, the amount of interference between error components determines the differences in the convergence behavior of the learning rules (cf. the identical diagonal elements of $A + B$ Eq. (S177-S179)). Whether WP or NP generates more interference to an error component $E_\mu$ depends on the concrete distribution of error components in the considered learning step. This distribution, in turn, depends on the initial conditions of the training.

In order to analyze the effect of interference on the convergence of error components related to strong and weak input strengths, we consider a single update. We assume that $\sigma_{\text{eff}}$ is negligible and that the target is realizable, $d = 0$ and $E_{\text{opt}} = 0$. This implies $b = 0$ (Eqs. (S183,S184)) and the error decay is determined by $A + B$ (Eq. (S176)). Inserting the expressions for $B^{\text{WP}|\text{NP}}$, Eqs. (S178,S179), into Eq. (S176) yields

$$\langle E^\mu(n)\rangle \overset{\text{WP}}{=} (A_{\mu\mu} + \eta^2\alpha^4_\mu)\cdot\langle E^\mu(n-1)\rangle + \eta^2\alpha^2_\mu M \cdot \sum_{\nu=1}^{N}\alpha^2_\nu\,\langle E^\nu(n-1)\rangle \tag{S185}$$

$$\langle E^\mu(n)\rangle \overset{\text{NP}}{=} (A_{\mu\mu} + \eta^2\alpha^4_\mu)\cdot\langle E^\mu(n-1)\rangle + \underbrace{\eta^2\alpha^2_\mu M \cdot \alpha^2_\mu \sum_{\nu=1}^{N}\langle E^\nu(n-1)\rangle}_{\equiv \Delta E^{\mu,\text{interference}}_{\text{upd}}}. \tag{S186}$$

The last terms describe the error increase of component $E^\mu$ due to interference with all other components $E^\nu$. Here we compare how the error components of WP and NP evolve after a single update when starting from the same distribution. Consider the ratio of the above interference terms,

$$\frac{\Delta E^{\mu,\text{interference}}_{\text{NP,upd}}}{\Delta E^{\mu,\text{interference}}_{\text{WP,upd}}} = \frac{\alpha^2_\mu \sum_{\nu=1}^{N}\langle E^\nu(n-1)\rangle}{\sum_{\nu=1}^{N}\alpha^2_\nu\langle E^\nu(n-1)\rangle} = \frac{\alpha^2_\mu}{\alpha^2_c}\begin{cases} > 1: & E^\mu \text{ receives less interference for WP,} \\ < 1: & E^\mu \text{ receives less interference for NP,}\end{cases} \tag{S187}$$

where we defined the critical input strength

$$\alpha^2_c = \frac{\sum_{\nu=1}^{N}\alpha^2_\nu\langle E^\nu(n-1)\rangle}{\sum_{\nu=1}^{N}\langle E^\nu(n-1)\rangle}. \tag{S188}$$

Eq. (S187) implies that components connected to strong inputs with $\alpha^2_\mu > \alpha^2_c$ are learned faster for WP while components related to weak inputs with $\alpha^2_\mu < \alpha^2_c$ improve faster for NP. In particular, the largest input component always improves faster or equally fast for WP compared to NP, while the reverse holds for the the smallest input component (in the absence of reward noise).

In networks that are highly noisy (like biological ones) the task output needs to be generated by sufficiently strong latent input components. In other words, there needs to be an input representation that fits the task and clearly exceeds the noise. For initially homogeneously distributed weights this means that the weights related to the strongest input components typically produce the largest errors. Their faster convergence for WP indicates that WP may typically improve faster than NP, at least in the beginning of learning. Towards the end of learning, for fine-tuning, also modifications of weights reading from weaker inputs may be important such that NP becomes faster (Fig. S7).

#### Final weight spread

By projecting the weight matrix onto an input component $\tilde{r}_{\mu t}$ (SM3, Sec. "Task relevant and irrelevant weights"), we can define related weights $W^\mu$ that read out from it and whose mismatch determines the corresponding error component $E^\mu$. Eq. (S173) then becomes

$$E^\mu = \tfrac{1}{2}\,\text{tr}[W\tilde{S}^\mu W^T] = \tfrac{1}{2}\,\text{tr}[W^\mu\tilde{S}^\mu(W^\mu)^T] = \tfrac{1}{2}\alpha^2_\mu|W^\mu|^2. \tag{S189}$$

The squared norm of the weight mismatch, $|W^\mu|^2$, is thus proportional to the related error component (Eq. (S176)) divided by the input strength $\alpha_\mu^2$. If the expectation value $W_{ij}^\mu$ is zero, as in all experiments in main text, Sec. "Theoretical analysis", except when there is weight decay or input noise (which bias $w_{ij}^\mu$ towards zero), then $\langle |W^\mu|^2 \rangle$ is the variance of the weights $w^\mu$.

We now estimate from Eqs. (S176–S181) how the final squared weight mismatch depends on the related input strength. When assuming large $M$ and $T$ and neglecting the self-interaction terms in $A$ and $B$, we can approximate $A$, $B$ and $b$ by their leading order terms

$$A_{\mu\nu} \approx (1 - 2\eta\alpha_\mu^2 + \cancel{\eta^2\alpha_\mu^4})\, \mathbb{1}_{\mu\nu}, \tag{S190}$$

$$B^{\mathrm{WP}}_{\mu\nu} \approx \eta^2\alpha_\mu^2\alpha_\nu^2 M + \cancel{\eta^2\alpha_\mu^4\delta_{\mu\nu}}, \tag{S191}$$

$$B^{\mathrm{NP}}_{\mu\nu} \approx \eta^2\alpha_\mu^4 M + \cancel{\eta^2\alpha_\mu^4\delta_{\mu\nu}}, \tag{S192}$$

$$b^{\mathrm{WP}}_\mu \approx \tfrac{1}{8}\eta^2\sigma_{\mathrm{WP}}^2 \cdot \Big( M^3\alpha_\mu^2 \Big( \sum_{\nu=1}^N \alpha_\nu^2 \Big)^2 + \cancel{2M^2\alpha_\mu^2 \sum_{\nu=1}^N \alpha_\nu^4 + 4M^2\alpha_\mu^4 \sum_{\nu=1}^N \alpha_\nu^2 + 8M\alpha_\mu^6} \Big), \tag{S193}$$

$$b^{\mathrm{NP}}_\mu \approx \tfrac{1}{8}\eta^2\sigma_{\mathrm{NP}}^2\alpha_\mu^4 \cdot \Big( M^3 T + \cancel{6M^2 + 8\frac{M}{T}} \Big) + \eta^2\alpha_\mu^4 M \cdot E_{\mathrm{opt}}. \tag{S194}$$

With these approximations, the expected evolution of the error components (Eq. (S176)) shows a simple dependence on the input strength $\alpha_\mu^2$, as the dependence on $\nu$ factors out. For WP we find

$$\begin{aligned} \langle E^\mu(n) \rangle &\overset{\mathrm{WP}}{\approx} (1 - 2\eta\alpha_\mu^2)\langle E^\mu(n-1)\rangle + \eta^2\alpha_\mu^2 M \sum_{\nu=1}^N \alpha_\nu^2 \langle E^\nu(n-1)\rangle + (\tfrac{1}{8}\eta^2\sigma_{\mathrm{WP}}^2 M^3 \alpha_\mu^2) \cdot \Big( \sum_{\nu=1}^N \alpha_\nu^2 \Big)^2 \\ &= \langle E^\mu(n-1)\rangle + \alpha_\mu^2 \cdot \underbrace{\Bigg( -2\eta\langle E^\mu(n-1)\rangle + \overbrace{\eta^2 M \sum_{\nu=1}^N \alpha_\nu^2 \langle E^\nu(n-1)\rangle + \tfrac{1}{8}\eta^2\sigma_{\mathrm{WP}}^2 M^3 \cdot \Big( \sum_{\nu=1}^N \alpha_\nu^2 \Big)^2}^{\text{independent of } \mu} \Bigg)}_{=\,0 \text{ after convergence if } \alpha_\mu \neq 0}. \end{aligned} \tag{S195}$$

After convergence $\langle E^\mu(n) \rangle = \langle E^\mu(n-1) \rangle$ such that either $\alpha_\mu = 0$ or the highlighted term inside the brackets must be zero. For relevant weights we can thus equate the bracket to zero and solve for the first $\langle E^\mu(n-1) \rangle$ inside it. This reveals that $\langle E^\mu(n \to \infty) \rangle$ is independent of $\mu$ and together with Eq. (S189) the scaling of $\langle |W^\mu(n \to \infty)|^2 \rangle$ with $\alpha_\mu$,

$$\langle E^\mu(n \to \infty) \rangle \text{ is independent of } \alpha_\mu^2, \qquad \langle |W^\mu(n \to \infty)|^2 \rangle \propto \frac{1}{\alpha_\mu^2}. \qquad \text{(WP)} \tag{S196}$$

We conclude that for WP, each error component $E^\mu$ adds the same contribution to the final error, regardless of the strength $\alpha_\mu^2$ of its corresponding input component, unless it is exactly zero. Eq. (S196) further states that for nonzero input components the squared norm $\langle |W^\mu|^2 \rangle$ after convergence, or the weights' variance, scales inversely with the input strengths $\alpha_\mu^2$. In particular, weights associated with weak inputs will diffuse strongly but settle with a large, finite variance. Only input components that are exactly zero contribute $E^\mu = 0$ to the error. Their weights are completely irrelevant and diffuse to an infinitely broad distribution. In the main text, Sec. "Input noise", and in the next Sec. "Small input components" we argue that small but nonzero input components can still be practically irrelevant if the length of the training and/or the weight strength are limited.

For NP, Eq. (S176) simplifies to

$$\begin{aligned} \langle E^\mu(n) \rangle &\overset{\mathrm{NP}}{\approx} (1 - 2\eta\alpha_\mu^2)\langle E^\mu(n-1)\rangle + \eta^2\alpha_\mu^4 M \langle E(n-1)\rangle + \tfrac{1}{8}\eta^2\sigma_{\mathrm{NP}}^2\alpha_\mu^4 M^3 T + \eta^2\alpha_\mu^4 M \cdot E_{\mathrm{opt}} \\ &= \langle E^\mu(n-1)\rangle \underbrace{-\alpha_\mu^2 \cdot 2\eta\langle E^\mu(n-1)\rangle + \alpha_\mu^4 \cdot \Big( \overbrace{\eta^2 M \langle E(n-1)\rangle + \tfrac{1}{8}\eta^2\sigma_{\mathrm{NP}}^2 M^3 T + \eta^2 M \cdot E_{\mathrm{opt}}}^{\text{independent of } \mu} \Big)}_{=\,0 \text{ after convergence}} \end{aligned} \tag{S197}$$

Again the highlighted terms cancel after convergence of the error and can for nonzero input strength $\alpha_\mu^2$ be solved for $\langle E^\mu(n \to \infty)$. This yields

$$\langle E^\mu(n \to \infty) \rangle \propto \alpha_\mu^2, \qquad \langle |W^\mu(n \to \infty)|^2 \rangle \text{ is independent of } \alpha_\mu^2. \qquad \text{(NP)} \tag{S198}$$

We conclude that, in contrast to WP, error components related to weak input components contribute little to the final error, while those related to strong input components contribute most. Eq. (S198) further shows that for NP the final variance of the weight distribution is independent of $\alpha_\mu^2$; each weight is learned with the same precision.

#### Small input components

Strictly speaking, weights are completely irrelevant only if the inputs that they read out from are exactly zero (assuming without loss of generality rotated inputs). This is because changing the weights of any nonzero input has some effect on the output. Similarly, target components are completely unrealizable only if they are not present at all in the input. This raises the question whether our findings generalize to small but nonzero inputs.

To address it, we note that weights that read out from small inputs need to be large to have a sizeable effect on output and error. Indeed, Eq. (S189) shows that the size of the weight mismatch necessary to cause an error contribution $E^\mu$ is inversely proportional to the input strength, $|W^\mu|^2 \sim \alpha_\mu^{-2}$. For a given error tolerance, this means that the contribution from weights that read out weak inputs can be neglected as long as these *effectively irrelevant* weights remain small enough.

One setting in which these weights remain small arises if the training duration is long enough for the relevant weights to converge, but too short for the effectively irrelevant weights to noticeably contribute to the error. Fig. 6c shows such a scenario, in which the weak inputs are given by white input noise.

Weight decay can also contain the growth of effectively irrelevant weights (Fig. 2bii,S1b, SM4, Sec. "Networks with weight decay"). We expect this to work particularly well if there is a separation of timescales: if the convergence time of relevant weights is shorter than the time after which the effectively irrelevant weights make a sizeable contribution to the error, then the weight decay can operate on the longer timescale and only weakly affect the relevant weights (Fig. 6c). Introducing an upper bound for the magnitude of individual weights can similarly limit the error contribution from effectively irrelevant weights. Depending on the task setting, there may be a bound that is both large enough to not limit the relevant weights and small enough so that effectively irrelevant weights can be neglected.

## SM 7

### Input and perturbation correlations

##### Invariance of GD, WP and NP to temporal correlations and task reordering

The learning of GD, WP and NP is not affected by correlations in the inputs. More precisely: the probability distribution of learning curves does not depend on the temporal correlations of the stochastic process that the input vectors and the additional, unrealizable target components are drawn from (Fig. S3), but only on its one-dimensional finite-dimensional distributions. GD and WP's independence of temporal input and target correlations as well as of reordering or permuting the task originates from the fact that they depend only on the current weight mismatch $W$, the matrix $S$ of instantaneous input correlations and (WP only) on the applied noise $\xi^{\mathrm{WP}}$ (Eqs. (S14,S24,S29,S30)). $W$ and $\xi^{\mathrm{WP}}$ do not depend on the input and $S_{jk}$ is not affected by relations between $r_{jt}$ and $r_{ks}$ with $s \neq t$. NP additionally depends on the projections of $\xi^{\mathrm{NP}}$ on the input $r$ and the unrealizable target component $d$ (Eqs. (S14,S41,42)). The probability distributions of these projections are not affected by temporal correlations in $r_{jt}$ and $d_{it}$, because the iid entries of the $i$th perturbation vector $\xi^{\mathrm{NP}}_{it}$ jointly have a $T$-dimensional, rotationally symmetric Gaussian distribution. Its projection on a constant vector is thus independent of the direction of the vector. In more detail:

- The error of GD, Eq. (8), depends on $W$ and $S$. As as consequence, the GD weight update depends on $W$ and $S$.
- The error of WP, Eq. (8), depends on $W$ and $S$. The error of WP after the perturbation, Eq. (S29), depends on $W$, $S$ and $\xi^{\mathrm{WP}}$. As a consequence, the WP weight update, Eq. (S30) depends on $W$, $S$ and $\xi^{\mathrm{WP}}$.
- The error of NP, Eq. (8), depends on $W$ and $S$. The error of NP after the perturbation, Eq. (S41), depends on W and the projections $\sum_t r_{jt}\xi^{\mathrm{NP}}_{jt}$ and $\sum_t d_{jt}\xi^{\mathrm{NP}}_{jt}$. The eligibility trace is $\sum_t r_{jt}\xi^{\mathrm{NP}}_{jt}$. The NP weight update Eq. (S42) therefore depends on $W$, $\xi^{\mathrm{NP}}$ and the projections $\sum_t r_{jt}\xi^{\mathrm{NP}}_{jt}$ and $\sum_t d_{jt}\xi^{\mathrm{NP}}_{jt}$.

Reordering and correlations within the input activity at different times do not affect $S$ (and $W$ and $\xi^{\mathrm{WP}}$). Therefore they do not affect the learning process of GD and WP. Further, reordering and correlations do not affect the probability distributions of the projections of $\xi^{\mathrm{NP}}$ on $r$ and $d$. The former holds because, the $\xi^{\mathrm{NP}}_{it}$ are are at different time points identically distributed. Temporal correlations in $r_{jt}$ and $d_{it}$ leave the distributions invariant, because the iid entries of the $i$th perturbation vector $\xi^{\mathrm{NP}}_{it}$ jointly have a $T$-dimensional, rotationally symmetric Gaussian distribution. Its projection on a constant vector is thus independent of the direction of the vector. Fig. S3 illustrates the invariance of the learning curves with respect to the introduction of input correlations by numerical simulations.

##### NPc: Learning with temporally correlated node perturbations

This section introduces and discusses in more detail NPc, which learns time-correlated tasks with time-correlated node perturbations. To obtain the correlated node perturbations we draw the initial perturbation at $t = 0$ and create later perturbations according to

$$\xi^{\mathrm{NPc}}_{i0} = \sigma_{\mathrm{eff}} \cdot \tilde{\xi}_{i0}, \qquad \xi^{\mathrm{NPc}}_{it+1} = \gamma \xi^{\mathrm{NPc}}_{it} + \sqrt{1-\gamma^2}\sigma_{\mathrm{eff}} \cdot \tilde{\xi}_{it+1}. \tag{S199}$$

Here $\tilde{\xi}_{it}$ is Gaussian white noise,

$$\tilde{\xi}_{it} \sim \mathcal{N}(0,1), \qquad \langle \xi^{\mathrm{NPc}}_{it} \xi^{\mathrm{NPc}}_{mt+s} \rangle = \delta_{im}\gamma^{|s|}\sigma^2_{\mathrm{eff}}, \tag{S200}$$

and the factor $\gamma = \exp(-1/\tau_{\mathrm{NPc}})$ determines the correlation time $\tau_{\mathrm{NPc}}$. This can be linked to an effective time dimension $T_{\mathrm{eff}}$,

$$T^{\mathrm{pert}}_{\mathrm{eff}} = \frac{T}{\tau_{\mathrm{NPc}}+1}, \qquad \gamma = \exp\left(-\frac{T^{\mathrm{pert}}_{\mathrm{eff}}}{T - T^{\mathrm{pert}}_{\mathrm{eff}}}\right). \tag{S201}$$

For $T^{\mathrm{pert}}_{\mathrm{eff}} = T$, we define $\gamma = \tau_{\mathrm{NPc}} = 0$; in this case NPc is exactly identical to NP. A low effective (temporal) perturbation dimension $T^{\mathrm{pert}}_{\mathrm{eff}}$ means that the perturbations' variance concentrates on only few components.

## SM 8

### Improved learning rules

The new insights into the mechanisms of WP and NP gained through this work enable the construction of related, under certain conditions more powerful learning rules. We note that also NPc (cf. main text, section "Input and perturbation correlations" and SM7) may be considered as such an improved method. In the current part we describe in detail the modified WP rule WP0 and hybrid perturbation (HP). WP0 is useful if the input is sparse, HP is useful if the inputs have the same strength.

##### WP0: Assign zero credit to zero inputs

The WP scheme has no direct way of solving the credit assignment problem of finding the weight perturbations that were responsible for causing the error signal $\Delta E^{\mathrm{lin}}_{\mathrm{pert}}$. Therefore it updates all weights, with the consequence that irrelevant weights diffuse and convergence slows down when multiple input patterns have to be learned.

There is a straightforward way to improve credit assignment for the case that some inputs are zero (or negligible), because then the corresponding weights do not affect the output and error - they cannot be credited for any reward changes. This information, i.e. the absence of presynaptic input, is locally available to synapses and its use means incorporation of model information, promising to improve the learning rule.

Using these observations, the update equation of the improved learning rule WP0 reads

$$\Delta w_{ij}^{\mathrm{WP0}} = \begin{cases} 0 & \text{if } r_{jt} = 0\ \forall t, \\ -\frac{\eta}{\sigma_{\mathrm{WP}}^2}\left(E^{\mathrm{pert}} - E\right)\xi_{ij}^{\mathrm{WP}} & \text{else,} \end{cases} \tag{S202}$$

where the weight perturbations of WP0 are drawn from the same distribution as the $\xi_{ij}^{\mathrm{WP}}$ of WP. We note that output nonlinearities with plateaus, i.e. $g'(y) = 0$ for total weighted inputs $y$ in some range, allow to further improve WP0 in alike manner: the weight $w_{ij}$ should not be changed if $g'(y_{it})r_{jt} = 0\ \forall t$, since the tried small perturbation $\xi_{ij}^{\mathrm{WP}}$ of $w_{ij}$ cannot have influenced the output and the error. Neurons with such nonlinearities are for example rate neurons with ReLU activation functions or spiking neurons with a spike threshold.

In practice, inputs and $g'$ may not be exactly zero. One can then introduce a threshold below which the weights are not updated. The largest improvements of WP0 over WP are expected if coding is sparse (many inputs are zero) and only a subset of postsynaptic neurons is non-saturated (has $g' \neq 0$).

##### Hybrid perturbation (HP): Using WP to produce output perturbations and NP to produce updates

WP and NP have different advantages: the output perturbations of WP lie completely in the realizable subspace and do not interfere with unrealizable target components, while NP's use of eligibility traces lets it solve part of the credit assignment problem such that irrelevant weights do not diffuse and NP converges faster than WP when multiple input patterns have to be learned. We here aim to combine the advantageous features of both learning rules into one rule, HP.

To this end, output perturbations are induced by perturbing the weights like in WP,

$$z_{it}^{\mathrm{pert,HP}} = \sum_{j=1}^{N}(w_{ij} + \xi_{ij}^{\mathrm{WP}})r_{jt}, \tag{S203}$$

where $g(\cdot) = \mathrm{Id}(\cdot)$ and we named the weight perturbations of HP $\xi_{ij}^{\mathrm{WP}}$, as they are drawn from the same distribution as for WP. Thus they do not interfere with unrealizable target components, which only shift the final error of HP by $E_{\mathrm{opt}}$ as for WP (Eq. (S124) and Fig. 4). They induce output perturbations of the form $\xi_{it}^{\mathrm{out}} = \sum_{j=1}^{N}\xi_{ij}^{\mathrm{WP}}r_{jt}$, which are used to calculate NP-like updates by using eligibility traces,

$$\begin{aligned}\Delta w_{ij}^{\mathrm{HP}} &= -\frac{\eta}{\sigma_{\mathrm{WP}}^2 T}(E^{\mathrm{pert}} - E)\sum_{t=1}^{T}\xi_{it}^{\mathrm{out}}r_{jt} \\ &= -\frac{\eta}{\sigma_{\mathrm{WP}}^2 T}(E^{\mathrm{pert}} - E)\sum_{t=1}^{T}\sum_{k=1}^{N}\xi_{ik}^{\mathrm{WP}}r_{kt}r_{jt} \\ &= -\frac{\eta}{\sigma_{\mathrm{WP}}^2}(E^{\mathrm{pert}} - E)\sum_{k=1}^{N}\xi_{ik}^{\mathrm{WP}}S_{kj}.\end{aligned} \tag{S204}$$

The normalization $1/(\sigma_{\mathrm{WP}}^2 T)$ contains an additional factor $1/T$. The perturbed error is the same as for WP,

$$E^{\mathrm{pert}} = \tfrac{1}{2}\,\mathrm{tr}[(W+\xi^{\mathrm{WP}})S(W+\xi^{\mathrm{WP}})^T] + E_{\mathrm{opt}} = E + \mathrm{tr}[WS\xi^{\mathrm{WP}^T}] + \tfrac{1}{2}\,\mathrm{tr}[\xi^{\mathrm{WP}}S\xi^{\mathrm{WP}^T}], \tag{S205}$$

such that the update reads

$$\Delta w_{ij} = -\frac{\eta}{\sigma_{\mathrm{WP}}^2}\left(\mathrm{tr}[WS\xi^{\mathrm{WP}^T}] + \tfrac{1}{2}\,\mathrm{tr}[\xi^{\mathrm{WP}}S\xi^{\mathrm{WP}^T}]\right)\sum_{k=1}^{N}\xi_{ik}^{\mathrm{WP}}S_{kj}. \tag{S206}$$

For brevity, we now only consider the linear components of the error signal (i.e. $\mathrm{tr}[WS\xi^{\mathrm{WP}^T}]$) that determine the convergence speed and behavior, neglecting the reward noise $\Delta E_{\mathrm{pert}}^{\mathrm{quad}} = \frac{1}{2}\,\mathrm{tr}[\xi^{\mathrm{WP}}S\xi^{\mathrm{WP}^T}]$. (This is equivalent to learning in the small $\sigma_{\mathrm{eff}}$ limit; realizability of targets does not affect HP anyways.) Then the $\mu$th component of the expected error after an update is

$$\langle E^\mu(n+1)\rangle = \langle E^\mu(n)\rangle + \langle\mathrm{tr}[WS^\mu\Delta w^T]\rangle + \tfrac{1}{2}\langle\mathrm{tr}[\Delta w S^\mu \Delta w^T]\rangle \tag{S207}$$

(Eq. (S173)), where the effect of the mean update is

$$\begin{aligned}
\langle\mathrm{tr}[WS^\mu\Delta w^T]\rangle &= -\frac{\eta}{\sigma_{\mathrm{WP}}^2}\langle\mathrm{tr}[WS^\mu S\xi^{\mathrm{WP}^T}]\,\mathrm{tr}[WS\xi^{\mathrm{WP}^T}]\rangle \\
&= -\frac{\eta}{\sigma_{\mathrm{WP}}^2}\sum_{im=1}^{M}\sum_{jklpq=1}^{N} W_{ij}S_{jk}^\mu S_{kl}W_{mp}S_{pq}\langle\xi_{il}^{\mathrm{WP}}\xi_{mq}^{\mathrm{WP}}\rangle \\
&= -\eta\,\mathrm{tr}[WS^\mu S^2 W^T] = -2\eta\alpha_\mu^4\cdot\langle E^\mu(n)\rangle.
\end{aligned} \tag{S208}$$

The quadratic contributions, which describe update fluctuations due to the credit assignment problem and which are responsible for slowing down the learning, are

$$\begin{aligned}
\tfrac{1}{2}\langle\mathrm{tr}[\Delta w S^\mu\Delta w^T]\rangle &= \frac{\eta^2}{\sigma_{\mathrm{WP}}^4}\tfrac{1}{2}\langle\mathrm{tr}[WS\xi^{\mathrm{WP}^T}]\,\mathrm{tr}[\xi^{\mathrm{WP}}SS^\mu S\xi^{\mathrm{WP}^T}]\,\mathrm{tr}[WS\xi^{\mathrm{WP}^T}]\rangle \\
&= \frac{\eta^2}{2\sigma_{\mathrm{WP}}^4}\sum_{imn=1}^{M}\sum_{jkpqrstu=1}^{N}\langle W_{ij}S_{jk}\xi_{ik}^{\mathrm{WP}}\xi_{mp}^{\mathrm{WP}}S_{pq}S_{qr}^\mu S_{rs}\xi_{ms}^{\mathrm{WP}}W_{nt}S_{tu}\xi_{nu}^{\mathrm{WP}}\rangle \\
&= \frac{\eta^2}{2\sigma_{\mathrm{WP}}^4}\sum_{imn=1}^{M}\sum_{jkpqrstu=1}^{N} W_{ij}S_{jk}S_{pq}S_{qr}^\mu S_{rs}W_{nt}S_{tu}\langle\xi_{ik}^{\mathrm{WP}}\xi_{mp}^{\mathrm{WP}}\xi_{ms}^{\mathrm{WP}}\xi_{nu}^{\mathrm{WP}}\rangle.
\end{aligned} \tag{S209}$$

Using $\langle\xi_{ij}^{\mathrm{WP}}\xi_{mk}^{\mathrm{WP}}\rangle = \sigma_{\mathrm{WP}}^2\delta_{im}\delta_{jk}$ and Isserli's theorem, the appearing moment of $\xi^{\mathrm{WP}}$ evaluates to

$$\langle\xi_{ik}^{\mathrm{WP}}\xi_{mp}^{\mathrm{WP}}\xi_{ms}^{\mathrm{WP}}\xi_{nu}^{\mathrm{WP}}\rangle = \sigma_{\mathrm{WP}}^4\left(\delta_{in}\delta_{ku}\delta_{ps} + \delta_{im}\delta_{mn}(\delta_{kp}\delta_{su} + \delta_{ks}\delta_{pu}\right) \tag{S210}$$

such that

$$\begin{aligned}
\tfrac{1}{2}\langle\mathrm{tr}[\Delta w S^\mu\Delta w^T]\rangle &= \tfrac{1}{2}\eta^2 M\,\mathrm{tr}[WSSW^T]\cdot\mathrm{tr}[SS^\mu S] + \eta^2\,\mathrm{tr}[WSSS^\mu SSW^T] \\
&= \eta^2 M\alpha_\mu^6\cdot\sum_{\nu=1}^{N}\alpha_\nu^2\langle E^\nu(n)\rangle + 2\eta^2\alpha_\mu^8\langle E^\mu(n)\rangle.
\end{aligned} \tag{S211}$$

With this, the evolution of the expected error is described by

$$\langle E^\mu(n+1)\rangle = \sum_{\nu=1}^{N}(A+B)_{\mu\nu}\langle E^\nu(n)\rangle, \tag{S212}$$

where

$$A_{\mu\nu} = (1 - 2\eta\alpha_\mu^4 + \eta^2\alpha_\mu^8)\delta_{\mu\nu}, \tag{S213}$$

$$B_{\mu\nu} = \eta^2 M\alpha_\mu^2\alpha_\nu^6 + \eta^2\alpha_\mu^8\delta_{\mu\nu}. \tag{S214}$$

For the case of repeating inputs and where $N_{\mathrm{eff}}$ eigenvalues of $S$ are equal to $\alpha^2$ and all others zero, the expected error evolves as

$$\langle E(n+1)\rangle = \left(\langle E(n)\rangle - E_{\mathrm{opt}}\right)\cdot a + E_{\mathrm{opt}}, \qquad a = 1 - 2\eta\alpha^4 + \eta^2\alpha^8(MN_{\mathrm{eff}}+2). \tag{S215}$$

Minimizing $a$ leads to an optimal learning rate and convergence factor of

$$\eta^* = \frac{1}{(MN_{\text{eff}} + 2)\alpha^4}, \qquad a^* = 1 - \frac{1}{MN_{\text{eff}} + 2}. \tag{S216}$$

In this case, HP converges as fast as both WP and NP. In addition, irrelevant weights do not diverge, which benefits the learning of multiple subtasks. Further we expect the final error to be lower than for NP because all output perturbations lie in the realizable subspace, which is confirmed by our numerical simulations, Fig. 4. Thus for latent inputs that have the same strength, HP combines the advantages of both WP and NP. Note that, as for WP and NP but in contrast to WP0, we can without loss of generality change to a set of rotated inputs, because weight perturbations are isotropic such that the update equation Eq. (S204) is invariant under the rotation. This is also reflected by Eqs. (S212–S214): the equations show that the error evolution only depends on the eigenvalues $\alpha_\mu^2$ of $S$, which are invariant to input rotations.

We observed that application of HP to the reservoir computing task gave worse performance than application of WP and NP (main text, Sec. "Conclusions from the theoretical analysis and new learning rules"). We explain this by the fact that HP generates biased updates for latent inputs with different strengths (a similar explanation may hold for the unsatisfactory performance on MNIST): weights connected to stronger input components both have a larger impact on output perturbations, as for WP, and get updated more strongly due to their eligibility traces being larger, as for NP (compare Eqs. (S178,S179)). The mean update is thus biased,

$$\begin{aligned}\langle \Delta w_{ij}^{\text{HP}} \rangle &= -\frac{\eta}{\sigma_{\text{WP}}^2} \left\langle \text{tr}[WS\xi^{\text{WP}T}] \sum_{k=1}^{N} \xi_{ik}^{\text{WP}} S_{kj} \right\rangle = -\frac{\eta}{\sigma_{\text{WP}}^2} \sum_{im=1}^{M} \sum_{klp=1}^{N} W_{ml} S_{lp} S_{kj} \langle \xi_{mp}^{\text{WP}} \xi_{ik}^{\text{WP}} \rangle \\ &= -\eta \sum_{i=1}^{M} \sum_{jk=1}^{N} W_{il} S_{lk} S_{kj} = -\eta \sum_{k=1}^{N} \frac{\partial E}{\partial w_{ik}} S_{kj}. \end{aligned} \tag{S217}$$

If inputs are rotated so that $w_{i\mu}$ reads out from the $\mu$th input component $r_{\mu t}$ of strength $\alpha_\mu^2$, then the mean updates $\langle \Delta w_{i\mu}^{\text{HP}} \rangle = -\eta \frac{\partial E}{\partial w_{i\mu}} \cdot \alpha_\mu^2$ are proportional to their weight error gradients multiplied by $\alpha_\mu^2$. This means that weights related to weak inputs are updated only little and will take a long time to converge. If such weak inputs are important to realize the target, HP will perform worse than NP and WP. Adding a network layer that equalizes the non-zero (or non-negligible) input strengths in each subtask might render HP beneficial and applicable.

Recovering unbiased updates in presence of inhomogeneous input strengths by multiplying updates with the inverse correlation matrix (which is a non-local operation), $\Delta w^{\text{HP}} \to \Delta w^{\text{HP}} S^{-1}$, simply reduces HP to WP,

$$\begin{aligned}\Delta w_{ij}^{\text{HP,unbiased}} &= -\frac{\eta}{\sigma_{\text{WP}}^2 T}(E^{\text{pert}} - E) \sum_{l=1}^{N} \sum_{t=1}^{T} \xi_{it}^{\text{out}} r_{lt} \cdot S_{lj}^{-1} = -\frac{\eta}{\sigma_{\text{WP}}^2 T}(E^{\text{pert}} - E) \sum_{t=1}^{T} \sum_{kl=1}^{N} \xi_{ik}^{\text{WP}} r_{kt} r_{lt} S_{lj}^{-1} \\ &= -\frac{\eta}{\sigma_{\text{WP}}^2}(E^{\text{pert}} - E) \sum_{kl=1}^{N} \xi_{ik}^{\text{WP}} S_{kl} S_{lj}^{-1} = -\frac{\eta}{\sigma_{\text{WP}}^2}(E^{\text{pert}} - E)\xi_{ij}^{\text{WP}} = \Delta w_{ij}^{\text{WP}}, \end{aligned} \tag{S218}$$

as long as all input strengths are non-zero such that $S^{-1}$ is well-defined. Setting the eigenvalues of $S^{-1}$ related to zero inputs to zero (a priori they are undefined) means that irrelevant weight combinations are not updated, which for rotated inputs reduces the unbiased version of HP to WP0.

### Figures and Data

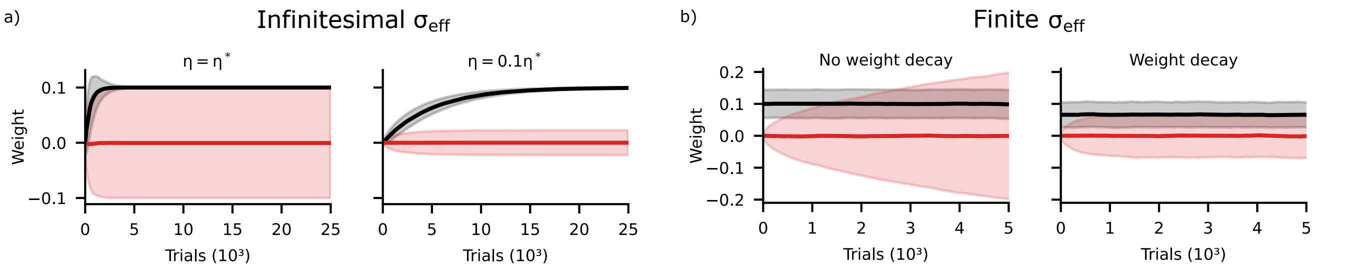


**Figure S1.** Further analysis of the weight diffusion in WP. a) Diffusion of irrelevant weights is transient for infinitesimal perturbation size. Display like main text, Fig. 2b, but for infinitesimal $\sigma_{\mathrm{WP}}$. The relevant weights converge to the relevant weights of the teacher network, $w^*_{\mathrm{rel},i} = 0.1$. Learning with optimal rate ($\eta = \eta^*$) leads to a final standard deviation of irrelevant weights of the same size as the mean relevant weights (left); a smaller learning rate leads to less weight diffusion (right). SM4, Sec. "Transient weight diffusion due to credit assignment-related update fluctuations" explains these observations. b) Weight diffusion for finite perturbation size, after the output error has decayed to its stationary residual value and the relevant weights fluctuate around their targets. The irrelevant weights, which are initially set to zero, diffuse without bounds (left). In particular, the standard deviation of the weight distribution grows like $\sim \sqrt{n}$ with the learning trial number $n$. A tendency of the weights to decay confines this growth (right).

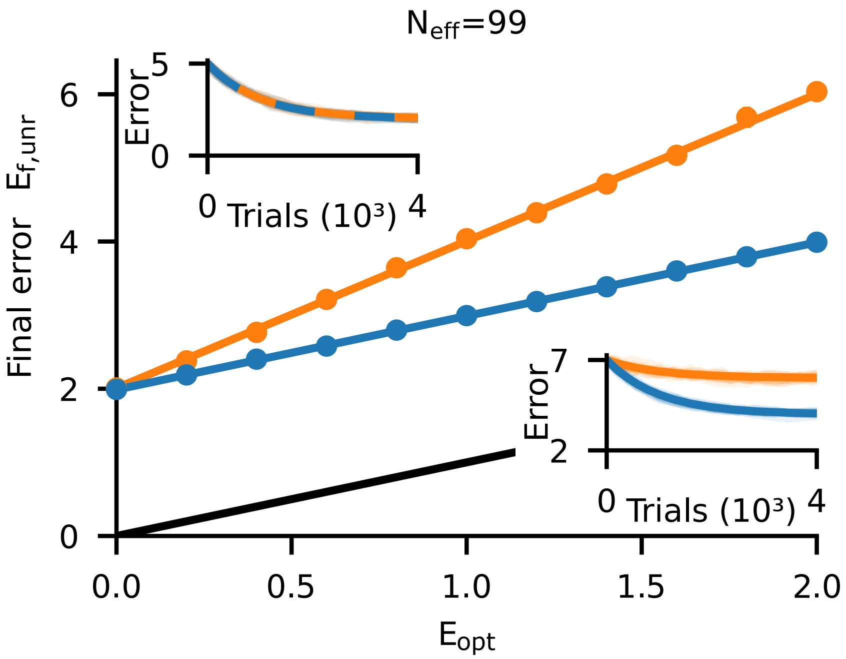


**Figure S2.** Final error after convergence as a function of the limiting error $E_{\mathrm{opt}}$ like Fig. 3b, but for effective input dimensionality $N_{\mathrm{eff}} = 99$ (instead of $N_{\mathrm{eff}} = 50$). Since $T = 100$ and $N_{\mathrm{eff}} = 99$ ($N_{\mathrm{eff}}$ must be smaller than $T$ to allow an unrealizable part), both algorithms perform basically the same for $E_{\mathrm{opt}} = 0$ (cf. also light curves in main text, Fig. 1b, left). For $E_{\mathrm{opt}} > 0$ the final error of WP is shifted by $E_{\mathrm{opt}}$ while that of NP increases approximately by $2E_{\mathrm{opt}}$.

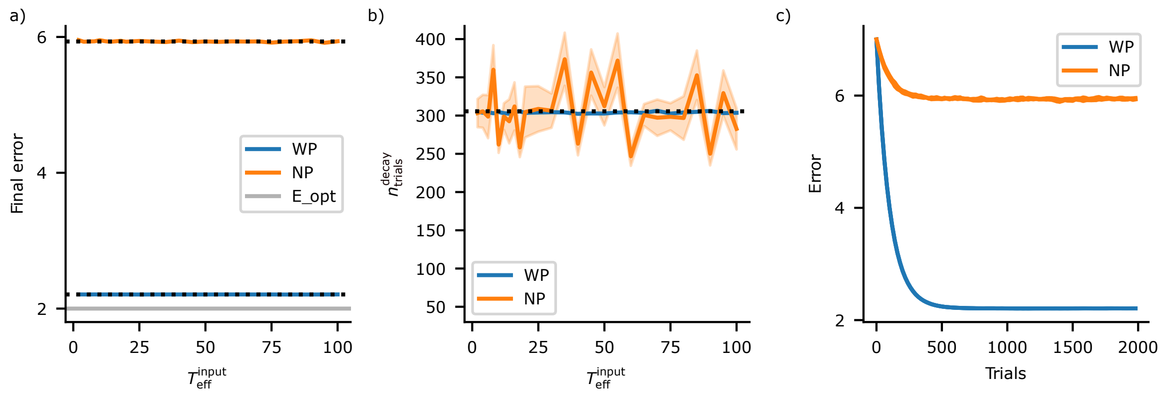


**Figure S3.** WP and NP are unaffected by input correlations. a) Final error of WP (blue) and NP (orange) in a task where the inputs are temporally correlated, versus the effective temporal dimension $T_{\mathrm{eff}}^{\mathrm{input}}$ of the inputs. For $T_{\mathrm{eff}}^{\mathrm{input}} = T = 100$ we have uncorrelated input, for $T_{\mathrm{eff}}^{\mathrm{input}} = 2$ the input traces vary very slowly. Inputs are generated by orthonormalizing exponentially filtered white noise. Realizable target components are linear combinations of the correlated inputs. We assume that there is an additional unrealizable target component, which, for simplicity, contains all modes orthogonal to the inputs with equal strength. Our theoretical results (black dotted curves) predict that the final error is independent of the correlation time for both WP and NP. This is confirmed by the numerical simulations (colored curves, overlayed by the theoretical ones). b) Number of trials to reach $95\%$ of the final error reduction for WP (blue) and NP (orange) as a function of $T_{\mathrm{eff}}^{\mathrm{input}}$. Our theoretical results (black dotted curves, overlapping) predict that the decay time of the expected error is independent of the correlation time and the same for both WP and NP. This is confirmed by the numerical simulations (mean: colored curves, partially underlaying the theoretical ones, shaded: standard error of the mean). Since the learning curves of NP are more variable, its mean convergence time has a higher standard error. c) Mean error (solid) and standard error of the mean error (shaded) as a function of trial number for different input correlation times (effective temporal input dimensions: $T_{\mathrm{eff}}^{\mathrm{input}} = 2, 10, 100$). The curves agree within their errors and are not visually distinguishable. Parameters: $M = N_{\mathrm{eff}} = 10$, $T = 100$, $\sigma_{\mathrm{eff}} = 0.04$, $E_{\mathrm{opt}} = 2$. $T_{\mathrm{eff}}^{\mathrm{input}}$ is in (a,b) varied from $2$ to $100$ in steps of $2$ between $2$ and $20$ and in steps of $5$ between $25$ and $100$, with $1000$ repetitions each. The final error in (a) is measured after $2000$ trials.

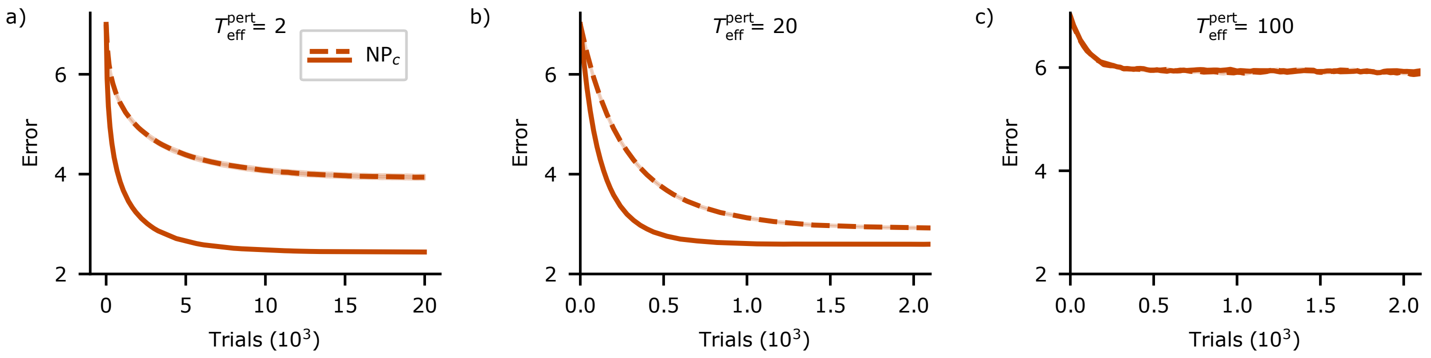


**Figure S4.** Exemplary error decay curves of NPc (mean and SEM) for different input and perturbation correlation times. a) For $T_{\text{eff}}^{\text{pert}} = 2$, that is for larger perturbation than input correlation time, convergence slows down (note the changed x-axis scale compared to b,c). For correlated inputs (solid) but not for uncorrelated inputs (dashed), NPc still achieves a low final error. b) For $T_{\text{eff}}^{\text{pert}} = 20$, the perturbations of NPc have the same effective temporal dimension as the correlated inputs (solid curves). NPc simultaneously achieves a low final error and fast convergence. c) For $T_{\text{eff}}^{\text{pert}} = 100 = T$, NPc reduces to NP and settles at the same high final error. As NP is insensitive to input correlations (Fig. S3), the curves of NPc for correlated and uncorrelated inputs here agree. Parameters: $N_{\text{eff}} = M = 10$, $N = T = 100$, $\sigma_{\text{eff}} = 0.04$, $E_{\text{opt}} = 2$. Latent inputs of equal strength are constructed by orthonormalizing white noise (red dashed) or exponentially filtered white noise (with time constant $\tau_{\text{corr}}^{\text{input}} = 4$ and thus $T_{\text{eff}}^{\text{input}} = 20$, red solid). Unrealizable target components are constructed from the last $T - N_{\text{eff}}$ orthonormalized noise traces with equal strengths.

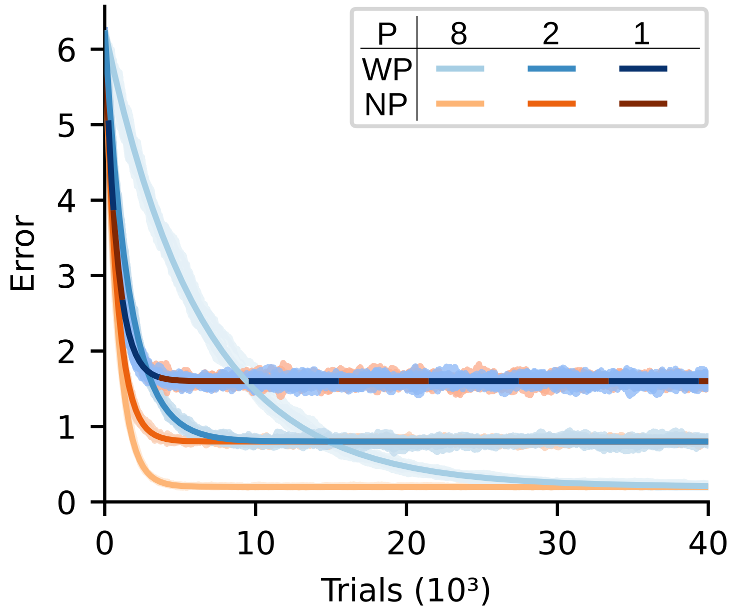


**Figure S5.** Error dynamics scale differently for WP and NP when splitting the input into different patterns for different trials. The network is the same as the one used in Fig. 1 ($N = 100$, $M = 10$, $\sigma_{\mathrm{eff}} = 4 \times 10^{-2}$). The task is to reproduce the output of a teacher network in response to input with a dimensionality of $N_{\mathrm{eff}}^{\mathrm{task}} = 80$, which we split into $P$ non-overlapping input patterns each having dimensionality $N_{\mathrm{eff}}^{\mathrm{trial}} = N_{\mathrm{eff}}^{\mathrm{task}}/P$ (hence $PN_{\mathrm{eff}}^{\mathrm{trial}} = 80$). For simplicity, we use input patterns where at each timestep a different input unit has the value $\sqrt{N/N_{\mathrm{eff}}^{\mathrm{task}}}$, while all other input units are zero. This implies $T = N_{\mathrm{eff}}^{\mathrm{trial}}$. The figure shows error curves for WP (blue) and NP (orange) from simulations (10 runs, shaded) together with analytical curves for the decay of the expected error (solid), for different values of $P$ (simulation results for NP with $P = 8$ are mostly covered by the analytical curve). Theoretical curves and simulations agree well. The convergence speed of WP decreases with increasing $P$ (Eqs. (S148,S162)). The convergence speed of NP is almost unaffected by $P$ for NP (Eqs. (S149,S163)). The residual error is inversely proportional to $P$ for both WP and NP (Eqs. (S152,S153)) and it is equal for WP and NP because $T = N_{\mathrm{eff}}^{\mathrm{trial}}$.

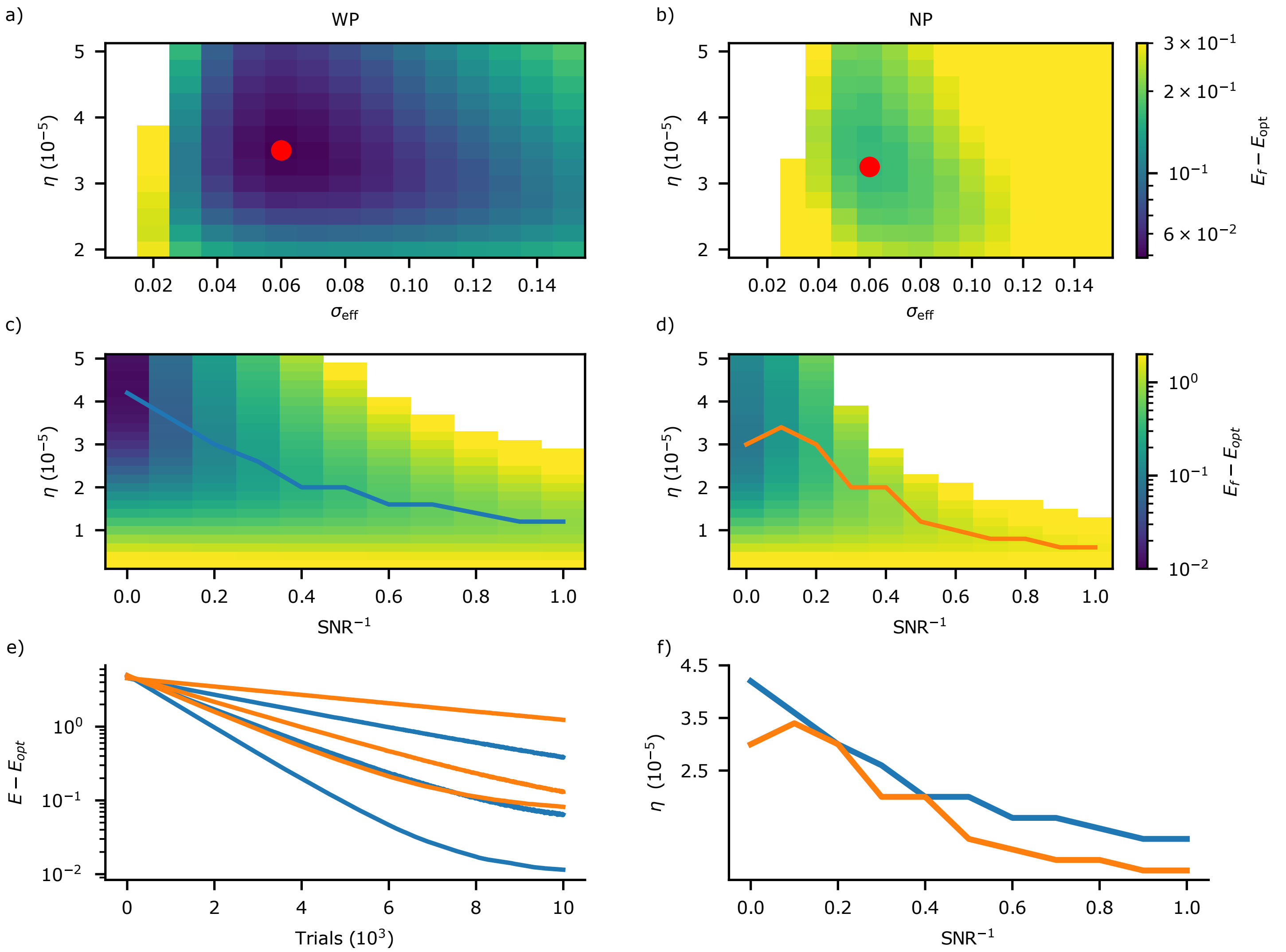


**Figure S6.** Optimal perturbation size and optimal learning rate for different levels of input noise. a,b) Excess of the final error of WP (a) and NP (b) beyond $E_{\mathrm{opt}}$, for input noise with $\mathrm{SNR}^{-1} = 0.1$ and different perturbation strengths and learning rates. The white region corresponds to parameters for which the algorithms did not converge. The red dots indicate the optimal parameter combinations. They have the same $\sigma_{\mathrm{eff}} = 0.06$ for WP and NP. WP can afford a higher learning rate and reaches a lower minimal error. Further it converges on a larger parameter region. c,d) Final error of WP (c) and NP (d) beyond $E_{\mathrm{opt}}$, for $\sigma_{\mathrm{eff}} = 0.06$ and different input noise strengths and learning rates. The optimal learning rates, which yield the lowest error, are highlighted (WP: blue connected points, NP: orange). e) Exemplary evolution (mean and SEM) of the error beyond $E_{\mathrm{opt}}$, for WP (blue) and NP (orange) and $\mathrm{SNR}^{-1} \in \{0, 0.3, 1\}$ (bottom to top curves). The learning rates are set to the optimal values. f) Joint depiction of the optimal learning rates of WP and NP, from (c) and (d). These are also the learning rates used in main text, Fig. 6. Parameters: $M = N_{\mathrm{eff}} = 10$, $N = T = 100$, $\alpha^2 = N/N_{\mathrm{eff}} = 10$, $\sigma_{\mathrm{eff}} = 0.04$, $d = 0$. The best achievable error $E_{\mathrm{opt}}$ is in presence of input noise nonzero despite $d = 0$, because the noise prevents an exact reproduction of the target. SNR is defined as the ratio of the total (summed) power in the input signal to that in the noise. Averages are taken over the last $1000$ of $10\,000$ trials and $100$ repetitions. e) shows mean and SEM.

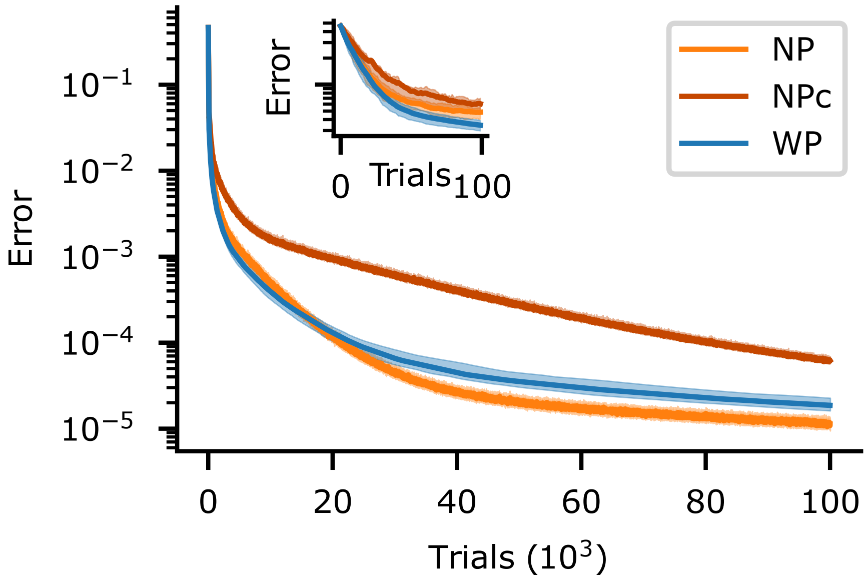


**Figure S7.** Error dynamics of the reservoir-based drawing task like Fig. 7c, with the same parameters but for infinitesimally small perturbation size $\sigma_{\mathrm{eff}}$. WP and NP perform similarly, NP slightly better. NPc performs worse, indicating that the optimal learning rate needs to be adapted when changing $T_{\mathrm{eff}}^{\mathrm{pert}}$, like in Fig. 4 (see main text “Materials and Methods”). The error curves of WP and NP differ because their error components interfere differently (SM6, Sec. “Evolution of error components related to strong and weak inputs”). Depending on the initial weight mismatch, either WP or NP can show the faster initial improvements. NP converges faster towards the end of training. Simulations suggest that the asymptotic ratio of the convergence rates of NP and WP is, however, only on the order of $1$, Fig. S9. The curves show median (solid) and interquartile range (shaded) computed over $100$ repetitions.

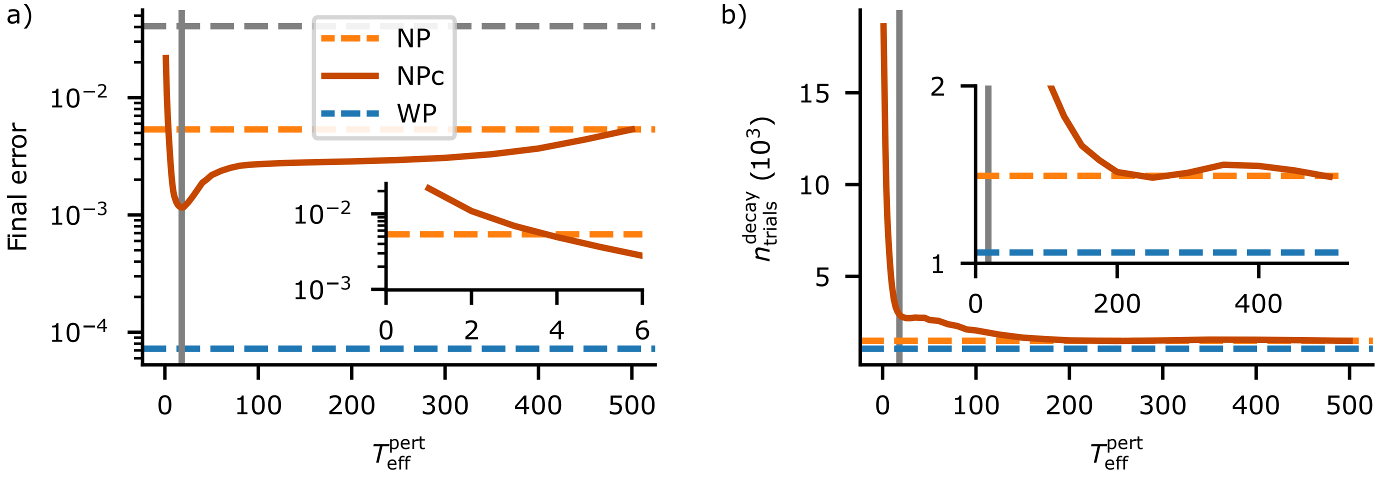


**Figure S8.** Final error and convergence time of NPc applied to the task in main text, "Reservoir computing-based drawing task", for $\eta_{NPc} = \eta^*_{NP}$ and different correlation times of the perturbations. We note that in contrast to Fig. 4 the learning rate is not adapted when changing $T_{eff}$, see main text "Materials and Methods". a) Final error of NPc (red) as a function of $T_{eff}^{pert}$. For $T_{eff}^{pert} \geq 4$ (see also inset), NPc achieves a final error equal to or lower than NP (orange dashed), but not as low as WP (blue dashed). The effective temporal perturbation dimension $T_{eff}^{pert,opt} = 18$, which minimizes the final error, is marked by a vertical line. The error achieved by a least squares fit using only the largest $5$ principle components of the reservoir is shown for comparison (gray dashed). The NPc curve shows mean and SEM of the final error over $1000$ repetitions and the last $1000$ of $30\,000$ trials. b) The number of trials needed to achieve $99\%$ of the final error reduction, $n_{trials}^{decay}$, stays largely constant for $T_{eff} > 200$ (inset) and increases slightly for $T_{eff}^{pert,opt} \leq T_{eff}^{pert} < 200$. Reducing $T_{eff}^{pert}$ below $T_{eff}^{pert,opt}$ strongly increases $n_{trials}^{decay}$. Results for NP (orange dashed) and WP (blue dashed) are shown for comparison. To determine $n_{trials}^{decay}$ we consider $10$ samples of $100$ runs each. For each sample, the mean error over runs is computed and additionally smoothed with a centered temporal running average of window size $100$. $n_{trials}^{decay}$ is then the trial at which the described average drops for the first time below $E_{f,unr} + 0.01 \cdot (E(0) - E_{f,unr})$. b) reports the mean and SEM of $n_{trials}^{decay}$ over samples.

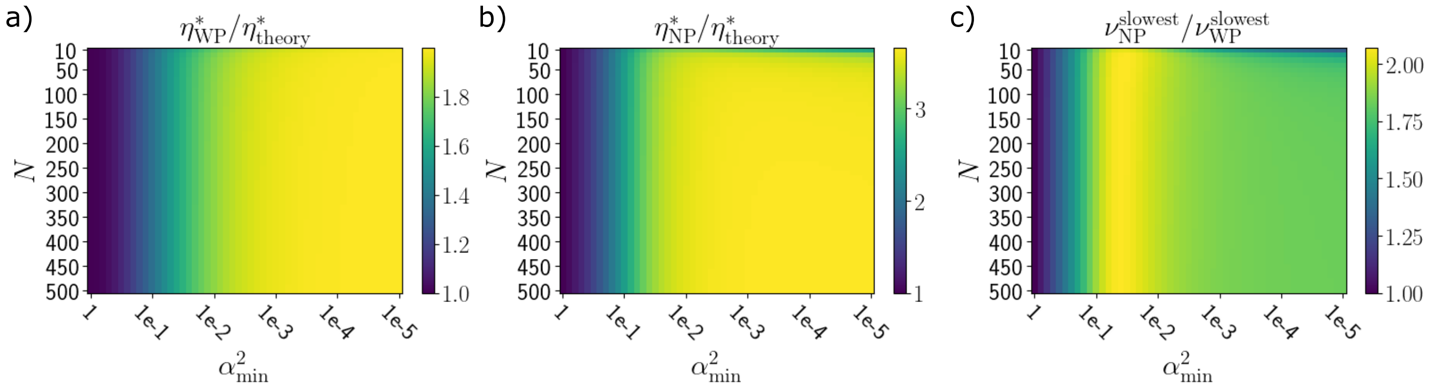


**Figure S9.** Learning rates and convergence speed of WP and NP when learning a single input-output pairing, for input strength distributions of different widths. (a,b) An analytical estimate $\eta^*_{\mathrm{theory}}$ of the optimal learning rate based on the participation ratio PR (main text, "Materials and Methods") broadly agrees with semi-analytical results $\eta^*_{\mathrm{WP\,|\,NP}}$ that maximize the convergence speed of the slowest decaying error mode (Eqs. (S173,S182)). c) The convergence rates $\nu^{\mathrm{slowest}}_{\mathrm{WP}}$ and $\nu^{\mathrm{slowest}}_{\mathrm{NP}}$ of the slowest decaying error component of WP and NP at optimal learning rates $\eta^*_{\mathrm{WP\,|\,NP}}$ stay comparable even when the input strengths differ by orders of magnitude. For each combination of the hyperparameters $N$ and $\alpha^2_{\mathrm{min}}$, $N$ orthogonal input components are constructed with exponentially decaying strengths $\alpha^2_\mu = \alpha^2_0 \cdot \gamma^\mu$ where $\gamma \leq 1$ is chosen such that $\alpha^2_0 = 1$ and $\alpha^2_N = \alpha^2_{\mathrm{min}}$. Here $\alpha^2_{\mathrm{min}} = 1$ reproduces the theory case with $\alpha^2 = 1$ and $N_{\mathrm{eff}} = N$, whereas $\alpha_{\mathrm{min}} = 1 \times 10^{-5}$ corresponds to a broad input strength distribution with $\mathrm{PR} \ll N$. $N$ and $\alpha^2_{\mathrm{min}}$ are varied on a $50 \times 51$ grid. In (a,b) we compute for each pair of hyperparameters the participation ratio PR from the distribution of input strengths and employ it to predict the optimal learning rate $\eta^*_{\mathrm{theory}} = 1/(M\,\mathrm{PR} + 2)\overline{\alpha^2}$. Here $M = 10$ and $\overline{\alpha^2} = \sum_{\mu=1}^{N} \alpha^2_\mu/\mathrm{PR}$ is the mean input strength per effective input dimension. For the comparison, we construct the matrices of error evolution $(A + B)_{\mathrm{WP|NP}}[\eta]$ (Eqs. (S177–S179)) and compute the optimal learning rate for the slowest component, $\eta^*_{\mathrm{WP|NP}}$, by numerically minimizing the largest eigenvalue of $(A + B)_{\mathrm{WP|NP}}[\eta]$. (c) displays $\nu_{\mathrm{WP|NP}} \approx 1 - a_{\mathrm{WP|NP}}$, where $a_{\mathrm{WP|NP}}$ is the largest eigenvalue of $(A + B)_{\mathrm{WP|NP}}[\eta^*]$.

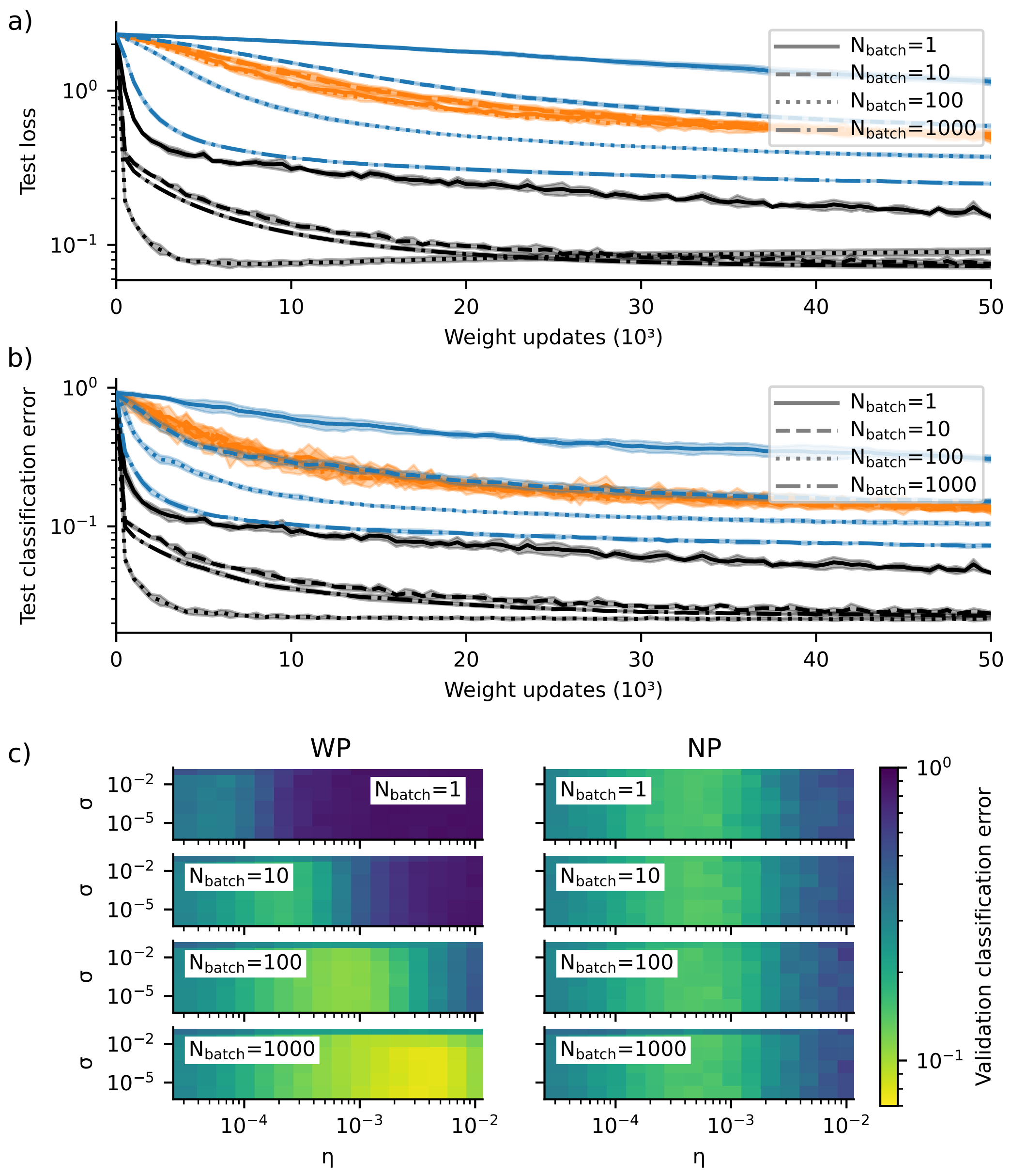


**Figure S10.** Test loss, test accuracy and grid search results for the MNIST task. a) Test loss (cross entropy loss) for the best parameters found in the grid search for WP (blue), NP (orange) and SGD (black). We note that the obtained optimal learning rate for SGD and $N_{\text{batch}} = 100$ is comparably large (Tab. 2); SGD therefore seems to overfit slightly. Lines show the mean and shaded areas show the standard deviation using 5 network instances. b) Same as a) but for the test classification error (one minus test accuracy). The grid search yields for SGD and $N_{\text{batch}} \geq 100$ similar final accuracies for very different learning rates. Therefore the best learning rates, which maximize the final accuracies, and thus also the learning curves shown here, can in this case be quite different from each other. c) Grid search to estimate the optimal learning rates for WP and NP with different perturbation strengths and batch sizes. The figure displays the mean classification error after 50000 weight updates, for 5 instances, on a validation data set not used for training.

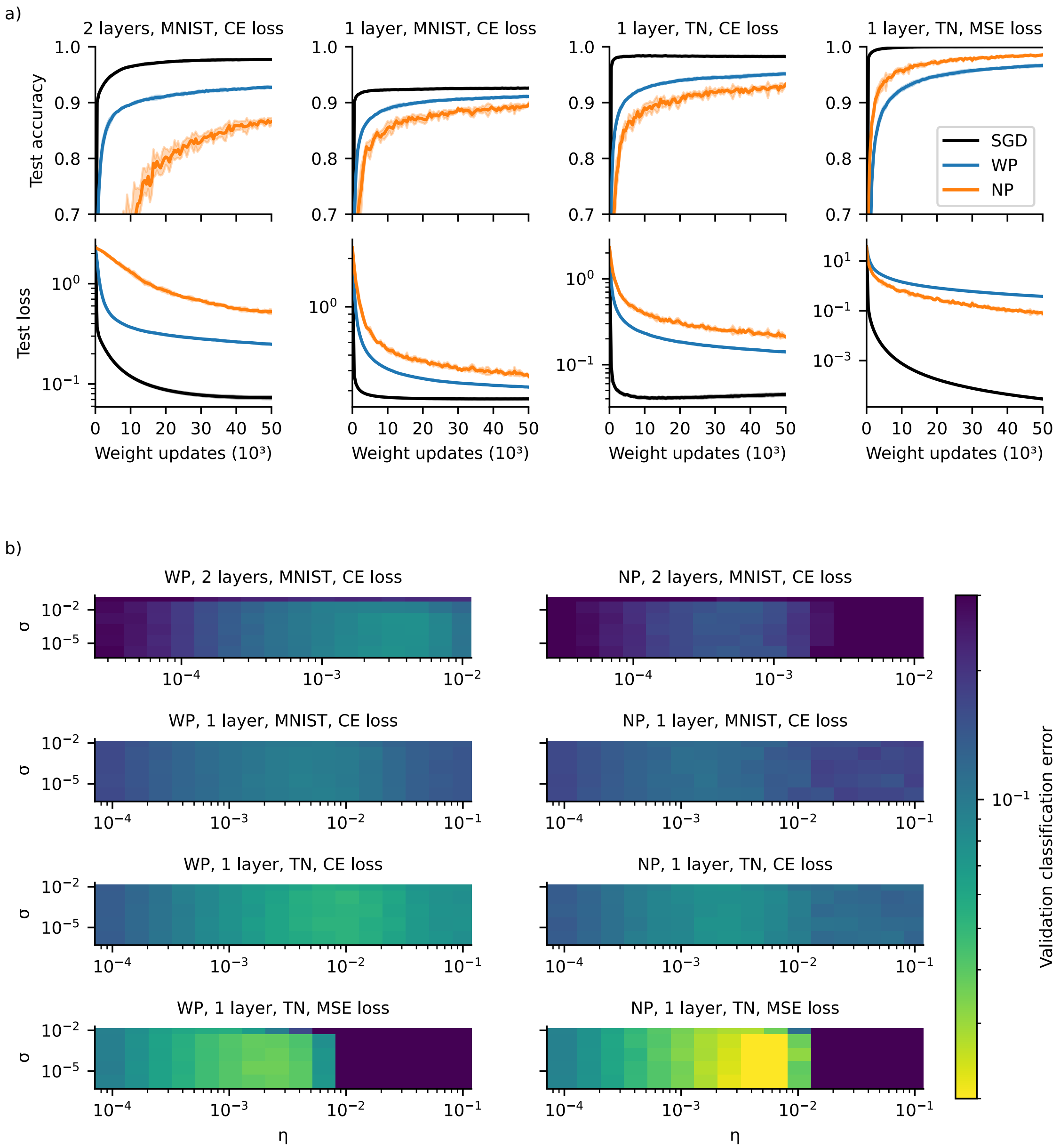


**Figure S11.** Learning performance and grid search results for four variations of the MNIST task. Specifically, the panels show learning in a two-layer network trained on MNIST using cross-entropy loss (2 layers, MNIST, CE loss; same as in Fig. 9b and Fig. S10), a single-layer network trained the same way (1 layer, MNIST, CE loss), a single-layer network with the MNIST images as input but with target labels determined by the maximal output of a teacher network that was trained on MNIST using SGD (1 layer, TN, CE loss) and a single-layer linear network with the MNIST images as input using mean-squared error loss with targets given by the raw output of the same teacher network (1 layer, TN, MSE loss). For the single-layer networks we simply remove the hidden layer from the two-layer network and in the last case (1 layer, TN, MSE loss) also remove the softmax-nonlinearity from the output layer. This yields a single layer linear network with realizable targets. We only consider $N_{\text{batch}} = 1000$ and perform a grid search to find the best performing learning parameters.
a) Test accuracy (upper row) and loss (lower row) for WP (blue), NP (orange) and SGD (black) for the best performing learning parameters. Removing the hidden layer worsens the performance of WP and SGD but improves it for NP (compare first to second column). Using a teacher network to create the target labels does not change the relative performance of WP and NP, indicating that in this task unrealizable target labels, e.g. due to bad handwriting, do not significantly harm NP (compare third to second column). Note that this does not mean that the targets are exactly realizable, because it is not possible for the network to reproduce the binary target output given by the one-hot encoded targets. Further removing all nonlinearities from the networks and using mean-squared error loss leads to better performance of NP compared to WP (compare fourth to third column). In the case of training using a teacher network, we determine the accuracy by using the index of the maximal output of the teacher network as the target label. Solid lines show the mean and shaded areas the standard deviation using 5 network instances. b) Grid search results for WP and NP as given by the mean classification error for 5 instances on a validation data set not used for training. The error is clipped at 0.3 for better visualization.

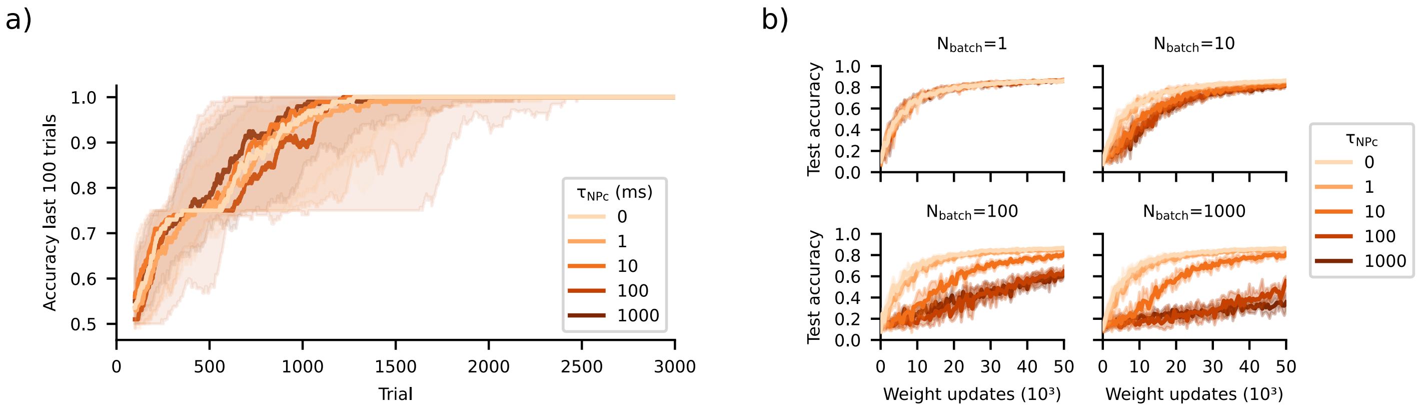


**Figure S12.** Time-correlated NP (NPc) does not improve task performance for the DNMS task and MNIST. a) Same as Fig. 8c but for NP ($\tau_{\mathrm{NPc}} = 0\,\mathrm{ms}$) and NPc with different filtering time constants ($\tau_{\mathrm{NPc}} = 1\,\mathrm{ms},\ 10\,\mathrm{ms},\ 100\,\mathrm{ms}$ and $1000\,\mathrm{ms}$). NPc does not improve task performance. b) Same as Fig. 9b but for NP ($\tau_{\mathrm{NPc}} = 0$) and NPc with different filtering time constants ($\tau_{\mathrm{NPc}} = 1,\ 10,\ 100$ and $1000$). For $N_{\mathrm{batch}} = 1$, NP and NPc are the same learning rule because trials are not temporally extended, i.e. $T = N_{\mathrm{batch}} = 1$. For larger values of $N_{\mathrm{batch}}$, NPc worsens with increasing filtering time constant. This is because the inputs in each trial are random sequences of images, whose pixel values are uncorrelated in time. Further, for a given filtering time constant $\tau_{\mathrm{NPc}} > 0$, NPc worsens with increasing batch size $N_{\mathrm{batch}}$. This may be because smaller batches are more likely to contain similar images of only a few numbers, for which learning with near-constant node perturbations still works.

| Algorithm | $N_{\text{batch}}$ | $\eta$ | Test loss | Test accuracy |
|---|---|---|---|---|
| WP | 1 | $6.81 \times 10^{-5}$ | 1.160(55) | 0.690(20) |
| | 10 | $2.15 \times 10^{-4}$ | 0.613(15) | 0.839(9) |
| | 100 | $6.81 \times 10^{-4}$ | 0.390(8) | 0.890(3) |
| | 1000 | $3.16 \times 10^{-3}$ | 0.270(7) | 0.923(2) |
| NP | 1 | $6.81 \times 10^{-4}$ | 0.515(26) | 0.856(7) |
| | 10 | $4.64 \times 10^{-4}$ | 0.541(19) | 0.860(11) |
| | 100 | $6.81 \times 10^{-4}$ | 0.510(36) | 0.860(14) |
| | 1000 | $4.64 \times 10^{-4}$ | 0.545(25) | 0.859(5) |
| SGD | 1 | 0.010 | 0.165(7) | 0.952(3) |
| | 10 | 0.056 | 0.083(6) | 0.976(1) |
| | 100 | 0.562 | 0.098(7) | 0.977(1) |
| | 1000 | 0.056 | 0.079(7) | 0.977(2) |

**Table 2.** Network performance for SGD, WP and NP on a held-out test set, after training, for the MNIST task. The third column shows the best learning rate obtained from the grid search. Values in the last two columns are the mean loss and the accuracy after 50000 weight updates, averaged over five instances (standard deviation in brackets).
